# Genome-Wide Sequencing as a First-Tier Screening Test for Short Tandem Repeat Expansions

**DOI:** 10.1101/2020.06.06.137356

**Authors:** Indhu-Shree Rajan-Babu, Junran Peng, Readman Chiu, IMAGINE Study, CAUSES Study, Arezoo Mohajeri, Egor Dolzhenko, Michael A. Eberle, Inanc Birol, Jan M. Friedman

## Abstract

Short tandem repeat (STR) expansions cause several neurological and neuromuscular disorders. Screening for STR expansions in genome-wide (exome and genome) sequencing data can enable diagnosis, optimal clinical management/treatment, and accurate genetic counselling of patients with repeat expansion disorders. We assessed the performance of lobSTR, HipSTR, RepeatSeq, ExpansionHunter, TREDPARSE, GangSTR, STRetch, and exSTRa – bioinformatics tools that have been developed to detect and/or genotype STR expansions – on experimental and simulated genome sequence data with known STR expansions aligned using two different aligners, Isaac and BWA. We then adjusted the parameter settings to optimize the sensitivity and specificity of the STR tools and fed the optimized results into a machine-learning decision tree classifier to determine the best combination of tools to detect full mutation expansions with high diagnostic sensitivity and specificity. The decision tree model supported using ExpansionHunter’s full mutation calls with those of either STRetch or exSTRa for detection of full mutations with precision, recall, and F1-score of 90%, 100%, and 95%, respectively.

We used this pipeline to screen the BWA-aligned exome or genome sequence data of 306 families of children with suspected genetic disorders for pathogenic expansions of known disease STR loci. We identified 27 samples, 17 with an apparent full-mutation expansion of the *AR*, *ATXN1*, *ATXN2*, *ATXN8*, *DMPK*, *FXN*, *HTT*, or *TBP* locus, nine with an intermediate or premutation allele in the *FMR1* locus, and one with a borderline allele in the *ATXN2* locus. We report the concordance between our bioinformatics findings and the clinical PCR results in a subset of these samples. Implementation of our bioinformatics workflow can improve the detection of disease STR expansions in exome and genome sequence diagnostics and enhance clinical outcomes for patients with repeat expansion disorders.

## INTRODUCTION

Expansions of short tandem repeats (STRs; tandemly repeated arrays of 1–6 base pair (bp) sequence motifs^1^) can cause several neurological and neuromuscular disorders^2^. Accurate genotyping (i.e., the determination of the number of copies of repeat units in an STR) is critical to the molecular diagnosis of STR expansion disorders as repeat length usually shows a positive correlation with severity and negative correlation with age of onset of clinical symptoms^3^. Repeat length also determines an STR’s allelic class (normal, NL; intermediate, IM; premutation, PM; or full-mutation, FM), which may differ with respect to associated disease phenotype^3; 4^. For example, the *FMR1* (MIM 309550) PM (55–200 CGG repeats) increases the risk for primary ovarian insufficiency (MIM 311360) and tremor/ataxia syndrome (MIM 300623). In contrast, *FMR1* FM (>200 CGG repeats) causes fragile X syndrome (MIM 300624), the most frequent Mendelian cause of intellectual disability^5^. PM and IM (also known as “mutable NL”) alleles that are meiotically unstable can expand into pathogenic FM in a single generation, while NL alleles rarely, if ever, do so^6; 7^. Expanded alleles tend to further increase in repeat length during intergenerational transmission, and, as a result, genetic anticipation (the earlier and more severe manifestation of disease symptoms with each successive generation) is common in repeat expansion disorders^8^.

Clinical laboratories typically use polymerase chain reaction (PCR) or Southern blot (SB) (alone or in combination) to characterize expansions at known disease STR loci^9^. Although highly sensitive in detecting and genotyping STR expansions, PCR and SB tests have several limitations. They are time- and labor-intensive, require extensive optimization, and do not permit concurrent analyses of more than a handful of STR loci. Next-generation sequencing (NGS), on the other hand, enables exome- and genome-wide characterization of STRs. Several algorithms have recently been developed to analyse STRs in NGS data^1; 10–14^. The incorporation of bioinformatics tools to screen for STR expansions may permit the diagnosis of repeat expansion disorders during routine diagnostic exome or genome sequencing, allow accurate genetic counseling of affected individuals and their families, and improve clinical outcomes.

The currently-available STR analysis algorithms have different attributes that determine their utility and sensitivity in detecting and characterizing repeat expansions in NGS data (Table 1). Methods like STRetch^11^ and exSTRa^12^ identify STR expansions via case-control analysis, with a caveat of either underestimating the repeat lengths of some expanded STRs^11^ or not genotyping STRs^12^. Methods that genotype STRs are known to perform better across certain repeat length ranges depending on the read type evidence considered. For instance, tools relying on reads that fully encompass an STR (“spanning reads”) to compute repeat length^15–17^ can size alleles within the length of an Illumina read (125–150 base pairs [bp]) but they perform poorly in detecting pathogenic FM expansions that exceed read length. More recent methods^1; 10; 18; 19^ that leverage on additional read types such as flanking or partially flanking reads (those that map to unique flanking sequences), in-repeat reads (IRR; those that are entirely composed of STRs with a mate that maps to the STR’s flanking sequence), and/or IRR pairs (both reads of a pair mapping to the STR) can size STRs that exceed read length. ExpansionHunter^10; 19^ and GangSTR^18^, in particular, enable the recovery of IRR and IRR pairs, which originate from an expanded STR but may incorrectly map to other STR (or “off-target”) regions with longer tracts of the same repeat motif. By allowing the inclusion of off-target sites (OTS) in analysis, ExpansionHunter and GangSTR facilitate sizing STRs that are longer than an Illumina sequencing library fragment length (350–500 bp).

**TABLE 1.** Features of some publicly available STR analysis algorithms.

| Features | lobSTR | RepeatSeq | HipSTR | TREDPARSE | ExpansionHunter | STRetch | exSTRa | GangSTR |
| --- | --- | --- | --- | --- | --- | --- | --- | --- |
| Outputs repeat length? | Y | Y | Y | Y | Y | Y |  | Y |
| Sequencing reads | Single & Paired-end | Single & Paired-end | Single & Paired-end | Paired-end | Paired-end | Paired-end | Paired-end | Paired-end |
| Sequencing platforms supported | Illumina, Sanger, 454, and IonTorrent | Illumina | Illumina | Illumina | Illumina | Illumina | Illumina | Illumina |
| Library prep. supported | PCR & PCR-free | n.a. | PCR & PCR-free | PCR & PCR-free | PCR & PCR-free | PCR & PCR-free | PCR & PCR-free | PCR & PCR-free |
| Library prep. (rcmd) | None | None | None | None | PCR-free | PCR-free | None | None |
| Aligners (rcmd) | lobSTR, BWA-MEM | Novoalign, Bowtie 2 | Indel-sensitive aligner | None | None | None | Bowtie2 | None |
| Analysis approach | Targeted & GW | Targeted & GW | Targeted & GW | Targeted | Targeted | GW | Targeted & GW | Targeted & GW |
| NGS data type supported | GS | GS | GS | GS | GS | GS & ES | GS & ES | GS & ES |
| NGS data format | .bam or .fastq/.fasta | .bam | .bam | .bam | .bam or .cram | .bam or .fastq | .bam | .bam |
| Built-In stutter correction model* | Y |  | Y | Y |  |  |  |  |
| Test of significance |  |  |  |  |  | Y | Y |  |
| Read types used | Spanning | Spanning | Spanning | Spanning, flanking or partial, paired-end reads, IRR | Spanning, flanking, IRR/IRR pairs | Anchored IRR | Flanking, anchored IRR | Spanning, flanking, IRR/IRR pairs |
| Phasing |  |  | Y |  |  |  |  |  |
| PL | C++ | C++ | C++ | Python | C++ | Java | Perl & R | C++ |
| Sizing limitation | RL | RL | RL | FL | Not limited | FL | n.a. | Not limited |
| Control dataset | Not required | Not required | Not required | Not required | Not required | Required | Not required | Not required |
| Complex repeats | n.a. | n.a. | n.a. | n.a. | Y | n.a. | n.a. | N |
| Output files | .vcf, .allelotype.stats | .repeatseq, .calls, .vcf | .vcf | .vcf, .json | .vcf, .json, .log | .tsv | p values, ECDF, tsum plots | .vcf |
| Customized regions file | Possible | Possible | Possible | Possible | Possible | Possible, but not recommended. | Possible | Possible |

\*Corrects the noise (stutters) introduced during PCR amplification-based library preparation
Library prep: library preparation protocol; rcmd: recommended; PL: programming language used
Y: Feature included; N: Feature not included
n.a.: not applicable; GW: genome-wide; GS: genome sequencing; ES: exome sequencing; IRR: in-repeat reads; RL: read-length; FL: fragment-length; Not limited: not limited by either RL or FL; ECDF: Empirical Cumulative Distribution Function; t-sum: aggregated T statistic

In terms of utility, some of these methods can analyse STRs in both exome sequencing (ES) and genome sequencing (GS) data^11; 12; 18^, while others are designed specifically for GS^1; 10; 19^. Some tools have specific NGS data requirements; for example, ExpansionHunter is designed for PCR-free GS, and exSTRa has only been extensively tested on bowtie-2^20^ alignments. Also, most methods have been recognized to perform less optimally on GC-rich STR expansions^10; 12^. These varied attributes and performance characteristics have led to the acknowledgment that a single bioinformatics tool is less likely to be able to identify pathogenic STR expansions of all repeat lengths and sequence content/composition in NGS data^12^. Recently, Tankard *et al* recommended a consensus calling approach using at least two out of four tools (TREDPARSE^1^, ExpansionHunter, STRetch, and exSTRa) to characterize expansions of known disease STRs^12^. However, it is not clear which of these (or other) STR methods alone or in combination yield optimal sensitivity and specificity.

In this study, we employed a decision tree classifier to identify the optimal tool(s) for classifying expanded FM and non-expanded alleles at known disease STR loci with high accuracy, precision, recall, and F1-score. We performed our analysis on the STR calls from nine different tools^1; 10–12; 15; 17–19; 21^ made on the GS data of patients with well-characterized STR expansions in one of eight different loci (*AR*, *ATN1*, *ATXN1*, *ATXN3*, *DMPK*, *FMR1*, *FXN*, or *HTT*)^10^ and simulated GS data harboring expansions of the GC-rich *FMR2* or *C9orf72* STR loci. These data were aligned using two different aligners, Isaac^22^, an ultra-fast aligner, and BWA-MEM^23^, recommended by the GATK best practices guidelines^24^ and widely used in GS studies^25^, to see if the choice of the aligner influences the performance of the STR methods. First, we tested the classifier on the results generated by the implementation of tools using default parameter settings. We then tweaked several parameters, such as the inclusion/exclusion of OTS and using a different FM repeat length threshold to define expansions at selected loci and implementation of exSTRa with a control cohort, to optimize the sensitivity and specificity of the STR tools included in this study. Once we established the parameters that yielded the best results, we input the data generated with these settings into the classifier and found a significant improvement in our model’s ability to detect FMs compared to our default parameter assessment. We then applied our decision tree model of STR algorithms to screen for expansions in known disease STR loci in the GS or ES data of 306 families (patient-parent trios (patient and both biological parents) or quads (patient, sibling, and both biological parents)) with a proband who is suspected to have a genetic disorder.

## METHODS AND APPROACHES

### GS Datasets with a Known Repeat Expansion

The GS datasets with a known repeat expansion analysed in this study include the BWA and Isaac alignments of: 1) the European Genome-phenome archive (EGA) dataset^10^ (EGAD00001003562), which consisted of data from 118 PCR-free GS of Coriell samples, each with an *AR*, *ATN1*, *ATXN1*, *ATXN3*, *DMPK*, *FMR1*, *FXN*, or *HTT* expansion (Supplementary Table 1a); and 2) *C9orf72* or *FMR2* expansions of varying repeat lengths simulated using the ART NGS read simulator^26^ (Supplementary Table 1b) as outlined in Supplementary Methods. The simulated GS data were included in our analysis to assess the performance of the STR algorithms on expansions of extremely high GC content (100%) that may be refractory to detection.

### Patient Cohorts and ES and GS Data Generation

The patient cohorts screened for known STR expansions in this study consist of the ES data of 146 trios or quads from the Clinical Assessment of the Utility of Sequencing and Evaluation as a Service (CAUSES) study and the GS data of 160 trios or quads from the Integrated Metabolomics And Genomics In Neurodevelopment (IMAGINE) or CAUSES studies. Subjects enrolled in the CAUSES study were children who were suspected on clinical grounds to have a single gene disorder but in whom conventional testing had not identified a genetic cause. Subjects enrolled in the IMAGINE study had impairment of motor function with onset before birth or within the first year of life and additional clinical features that made perinatal complications such as hypoxia or intracranial hemorrhage an unlikely explanation for their problems. Most of the subjects enrolled in the CAUSES or IMAGINE studies had intellectual disability. The ES or GS data from the unaffected parents were used to verify the inheritance or unstable transmission of variants. These studies were approved by the Institutional Review Board of the BC Children’s and Women’s Hospital and the University of British Columbia (H15-00092 and H16-02126).

The trio/quad ES data were sequenced by Ambry Genetics (Aliso Viejo, United States), Centogene (Rostock, Germany), or BC Cancer Agency Genome Sciences Centre (Vancouver, Canada) to a mean coverage of ~60x. The library preparation protocols and sequencers used to generate the trio/quad ES data are described in Supplementary Table 2.

The median coverage of the trio/quad GS data ranged from 36 to 80x and was generated by the McGill University and Genome Quebec Innovation Centre (Quebec, Canada). GS libraries were prepared using the NxSeq® AmpFREE Low DNA Library Kit Library Preparation Kit and Adaptors (Lucigen, Wisconsin, US) or xGen Dual Index UMI Adapters (Integrated DNA Technologies, Coralville, US) and sequenced on an Illumina HiSeqX sequencer.

The paired-end reads (125 or 150 bp) of both the ES and GS datasets were aligned to the UCSC hg19 human reference genome using BWA-MEM, and duplicates were marked with Picard^27^. All patient ES data underwent single-nucleotide variant (SNV) and indel analysis, and 145 out of the 146 trios or quads included in this study had no clinically-relevant SNV/indel variants. We also analysed the ES data of a quad with known myotonic dystrophy (Type 1; DM1 – MIM 160900) in the proband and his mother as a positive control. Our patient GS data underwent SNV, indel, structural, and mitochondrial variant analysis, with a causal variant identified in about half of the trios (unpublished data). We included the GS data of all cases in this study.

### Bioinformatics Tools for STR Analysis

The STR analysis tools implemented in this study include lobSTR^15^, HipSTR^28^, RepeatSeq^17^, TREDPARSE^1^, ExpansionHunter^10; 19^, GangSTR^18^, STRetch^11^, and exSTRa^12^. The key features of these tools and the commands and parameters used to execute them are described in Table 1 and Supplementary Table 3, respectively. We first used ExpansionHunter (EH) version 2 in this study^10^ and later included the improved iteration (version 3) of EH optimized to genotype STRs with complex or mixed repeat motifs^19^.

### Disease STR Catalogs

The STR analysis tools assess known disease STRs included within a pre-defined catalog supplied by the authors. The known pathogenic STR loci included in these catalogs, as well as their allelic categories and corresponding repeat lengths, are summarized in Supplementary Table 4. Notably, the region files for ExpansionHunter only included pre-defined OTS for *FMR1* and *C9orf72* loci, while GangSTR included OTS in the region files of all 12 pathogenic STR loci provided with the tool. Some of the region files of known disease STRs analysed in this study (*AR*, *ATN1*, *FXN*, and *FMR2*) were missing for GangSTR. Therefore, we added these loci and included their OTS as described in Mousavi *et al.* (2019)^13^.

### Interpretation of FMs and non-FMs

The data from the genotyping methods were classified as “FM” if the estimated repeat lengths of the STRs exceeded their respective FM thresholds (Supplementary Table 4). STRetch and exSTRa calls were classified as “FM” if the *p*-values post-multiple-testing-adjustment were significant (<0.05). For STRetch, we used the control file (containing data from 143 healthy individuals) provided with the tool.

### Decision Tree Classification

Decision tree analysis is a supervised machine learning (ML) classification method^29^. We employed this approach to infer the best model or the best combination of STR analysis tools to detect FM expansions with optimal sensitivity and specificity. We used the Python Scikit-Learn ML library^30^ to implement the decision tree classifier and used the STR calls from the EGA/simulated GS to train and test the classifiers on the data from the Isaac and BWA alignments.

For our preliminary decision tree analysis, we used the outputs generated using the default parameters for each of the STR analysis tools. We compiled the results generated by the STR analysis tools on the Isaac and BWA-aligned GS data. We labeled the EGA and simulated genome’s true STR expansion status or class label (FM or non-FM for a given locus). Essentially, the single known or characterized STR expansion in each of the EGA and simulated genomes was assigned to the “FM” class, while the status of the other STR loci was assigned to “non-FM”. The data from the STR callers were then transformed into binary flags: 1 indicating at least one of the two alleles was called as “FM”, and 0 indicating both alleles were “non-FM”. From there, we removed all rows with missing values and supplied the data to the classifier. We divided our dataset into 80 and 20% to train and test the classifier, respectively, and then implemented the classifier. We used the Gini index approach to ascertain the efficiency of an attribute (i.e., the STR caller) in differentiating samples belonging to the FM and non-FM classes. To evaluate the performance of the classifier, we extracted different metrics, including precision (true positives TP/(TP + false positives [FP])), recall (TP/(TP + false negatives [FN])), accuracy, and F1-score (2*((precision*recall)/(precision+recall))), and analysed the receiver operating characteristic (ROC) curve, a ratio of sensitivity (TP/(TP + FN)) and inverted specificity (1-(TN/(TN + FP))), and the precision-recall curve, a ratio of precision and recall or sensitivity. To avoid over-fitting of the data and to evaluate the robustness of the classifier, we performed 10-fold cross-validation on the training dataset and identified the best model for targeted disease STR analysis in both Isaac and BWA-aligned GS data.

We next ascertained whether tweaking some of the parameters would improve the performance of the STR analysis tools and the resultant decision tree model. First, we assessed the performance of ExpansionHunter with OTS on selected STR loci that are known to harbor expansions exceeding sequencing fragment lengths. This was to retrieve unmapped and mismapped IRR/IRR pairs and improve the repeat length estimation and detection of FMs. Second, we used a PM or IM repeat length threshold instead of FM threshold for *FMR1* and *FMR2* STR loci to classify expanded alleles and documented the sensitivity as well as the FP rates of the genotypers. Third, we tested exSTRa’s performance on BWA-aligned GS with control data from a cohort of 100 healthy individuals. We could not perform a similar analysis on Isaac-aligned GS due to the lack of Isaac-aligned GS data of healthy subjects. We carefully evaluated how these parameter tweaks influenced the performance of the STR analysis tools and selected the optimized outcomes to rerun our decision tree classifier. The precision, recall, accuracy, and F1-score metrics of this newer model generated on the test dataset and cross-validation on the training dataset were then compared to our preliminary decision tree analysis with default parameters.

### Screening for Known Disease STR Expansions in Patient Data

Finally, we screened our patient trio/quad ES and GS data for known disease STR expansions using the tools identified by the classifier. Of the probands analysed in this study, 60 have had clinical *FMR1* STR testing, three have had clinical SCA STR panel tests, one has had a clinical *FXN* STR test, and four others have had clinical *DMPK* STR tests. All of these clinical PCR-based STR tests were negative for a pathogenic expansion, except for a confirmed *DMPK* FM in a proband and his mother. All individuals who were expansion-negative at the tested locus were used as negative controls.

For all the expanded STRs identified in the patients, we analysed the parental genotype calls to verify the inheritance or unstable transmission of the alleles. Subjects with potential expansions of known disease STRs were identified for orthogonal validation to ascertain the specificity of our decision tree. Molecular testing (PCR and capillary electrophoresis) of some of the identified STR candidates was performed by Centogene (Germany).

## RESULTS

### Performance of STR Algorithms on Isaac versus BWA-aligned GS Data

The lobSTR, HipSTR, RepeatSeq, EH versions 2 and 3, GangSTR, TREDPARSE, STRetch, and exSTRa results of Isaac- and BWA-aligned EGA and simulated GS data are shown in Supplementary Tables 5 and 6, respectively. The spanning-read-only algorithms (lobSTR, HipSTR, and RepeatSeq) did not detect any FMs in either Isaac-or BWA-aligned GS data, as expected. Therefore, we omitted these tools from all subsequent analyses.

The sensitivity of EH_v2 and EH_v3, GangSTR, TREDPARSE, STRetch, and exSTRa run with default parameters in detecting FMs in Isaac- and BWA-aligned GS is summarized in Table 2. EH_v2 and EH_v3, TREDPARSE, and STRetch exhibited consistent performance and had a sensitivity of ~70% in both Isaac and BWA alignments. GangSTR’s sensitivity was better on Isaac (55%) compared to BWA (38%) alignments. In marked contrast, exSTRa detected more FMs in the BWA (88%) than Isaac (56%) alignments (see Supplementary Figures 1a and 1b for exSTRa’s plots on Isaac- and BWA-aligned GS, respectively). On Isaac-aligned data, STRetch, EH_v2, and EH_v3 detected the most FMs, followed by TREDPARSE, exSTRa, and GangSTR. On BWA-aligned data, exSTRa detected the most FMs, followed by STRetch, EH_v2, EH_v3, TREDPARSE, and GangSTR. Notably, although exSTRa and STRetch detected more FMs, they also had the most FP calls.

**TABLE 2.** Full-mutation (FM) samples detected in the Isaac- and BWA-aligned European Genome-phenome Archive (EGA) and simulated genomes by the STR tools (ExpansionHunter versions 2 and 3 (EH_v2 and EH_v3), GangSTR, TREDPARSE, STRetch, and exSTRa) implemented using default parameters. The analysed EGA and simulated dataset had 86 samples with at least one known FM allele. The number of true-positives detected by the tools, sensitivity, and the number of false-positives identified in our default analysis of the Isaac- and BWA-aligned genomes are shown.

|  | Isaac |  |  |  | BWA |  |  |  |
| --- | --- | --- | --- | --- | --- | --- | --- | --- |
|  | Detected FM Samples | True FM Samples | Sensitivity | False-Positives | Detected FM Samples | True FM Samples | Sensitivity | False-Positives |
| **EH_v2** | 65 | 86 | 0.755813953 | 6 | 64 | 86 | 0.744186047 | 6 |
| **EH_v3** | 64 | 86 | 0.744186047 | 5 | 64 | 86 | 0.744186047 | 5 |
| **GangSTR** | 47 | 86 | 0.546511628 | 8 | 33 | 86 | 0.38372093 | 8 |
| **TREDPARSE** | 62 | 86 | 0.720930233 | 3 | 62 | 86 | 0.720930233 | 10 |
| **STRetch** | 65 | 86 | 0.755813953 | 26 | 65 | 86 | 0.755813953 | 26 |
| **exSTRA** | 48 | 86 | 0.558139535 | 33 | 76 | 86 | 0.88372093 | 35 |

All FMs missed by the genotypers were under-sized and classified incorrectly as PM, IM, or NL (Supplementary Tables 7a and 7b). Additional results on the performance of the genotypers in classifying NL, IM, and PM alleles are included in Supplementary Tables 8 and 9 and Supplementary Results. Among the analysed STR loci, *FMR1*, *FMR2*, and homozygous *FXN* FMs were particularly refractory to detection (Supplementary Tables 7a and 7b).

### Decision Tree Classification

We first trained and tested the decision tree classifier on the generated default-parameter results of EH_v2, EH_v3, GangSTR, TREDPARSE, STRetch, and exSTRa. After removing the rows with missing values, the compiled STR calls of the Isaac- and BWA-aligned EGA and simulated GS datasets had 1238 and 1232 rows (one row per sample per STR locus), respectively. In Isaac-aligned data, EH_v2, which had the lowest Gini impurity or performed the best in classifying alleles was assigned to the root node (node #0) and correctly classified 47 out of 66 FMs and 918 out of 924 non-FMs in the training dataset (Supplementary Figure 2a). STRetch (node #1) and EH_v3 (node #11) detected one of the FMs missed by EH_v2. In the test dataset, the decision tree model had precision, recall, and F1-score of 100, 90, and 95%, respectively, to detect FMs; for non-FMs, the precision, recall, and F1-score were 99, 100, and 100%, respectively. The ROC and precision-recall plots are shown in Supplementary Figure 2b. The 10-fold cross-validation of this model on the training dataset yielded a ROC_AUC (Area Under the Curve) of 85.48 ± 12.58% (mean ± standard deviation).

In the BWA-aligned data, EH_v3 at the root node correctly classified 43 out of 60 FMs and 921 out of 925 non-FMs in the training dataset, with exSTRa and GangSTR recovering one of the FMs missed by EH_v3 (Supplementary Figure 3a). The precision, recall, and F1-score to detect FMs and non-FMs in the test data were 95, 81, and 88% and 98, 100, and 99%, respectively. The ROC and precision-recall curves are shown in Supplementary Figure 3b. The ROC_AUC metric of the model’s 10-fold cross-validation on the training dataset was 86.24 ± 8.38%.

In both Isaac and BWA analyses, nearly five out of the six features (STR tools) contributed to the performance of the model (Supplementary Figures 2c and 3c), led by either EH_v2 or EH_v3. The sensitivity for detecting FMs in BWA-aligned data was slightly lower compared to the Isaac analysis. Overall, the decision tree classifier on the Isaac and BWA test datasets generated using the default-parameter settings missed 10 to 20% of the FMs. To improve the detection sensitivity, we evaluated some parameters that we believed might help capture more of the true FMs.

#### TestedParameters

First we tested the effect of including OTS in the detection of FMs. While GangSTR’s region files included OTS for all analysed loci, the author-supplied JSON files of EH did not include OTS for *DMPK*, *FXN*, or *FMR2* loci, which are known to harbor expansions exceeding fragment lengths. In our initial EH run without OTS, we noted reduced sensitivity in the detection of *FXN* and *FMR2* FMs (Supplementary Table 7). Therefore, we added OTS for analysing these loci with EH_v2, which helped identify two out of three *FMR2* expansions in both Isaac- and BWA-aligned data (Supplementary Table 10). For the *FXN* locus, there was no improvement in sensitivity, highlighting the general limitation of the genotypers in reliably detecting homozygous *FXN* FM expansions. Second, because the GC-rich expansions such as those at the *FMR1* locus tend to be under-sized owing to reduced coverage even in PCR-free Illumina GS datasets^10^, we used an IM (54 repeats) and PM (60 repeats) repeat length threshold for *FMR1* and *FMR2* loci, respectively, instead of their FM threshold (both at 200 repeats). With this tweak, EH_v2 and EH_v3 detected all *FMR1* and *FMR2* FMs in Isaac-as well as BWA-aligned data (Table 3). TREDPARSE detected 83 to 89% of the *FMR1* FMs, but none of the *FMR2* FMs, while GangSTR detected 16 to 22% of the *FMR1* FMs and none of the *FMR2* FMs. The identified FPs in this analysis include the known *FMR1* PMs and a few borderline *FMR1*IM alleles that are closer to the threshold. Lastly, we hypothesized that adding data from a control cohort to exSTRa’s analysis of BWA alignments would further improve its FM detection potential. With controls, exSTRa yielded a sensitivity of 95% and detected all homozygous *FXN* FM expansions, as well as all *FMR1* and *FMR2* FMs (Supplementary Figure 1c).

**TABLE 3.**
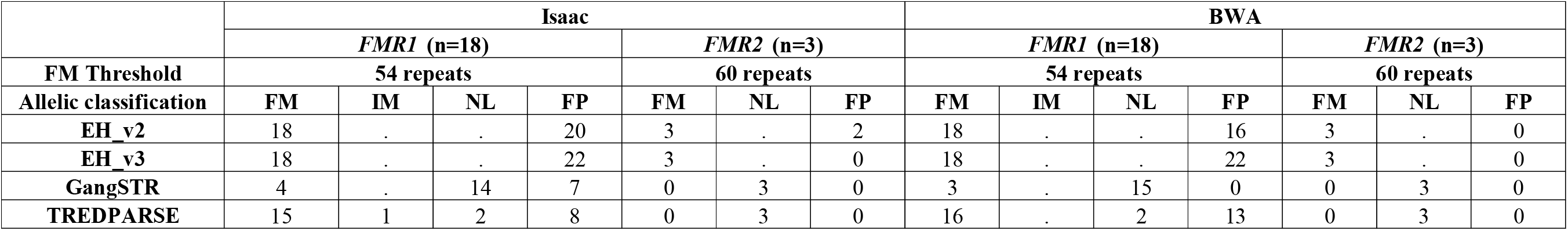
Classification of the *FMR1* and *FMR2* ExpansionHunter versions 2 and 3 (EH_v2 and EH_v3), GangSTR, and TREDPARSE genotype calls using lowered thresholds to detect FMs in the Isaac- and BWA-aligned EGA and simulated genomes of samples with known *FMR1* and *FMR2* FM expansions. The number of FMs misclassified as normal (NL) or intermediate (IM) alleles are shown. The true number (n) of known FM alleles in the *FMR1* and *FMR2* genes is indicated in parenthesis. False-positive (FP) calls made by the tools are also reported.

Of these parameters, using the IM/PM threshold for *FMR1* and *FMR2*genotype analysis and performing exSTRa’s BWA analysis with controls were useful in detecting refractory STR expansions. We fed these improved results into the classifier. In both Isaac- and BWA-aligned training datasets, EH_v2 at the root node correctly classified all but one FM and most of the non-FM alleles (Figures 1a and 2a). The classifier’s precision, recall, and F1-score in the Isaac- and BWA-aligned test datasets were 83, 100, and 91% and 90, 100, and 95% to detect FMs and 100, 98, and 99% and 100, 99, and 99% to detect non-FMs, respectively. The ROC and precision-recall plots are shown in Figures 1b and 2b. The ROC_AUC metric for cross-validation was 95.14 ± 5.12% for Isaac and 96.99 ± 3.72% for BWA. All six STR analysis tools contributed to the performance of the classifier on the improved results of Isaac-aligned GS (Figure 1c), and all but GangSTR contributed to the performance of the classifier on the BWA-aligned GS (Figure 2c). Among the STR tools, EH_v2 ranked first in both Isaac and BWA alignments. This model on the optimized results of STR algorithms performed significantly better, detecting all FMs. The decision rules that emerged from this analysis suggest the best approach to categorizing FMs is to support EH_v2 and/or EH_v3 FM calls with (at least) one other tool (STRetch, TREDPARSE, exSTRa, or GangSTR for Isaac, and STRetch or exSTRa for BWA). Unsurprisingly, we also noticed a drop in precision due to the increase in FP counts, possibly precipitated by the inaccurate identification of *FMR1* PM and some IM alleles.

**Figure 1.**
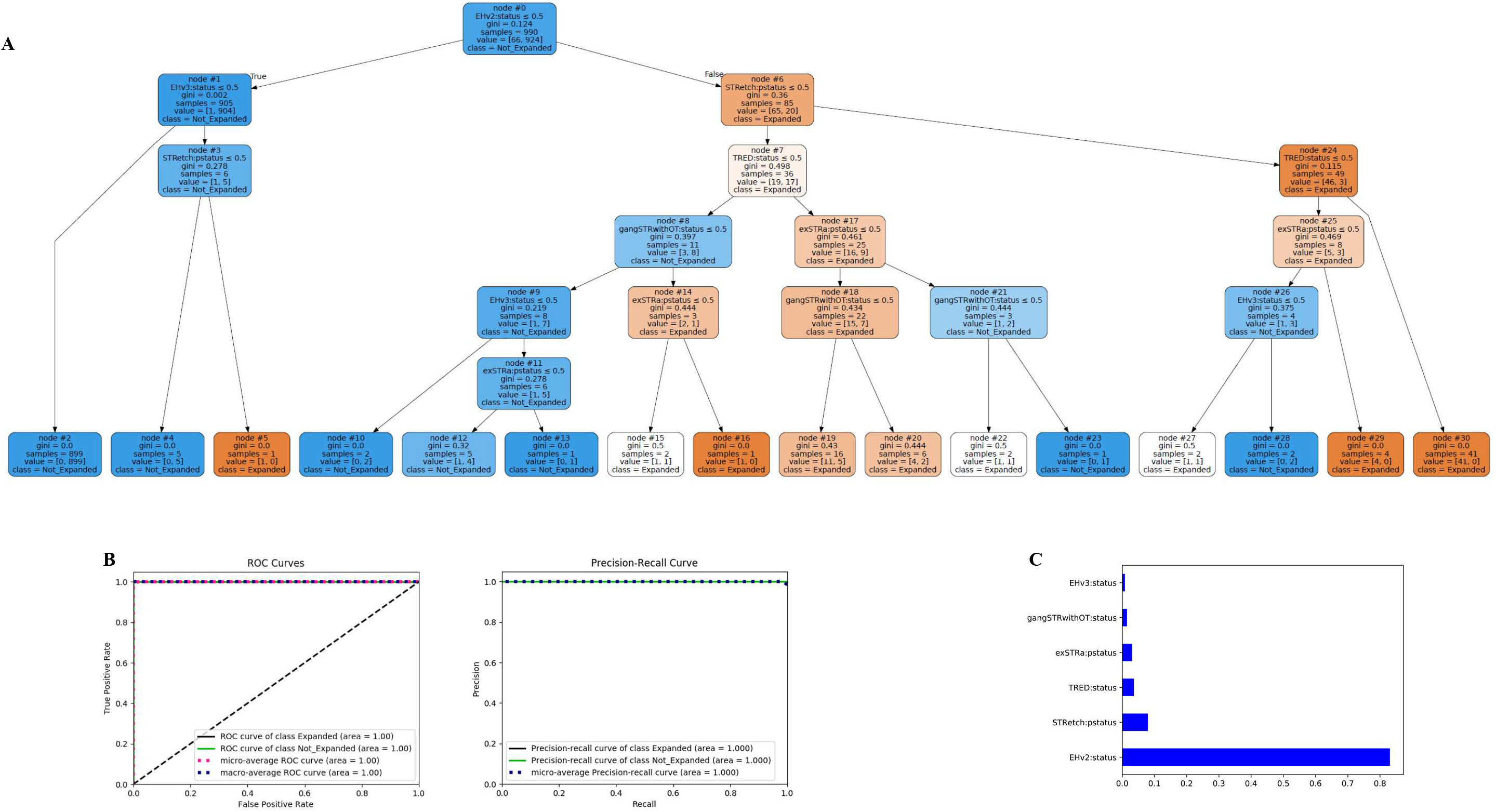
Decision tree classification of the STR calls of the Isaac-aligned EGA and simulated genome sequence (GS) data by Expansion Hunter versions 2 and 3 (EH_v2 and EH_v3), GangSTR, TREDPARSE, STRetch, and exSTRa using modified parameters. Panel (A) shows the decision tree generated by the classifier on the training dataset. Node #0 at the top of the tree is the root node. Each node lists an STR tool (feature). The “samples” number represents the total number of data points present within a particular node, and “value” shows the number of expanded (or full-mutation or FM) and non-expanded (non-FM) data points. The shade of the colour of each node reflects the proportion of expanded to non-expanded data points, with deeper blue and orange meaning more non-expanded and expanded data points, respectively. Gini index shows the impurity at each node. The terminal nodes shown in the last rows are the leaves. Leaves with a Gini of 0 have data points belonging to either the expanded or the non-expanded class. Panel B shows the ROC and precision-recall plots generated by the classifier on the test dataset. Panel C shows the ranking of the STR tools that contributed to the decision tree model.

**Figure 2.**
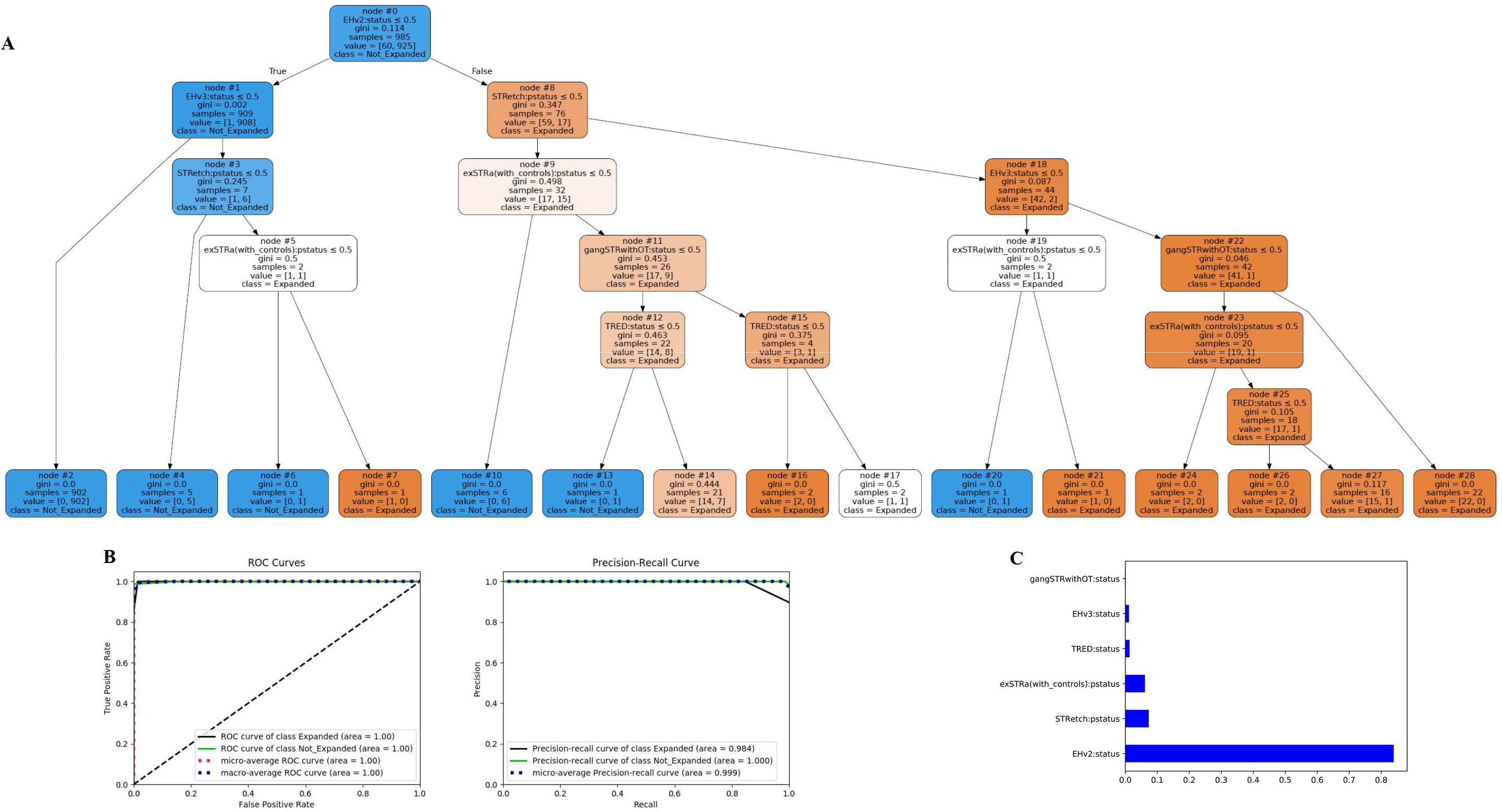
Decision tree classification of the STR calls of the BWA-aligned EGA and simulated GS data by ExpansionHunter versions 2 and 3 (EH_v2 and EH_v3), GangSTR, TREDPARSE, STRetch, and exSTRa using modified parameters. The decision tree generated by the classifier on the training dataset (A), ROC and precision-recall plots generated by the classifier on the test dataset (B) and ranking of the STR tools that contributed to the decision tree model (C) are shown.

### Analysis of Known Disease STR Loci in Clinical NGS Data

All our patient ES and GS data were BWA-aligned, so we followed the decision tree model generated on the BWA-aligned EGA and simulated GS datasets, which suggested using EH_v2 and/or EH_v3 in addition to STRetch or exSTRa. We added some additional disease STR loci to the EH_v2 variant catalog (Supplementary Table 4), analysing a total of 21 disease STRs using all four tools in our patient cohort.

First, we identified 16 EH_v2 FM expansions that were supported by at least one of EH_v3, STRetch, or exSTRa. Of the samples that were not called as expanded by EH_v2, we screened for positive calls in EH_v3, STRetch, and exSTRa outputs. STRetch and exSTRa, which had higher FP call rates in the EGA and simulated datasets, identified 298 and 442 disease STR in our patient cohort. Therefore, any positive calls made on these two tools needed to be supported by either EH_v2 or EH_v3. In total, we identified 27 samples, 17 with FM expansions of the *AR*, *ATXN1*, *ATXN2*, *ATXN8*, *DMPK*, *FXN*, *HTT*, or *TBP* locus, nine with IM or PM alleles in the *FMR1* locus, and one with a borderline allele in the *ATXN2* locus (summarized in Table 4). Supplementary Table 11 shows the EH_v2, EH_v3, STRetch, and exSTRa results of the identified STR candidates.

**TABLE 4.**
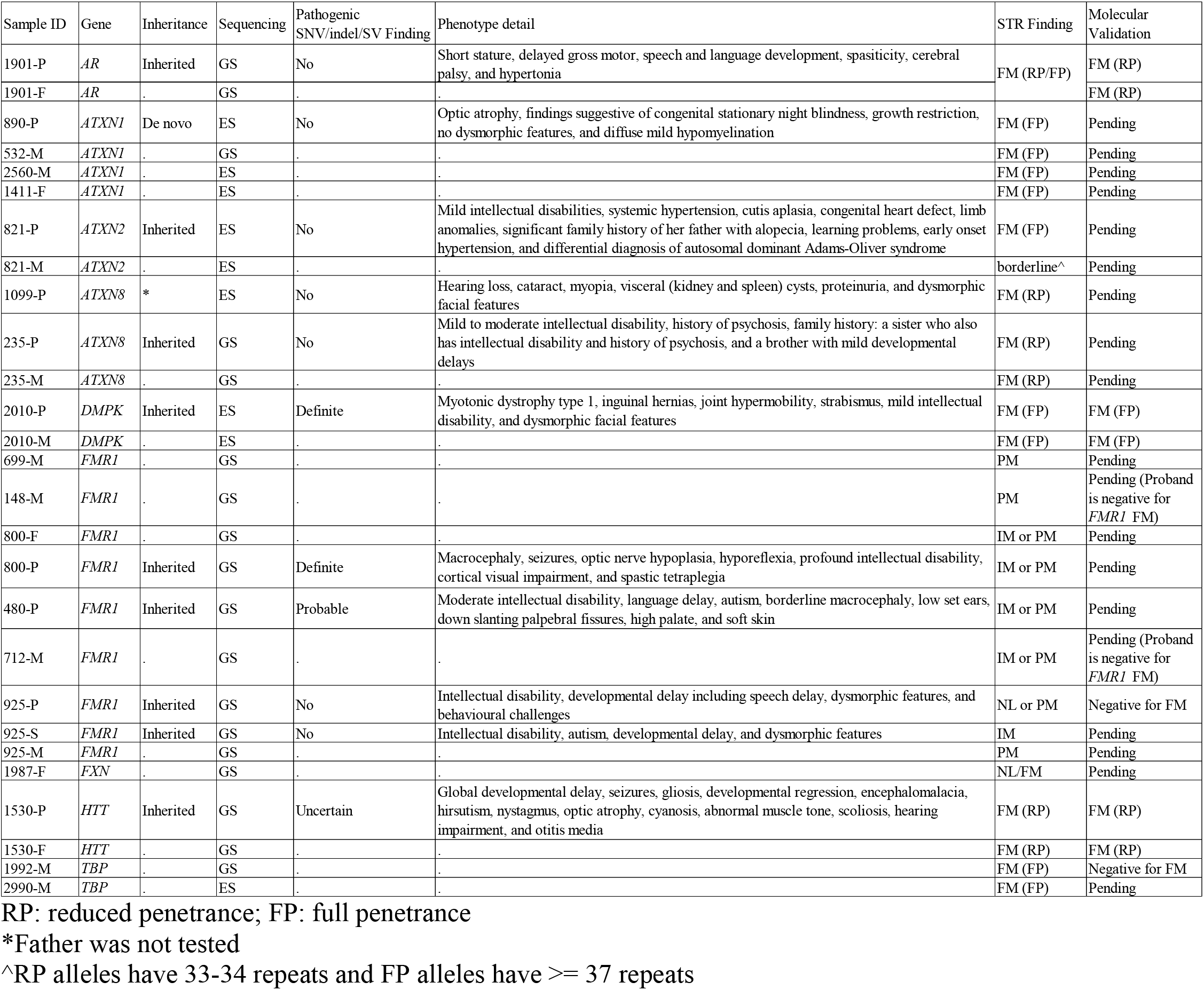
STR candidates identified in our patient cohort. Probands with an identified STR candidate are given a “-P” suffix in the “Sample ID” column, siblings, “-S”, mother, “-M”, and father, “-F”. The genes harboring the STR candidate identified by our bioinformatics workflow and the inheritance pattern deciphered by comparing the proband’s STR call with that of the parents are reported. “Sequencing” column shows the technology used: genome sequencing (GS) or exome sequencing (ES). The “Pathogenic SNV/indel/SV Finding” column indicates whether the proband has had a definite, probable, certain, or no diagnosis of a single nucleotide variant (SNV), indel, or structural variant (SV). Phenotypic presentations reported in the probands, STR Finding from our bioinformatics analysis, and the results from the molecular validation (if available) are also presented.

We found that most probands with an identified STR candidate inherited the allele from a parent, except for the *ATXN1* FM in a proband (890-P) with 39 repeats (Supplementary Table 11) compared to the parental *ATXN1* NL alleles that had 28 to 31 repeats (data not shown). The inherited expansions either remained unchanged or decreased by one or a few repeat units or increased by 1 to ~15 repeats during intergenerational transmission. We also found seven FM expansions in parents that were not inherited by the proband.

All individuals who tested negative in their molecular assessments for *FMR1*, *FXN*, *SCA*, or *DMPK* FM expansions were also categorized as non-expanded by our bioinformatics workflow (data not shown). In the ES data of the proband (2010-P) and his mother (2010-M) with DM1 and a *DMPK* FM (>50 repeats) finding on molecular assessment, EH_v2, EH_v3, and exSTRa identified the FM expansion. However, the repeat length estimated by EH_v2 and EH_v3 in 2010-P and 2010-M was ~50 repeats, which is significantly lower than the molecular findings of 150 repeats in 2010-P and 430 repeats in 2010-M (Supplementary Table 11). After including OTS to EH’s analysis of the *DMPK* locus, the FM estimate of EH_v2 and EH_v3 was ~80 repeats (data not shown).

Based on the repeat lengths estimated by EH_v2 and EH_v3, we categorized the identified FMs as reduced-or full-penetrance (Table 4; the different repeat size ranges associated with reduced- and full-penetrance of the STR expansion disorders are summarized in Supplementary Table 4). Nine of the FMs we identified in the probands and parents were in the fully-penetrant repeat size range, with another five in the reduced-penetrance range. The *AR* FM in a proband (1901-P) and her father (1901-F) was categorized as full-penetrance by EH_v3 (38 repeats) and reduced-penetrance by EH_v2 (37 repeats).

We performed PCR-based molecular tests to verify the expansion status of a subset of the identified FMs (molecular findings summarized in the last column of Table 4 and Supplementary Table 11). The *HTT* FMs identified by EH_v2 (37 repeats), EH_v3 (37 repeats), STRetch, and exSTRa in a proband (1530-P) and his father (1530-F) were concordant with the molecular test (37 ± 1 repeats). Also, the *AR* FMs in a father (1901-F) and proband (1905-P) identified by EH_v2 (37 repeats), EH_v3 (38 repeats), and STRetch were consistent with the PCR result (37 ± 1 repeats). On the other hand, the *TBP* FM in a mother (1992-M) identified by EH_v2 (52 repeats) and EH_v3 (53 repeats) could not be verified by PCR (37 ± 1 repeats). For the other identified FMs with an unknown STR expansion status, we are currently performing molecular validation.

Lastly, we investigated the genotype calls of the disease STRs made by EH_v2, EH_v3, and GangSTR in our patient ES and GS datasets to see if the NL allele frequency distribution at these loci agreed with the reported population frequencies of NL alleles (Supplementary Figures 4 and 5, and Supplementary Table 12). In general, the repeat length distribution pattern of the STR alleles for most loci was consistent across the ES (Supplementary Figure 4) and GS (Supplementary Figure 5) data, except for the *FMR1*and *FMR2* loci, which were characterized inconsistently in the ES data. EH_v3 genotyped fewer *ATXN8* alleles and also had a different repeat length distribution profile for the *ATXN7* and *HTT* loci in the ES data. For the *CSTB* locus, more 1-repeat genotype calls were made by the tools in the ES data, while we found none in the GS data. More than half of the individuals in our clinical cohort are of European ancestry, so we compared the frequency of the three most common alleles ascertained in the GS data to the common NL allele in the Caucasian population reported in the literature (Supplementary Table 12). Except for a few loci, the repeat lengths of the most common alleles determined by the tools were generally in good agreement with the reported repeat length of the common NL allele in the Caucasian population.

## DISCUSSION

The contribution of STR expansions to disease is just beginning to be understood. Hitherto, ~40 neurological disorders have been found to have a causal STR expansion mutation underlying their pathogenesis^2^, with some recent studies reporting the identification of novel pathogenic STR expansions through NGS or the more advanced third-generation long-read sequencing technologies^31–35^. The challenges in detecting and characterizing the repeat lengths of STR expansions in short-read NGS are well recognized^36^. However, recent algorithmic improvements facilitate the detection of STR expansions that exceed read and/or fragment lengths, providing us the opportunity to analyze a larger panel of known disease STR loci simultaneously through ES and GS^1; 10–14^. Some of these methods may also be useful in scanning the entire genome or exome for novel disease-causing STR expansions^11; 13^.

Of the available STR algorithms, EH, GangSTR, and TREDPARSE are particularly valuable for identifying disease-causing expansions because these programs leverage evidence beyond the reads that span an STR, enabling the genotyping of larger repeat expansions. Other methods like STRetch and exSTRa detect STR expansions but do not reliably genotype them (STRetch) or do not genotype them at all (exSTRa).

Our assessment of the performance of these STR tools on GS datasets with known repeat expansions mapped using two different aligners, Isaac and BWA, showed that the choice of aligner impacts the sensitivity of GangSTR and exSTRa. GangSTR performed better on Isaac alignments, whereas exSTRa performed better on BWA alignments.

Generally, of all the analysed disease STR loci, the detection of homozygous *FXN* FMs and the GC-rich *FMR1*and *FMR2* FMs were the most challenging. We modified some parameters to increase the FM detection potential at these loci and found that exSTRa’s sensitivity improved with control datasets, detecting all *FXN*, *FMR1*, and *FMR2* FMs in the BWA-aligned data. Also, reducing the repeat length thresholds from FM to PM/IM size ranges enabled the detection of *FMR1* and/or *FMR2* FMs with EH_v2, EH_v3, and TREDPARSE. Using this reduced cut-off also might detect some IM and PM carriers who, although not affected, may be at risk of having affected children if their IM/PM allele is highly unstable and/or susceptible to late-onset conditions^37^. Early detection and genetic counselling of these at-risk individuals might, therefore, help IM/PM allele carriers make informed reproductive decisions and avoid affected pregnancies^37^.

The ML decision tree analysis on the STR results generated using the afore-mentioned parameter modifications detected all FMs with EH_v2 and/or EH_v3 with support from one other tool (STRetch, TREDPARSE, exSTRa, or GangSTR for Isaac, and STRetch or exSTRa for BWA). EH contributed significantly to the better overall performance of the classifier on both Isaac and BWA alignments. Applying these decision rules to our clinical cohort, we identified 27 individuals with an expansion in a known disease STR locus. Of these, 17 individuals had an FM expansion of the *AR*, *ATXN1*, *ATXN2*, *ATXN8*, *DMPK*, *FXN*, *HTT*, or *TBP* locus, nine individuals had an *FMR1* allele in the IM or PM size range, and one individual had a borderline *ATXN2* allele.

Using our approach, we were able to confirm the presence of a clinically-validated *DMPK* FM in the ES data of a proband and his mother with DM1 and also confirm the inherited *HTT* and *AR* FM in two families using clinical PCR and capillary electrophoresis. We classified a *TBP* FM detected by EH_v2 and EH_v3, but unverified by PCR, as a false-positive. Importantly, none of the 68 individuals who previously had a negative clinical *FMR1*, *FXN*, *SCA*, or *HTT* test result were falsely-identified as “expanded” by our computational workflow.

For the analysis of the *DMPK* locus with EH (the default catalog file of which does not include OTS), we recommend including OTS as this could result in a significant improvement in the repeat length estimation, particularly in the GS data, and yield clinically-relevant information. Although the threshold for defining pathogenic *DMPK* FMs that cause DM1 is only 50 repeats, the different clinical forms of DM1 (mild, classic, and congenital), associated with varying severity and age of onset of symptoms, are caused by *DMPK*FMs in the range of 50-~150, ~100-~1000, and >1000 repeat units, respectively^38^. We show that with OTS, EH performs better at sizing *DMPK*FMs that ranged from ~130 to over 2000 repeats in the EGA GS data and yields estimates that correlate better with the FM repeat lengths in these individuals (Supplementary Figure 6).

Although the methods presented in this study perform well in detecting and sizing FMs, for some disease STR loci, the difference between a non-FM and an FM, or between a reduced-penetrance and full-penetrance FM is only a few repeat units, making it difficult to discriminate these borderline alleles of clinical significance. This limitation is also inherent to PCR-based tests as DNA polymerase slippage during STR amplification may result in under-or over-estimation of an STR’s size by one or two repeat units^39^.

In summary, implementation of a clinical bioinformatics workflow, such as the approach outlined in this study, to screen for STR expansions in ES and GS data can help identify disease-associated variants that would otherwise have gone undetected, promote cascade testing, and improve diagnostics and treatment/management of repeat expansion disorders.

## Supporting information

Supplementary Results

## ACKNOWLEDGMENTS

We would like to thank all the CAUSES and IMAGINE Study investigators. CAUSES Study investigators include Shelin Adam, Christele Du Souich, Alison Elliott, Anna Lehman, Jill Mwenifumbo, Tanya Nelson, Clara van Karnebeek, and Jan Friedman. The CAUSES Study is funded by Mining for Miracles, British Columbia Children’s Hospital Foundation, and Genome British Columbia. IMAGINE Study investigators include Patricia Birch, Madeline Couse, Colleen Guimond, Anna Lehman, Jill Mwenifumbo, Clara van Karnebeek, and Jan Friedman. The IMAGINE study is supported by the Canadian Institutes of Health Research (CIHR – SCA-145104) through CHILD-BRIGHT (Child Health Initiative Limiting Disability – Brain Research Improving Growth and Health Trajectories), with additional support provided by BC Children's Hospital Foundation and the Michael Smith Foundation for Health Research (MSFHR). Indhu Shree Rajan Babu is supported by the MSFHR Research Trainee Award. We thank the Rare Disease Foundation for funding our research on developing a bioinformatics pipeline to analyse STRs in next-generation sequencing data (Grant # 2332). We thank Compute Canada for the Research Allocation Competitions allocation, which facilitated our analysis of the IMAGINE and EGA genomes, and Julia Handra for coordinating the STR molecular testing of the clinical samples.

