## Supplementary Results for "Genome-Wide Sequencing as a First-Tier Screening Test for Short Tandem Repeat Expansions"

### Classification of Known NL, IM, and PM alleles

Of the known NL alleles in the EGA/simulated Isaac aligned GS (n=94), GangSTR was the only tool that detected all alleles (Supplementary Table 8a). TREDPARSE, EH_v2, and EH_v3 genotyped most of the NL alleles and overestimated the repeat lengths of one, three, and ten NL alleles, respectively, misclassifying them as an IM, PM, or FM alleles. lobSTR, HipSTR, and RepeatSeq, on the other hand, correctly genotyped 71, 30, and 54 NL alleles but were inconsistent in genotyping *C9orf72*, *FMR1*, *FMR2*, *FXN*, and/or *HTT* loci. Of the known *FMR1* IM and PM alleles (n=21), EH_v2 and EH_v3 detected 18 and 16 alleles, respectively, followed by TREDPARSE (8 alleles) and GangSTR (6 alleles) (Supplementary Table 8b). lobSTR, HipSTR, and RepeatSeq either under-sized all known IM/PM alleles or did not genotype them. We observed a similar trend among the genotyped NL and IM/PM alleles in the BWA data, except that GangSTR did not identify any of the IM/PM alleles (Supplementary Tables 9a & 9b).
