## Supplementary Tables & Figures for "Genome-Wide Sequencing as a First-Tier Screening Test for Short Tandem Repeat Expansions"

**SUPPLEMENTARY TABLE 1a:** Experimental repeat size data for the Coriell European Genome-phenome Archive dataset.

| Sample ID | Disease | Gene | Sex | Repeat Motif | Coriell Size | Genotype | Repeat Sizes Reported by Other Studies | Consensus Repeat Length |
|---|---|---|---|---|---|---|---|---|
| NA06075* | DM1 | *DMPK* | Male | $(CTG)_n$ | NL/66 | NL/FM | 12/56,70 (±0.9)[1]<br>12/55,71[2]<br>12/56,71[3] | 12/56,71 [64 (mean)] |
| NA04567 | DM1 | *DMPK* | Female | $(CTG)_n$ | NL/~700 | NL/FM | 21/637 (±33)[1]<br>21/>180[2]<br>21/>200[3] | 21/637 |
| NA05164 | DM1 | *DMPK* | Female | $(CTG)_n$ | 21/~340 | NL/FM | 21/377 (±53)[1]<br>21/>180[2]<br>21/>200[3] | 21/377 |
| NA04648 | DM1 | *DMPK* | Male | $(CTG)_n$ | NL/~1000 (minor species at ~700-800) | NL/FM | 5/1008 (±49)[1]<br>5/>180[2]<br>5/>200[3] | 5/1008 |
| NA05152 | DM1 | *DMPK* | Male | $(CTG)_n$ | NL/~1500 | NL/FM | 5/1621 (±30)[1]<br>5/>200[3] | 5/1621 |
| NA23378 | DM1 | *DMPK* | Male | $(CTG)_n$ | NL/~80-90 | NL/FM | 22/138 (±5)[1]<br>22/129-132[2]<br>22/75,126[3] | 22/138 |
| NA23374 | DM1 | *DMPK* | Female | $(CTG)_n$ | NL/~130-140 | NL/FM | NL/475[1] | NL/475 |
| NA23300* | DM1 | *DMPK* | Male | $(CTG)_n$ | NL/~150-160 | NL/FM | NL/550[1]<br>5/>180[2]<br>5/>200[3] | 5/550 |
| NA03986* | DM1 | *DMPK* | Male | $(CTG)_n$ | NL/<500 | NL/FM | 12/550*[1; 4] | 12/550 |
| NA03989 | DM1 | *DMPK* | Male | $(CTG)_n$ | NL/<2000 | NL/FM | 13/>180[2] | 13/2000 |
| NA03990 | DM1 | *DMPK* | Female | $(CTG)_n$ | NL/50-80 | NL/FM | NL/78[1] (NA03990 - LCL);<br>14/108-110[3] (NA03991- fibroblast) | 14/78 |

| | | | | | | | | |
|---|---|---|---|---|---|---|---|---|
| NA03696 | DM1 | *DMPK* | Female | $(CTG)_n$ | NL/<1000 | NL/FM | 12/697 (±13)[1]<br>12/>165,>180[2]<br>12/>200[3] | 12/697 |
| NA03759 | DM1 | *DMPK* | Male | $(CTG)_n$ | NL/<2000 | NL/FM | 14/>200[3] | 14/2000 |
| NA04034 | DM1 | *DMPK* | Male | $(CTG)_n$ | NL/<1000 | NL/FM | 12/700[1; 4] | 12/700 |
| NA03697 | DM1 | *DMPK* | Male | $(CTG)_n$ | NL/<500 | NL/FM | 12/412 (±33)[1]<br>12/>200[3] | 12/412 |
| NA03132 | DM1 | *DMPK* | Male | $(CTG)_n$ | 5/~1700 | NL/FM | 5/2078 (±217)[1]<br>5/>200[3] | 5/2078 |
| NA03756* | DM1 | *DMPK* | Male | $(CTG)_n$ | NL/<500 | NL/FM | NL/450[1]<br>13/>200[3] | 13/450 |
| NA13716 | DRPLA | *ATN1* | Male | $(CAG)_n$ | 16/68 | NL/FM | Nil | 16/68 |
| NA13717 | DRPLA | *ATN1* | Male | $(CAG)_n$ | 15/65 | NL/FM | Nil | 15/65 |
| NA03816 | FRDA | *FXN* | Female | $(GAA)_n$ | ~330/~380 | FM/FM | Nil | 330/380 |
| NA04079 | FRDA | *FXN* | Male | $(GAA)_n$ | ~420/~541 | FM/FM | Nil | 420/541 |
| NA14519 | FRDA | *FXN* | Female | $(GAA)_n$ | 9/~1335 | NL/FM | Nil | 9/1335 |
| NA15850 | FRDA | *FXN* | Male | $(GAA)_n$ | ~650/~1030 | FM/FM | Nil | 650/1030 |
| NA15847 | FRDA | *FXN* | Female | $(GAA)_n$ | NL/760 | NL/FM | Nil | NL/760 |
| NA15848 | FRDA | *FXN* | Male | $(GAA)_n$ | NL/830 | NL/FM | Nil | NL/830 |
| NA16197 | FRDA | *FXN* | Male | $(GAA)_n$ | ~760/~830 | FM/FM | Nil | 760/830 |
| NA16200 | FRDA | *FXN* | Female | $(GAA)_n$ | NL/830 | NL/FM | Nil | NL/830 |
| NA16202 | FRDA | *FXN* | Female | $(GAA)_n$ | NL/830 | NL/FM | Nil | NL/830 |
| NA16203 | FRDA | *FXN* | Female | $(GAA)_n$ | ~670/~830 | FM/FM | Nil | 670/830 |
| NA16205 | FRDA | *FXN* | Male | $(GAA)_n$ | ~530/~530 | FM/FM | Nil | 530/530 |
| NA16209 | FRDA | *FXN* | Female | $(GAA)_n$ | ~800/~800 | FM/FM | Nil | 800/800 |
| NA16210 | FRDA | *FXN* | Male | $(GAA)_n$ | ~580/~580 | FM/FM | Nil | 580/580 |
| NA16212 | FRDA | *FXN* | Female | $(GAA)_n$ | NL/500 | NL/FM | Nil | NL/500 |
| NA16216 | FRDA | *FXN* | Female | $(GAA)_n$ | ~200/~500 | FM/FM | Nil | 200/500 |
| NA16213 | FRDA | *FXN* | Male | $(GAA)_n$ | NL/420 | NL/FM | Nil | NL/420 |
| NA16215 | FRDA | *FXN* | Female | $(GAA)_n$ | NL/830 | NL/FM | Nil | NL/830 |

| | | | | | | | | |
|---|---|---|---|---|---|---|---|---|
| NA16214 | FRDA | *FXN* | Male | $(GAA)_n$ | ~600/~700 | FM/FM | Nil | 600/700 |
| NA16227 | FRDA | *FXN* | Female | $(GAA)_n$ | ~630/~830 | FM/FM | Nil | 630/830 |
| NA16229 | FRDA | *FXN* | Female | $(GAA)_n$ | NL/670 | NL/FM | Nil | NL/670 |
| NA16228 | FRDA | *FXN* | Female | $(GAA)_n$ | ~670/~830 | FM/FM | Nil | 670/830 |
| NA16237 | FRDA | *FXN* | Female | $(GAA)_n$ | NL/700 | NL/FM | Nil | NL/700 |
| NA16243 | FRDA | *FXN* | Male | $(GAA)_n$ | ~670/~1170 | FM/FM | Nil | 670/1170 |
| NA16240 | FRDA | *FXN* | Male | $(GAA)_n$ | NL/830 | NL/FM | Nil | NL/830 |
| NA16207 | FRDA | *FXN* | Female | $(GAA)_n$ | ~280/~830 | FM/FM | Nil | 280/830 |
| NA06895 | FXS | *FMR1* | Male | $(CGG)_n$ | 23 | NL | Nil | 23 |
| NA04025 | FXS | *FMR1* | Male | $(CGG)_n$ | 645 | FM | 795 (SB); >250 (PCR)[5] >200[6; 7] | 795 |
| NA04926 | FXS | *FMR1* | Male | $(CGG)_n$ | Not reported | FM | Nil | - |
| NA05131 | FXS | *FMR1* | Male | $(CGG)_n$ | Not reported | FM | Nil | - |
| NA05185 | FXS | *FMR1* | Male | $(CGG)_n$ | Not reported | FM | Nil | - |
| NA09145 | FXS | *FMR1* | Male | $(CGG)_n$ | Not reported | FM | ~660-990 (SB); >250 (PCR)[5] >200[6] | 660-990 [825 (mean)] |
| NA09237 | FXS | *FMR1* | Male | $(CGG)_n$ | 931-940 | FM | 1062 (SB); >250 (PCR)[5] >200[7] | 1062 |
| NA07063 | FXS | *FMR1* | Female | $(CGG)_n$ | Not reported | NL/FM | 32/>200[8] | 32/- |
| NA07539 | FXS | *FMR1* | Male | $(CGG)_n$ | 23 | NL | Nil | 23 |
| NA06890 | FXS | *FMR1* | Male | $(CGG)_n$ | 30 | NL | 30[6-9] | 30 |
| NA06905 | FXS | *FMR1* | Female | $(CGG)_n$ | 23/70 | NL/PM | Nil | 20/70 |
| NA07536 | FXS | *FMR1* | Male | $(CGG)_n$ | 23 | NL | Nil | 23 |
| NA07540 | FXS | *FMR1* | Female | $(CGG)_n$ | 23/29 | NL/NL | Nil | 23/29 |
| NA07542 | FXS | *FMR1* | Male | $(CGG)_n$ | 23 | NL | Nil | 23 |
| NA06910 | FXS | *FMR1* | Female | $(CGG)_n$ | 30/75-89 | NL/PM | 30/88[10] | 30/88 |

| | | | | | | | | |
|---|---|---|---|---|---|---|---|---|
| NA06894 | FXS | *FMR1* | Female | $(CGG)_n$ | 30/78 | NL/PM | Nil | 30/78 |
| NA07541 | FXS | *FMR1* | Female | $(CGG)_n$ | 29/31 | NL/NL | 29/31[6] | 29/31 |
| NA07175 | FXS | *FMR1* | Female | $(CGG)_n$ | 23/30 | NL/NL | 23/30[8; 9] | 23/30 |
| NA06889 | FXS | *FMR1* | Female | $(CGG)_n$ | 23/30 | NL/NL | Nil | 23/30 |
| NA06893 | FXS | *FMR1* | Female | $(CGG)_n$ | 23/30 | NL/NL | Nil | 23/30 |
| NA06896* | FXS | *FMR1* | Female | $(CGG)_n$ | 23/95-120-140 | NL/PM | 23/115[8; 9]<br>NL/148-201 (SB);<br>23/112,136,153,175,>250(PCR)[5]<br>23/113,133-138,155,175,198,>200[6]<br>23/183[7] | 23/No consensus |
| NA07538 | FXS | *FMR1* | Female | $(CGG)_n$ | 29/29 | NL/NL | 29/29[6-9; 11; 12] | 29/29 |
| NA07537 | FXS | *FMR1* | Female | $(CGG)_n$ | 28-29/>200 | NL/FM | 29/>200[6-9]<br>NL/329 (SB); 29/>250(PCR)[5] | 29/329 |
| NA06897 | FXS | *FMR1* | Male | $(CGG)_n$ | 477 | FM | >200[6; 7] | 477 |
| NA07174 | FXS | *FMR1* | Male | $(CGG)_n$ | 30 | NL | 30[8] | 30 |
| NA06903 | FXS | *FMR1* | Female | $(CGG)_n$ | 23/95 | NL/PM | Nil | 23/95 |
| NA07543 | FXS | *FMR1* | Female | $(CGG)_n$ | 20/29 | NL/NL | Nil | 20/29 |
| NA06852 | FXS | *FMR1* | Male | $(CGG)_n$ | >200 | FM | 395 (SB); >250 (PCR)[5]<br>>200[6-9] | 395 |
| NA06891 | FXS | *FMR1* | Male | $(CGG)_n$ | ~118 | PM | ~160[8; 9#]<br>119[6]<br>121[7] | 119 |
| NA06907 | FXS | *FMR1* | Female | $(CGG)_n$ | 29/85 | NL/PM | 29/91[8; 9] | 29/91 |
| NA06906 | FXS | *FMR1* | Male | $(CGG)_n$ | 96 | PM | 101[7] | 101 |
| NA06892 | FXS | *FMR1* | Male | $(CGG)_n$ | 93 | PM | 93[7-9]<br>~110 (SB); 90 (PCR)[5] | 93 |
| NA06904 | FXS | *FMR1* | Female | $(CGG)_n$ | 23/29 | NL/NL | Nil | 23/29 |
| NA06968 | FXS | *FMR1* | Female | $(CGG)_n$ | 32/107 | NL/PM | Nil | 32/107 |
| NA07294 | FXS | *FMR1* | Male | $(CGG)_n$ | Not reported | FM | >200[8; 9] | - |

| | | | | | | | | |
|---|---|---|---|---|---|---|---|---|
| NA09316 | FXS | *FMR1* | Male | $(CGG)_n$ | Not reported | FM | Nil | - |
| NA09317 | FXS | *FMR1* | Male | $(CGG)_n$ | Not reported | FM | Nil | - |
| NA09497 | FXS | *FMR1* | Male | $(CGG)_n$ | Not reported | FM | Nil | - |
| NA07730 | FXS | *FMR1* | Male | $(CGG)_n$ | Not reported | FM | Nil | - |
| NA03200 | FXS | *FMR1* | Male | $(CGG)_n$ | Not reported | FM | Nil | - |
| NA20235 | FXS | *FMR1* | Female | $(CGG)_n$ | 29/45 | NL/IM | 29/45[6-9; 11; 12]<br>30/45[12] | 29/45 |
| NA20238 | FXS | *FMR1* | Female | $(CGG)_n$ | 29/30 | NL/NL | 29/30[6; 8; 9; 11; 12] | 29/30 |
| NA20237 | FXS | *FMR1* | Male | $(CGG)_n$ | 100-104 | PM | 139[7]<br>100,137[12]<br>99,135[12] | 100,137 [119 (mean)] |
| NA20239 | FXS | *FMR1* | Female | $(CGG)_n$ | 20/183-193 | NL/FM | 20/~200[6-9]<br>20/No consensus[11]<br>21/200[12]<br>21/202[12] | 20/200 |
| NA20242 | FXS | *FMR1* | Female | $(CGG)_n$ | 30/73 | NL/PM | 30/74[8; 9; 12]<br>30/73,105[6]<br>30/105[7]<br>30/73[11; 12] | 30/73 |
| NA20243 | FXS | *FMR1* | Female | $(CGG)_n$ | 29/41 | NL/NL | 29/41[6-9; 11; 12] | 29/41 |
| NA20230 | FXS | *FMR1* | Male | $(CGG)_n$ | 53 | IM | 54[7-9; 12]<br>53[11] | 54 |
| NA20232 | FXS | *FMR1* | Male | $(CGG)_n$ | 46 | IM | 46[7-9; 11; 12] | 46 |

| | | | | | | | | |
|---|---|---|---|---|---|---|---|---|
| NA20233 | FXS | *FMR1* | Male | $(CGG)_n$ | 117 | PM | 120[7]<br>117[11]<br>119[12]<br>118[12] | 118 (mean) |
| NA20234 | FXS | *FMR1* | Female | $(CGG)_n$ | 31/46 | NL/IM | 31/46[8; 9; 11; 12]<br>26/46[7] | 31/46 |
| NA20236 | FXS | *FMR1* | Female | $(CGG)_n$ | 31/53 | NL/IM | 31/54[8; 9; 12]<br>31/53[11] | 31/54 |
| NA20231 | FXS | *FMR1* | Male | $(CGG)_n$ | 76 | PM | 78[7-9; 12]<br>76[11]<br>77[12] | 78 |
| NA20240 | FXS | *FMR1* | Female | $(CGG)_n$ | 30/80 | NL/PM | 30/87[8; 9]<br>30/81[6]<br>30/83[7]<br>30/80[11]<br>31/82[12]<br>31/81[12] | 30/83 (mean) |
| NA20241 | FXS | *FMR1* | Female | $(CGG)_n$ | 29/93-110 | NL/PM | NL/103-130 (SB); 29/88,111,116 (PCR)[5]<br>29/125[7]<br>29/No consensus[11]<br>30/91[12]<br>29/90[12] | 29/No consensus |
| NA20244 | FXS | *FMR1* | Male | $(CGG)_n$ | 41 | NL | 41[7-9; 11; 12] | 41 |
| NA07862 | FXS | *FMR1* | Male | $(CGG)_n$ | 501-550 | FM | >200[6-9] | 501-550 [525 (mean)] |
| NA13509 | HD | *HTT* | Female | $(CAG)_n$ | 15/70 | NL/FM | 15/72[13] | 15/72 |
| NA13515 | HD | *HTT* | Male | $(CAG)_n$ | 16/66 | NL/FM | 16/65[13] | 16/65 |
| NA13507 | HD | *HTT* | Male | $(CAG)_n$ | 15/55 | NL/FM | 15/54[13] | 15/54 |
| NA13508 | HD | *HTT* | Male | $(CAG)_n$ | 22/58 | NL/FM | 22/57[13] | 22/57 |

| | | | | | | | | |
|---|---|---|---|---|---|---|---|---|
| NA13510 | HD | *HTT* | Male | $(CAG)_n$ | 15/44 | NL/FM | 15/44[13] | 15/44 |
| NA13511 | HD | *HTT* | Male | $(CAG)_n$ | 45/47 | FM/FM | 45/47[13] | 45/47 |
| NA13512 | HD | *HTT* | Female | $(CAG)_n$ | 16/44 | NL/FM | 16/44[13] | 16/44 |
| NA13513 | HD | *HTT* | Female | $(CAG)_n$ | 15/49 | NL/FM | 15/49[13] | 15/49 |
| NA13514 | HD | *HTT* | Female | $(CAG)_n$ | 15/52 | NL/FM | 15/52[13] | 15/52 |
| NA13503 | HD | *HTT* | Female | $(CAG)_n$ | 17/45 | NL/FM | 17/45[13] | 17/45 |
| NA13504 | HD | *HTT* | Male | $(CAG)_n$ | 16/46 | NL/FM | 16/46[13] | 16/46 |
| NA13505 | HD | *HTT* | Male | $(CAG)_n$ | 22/50 | NL/FM | 22/50[13] | 22/50 |
| NA13506 | HD | *HTT* | Male | $(CAG)_n$ | 17/48 | NL/FM | 17/48[13] | 17/48 |
| NA06926 | SCA1 | *ATXN1* | Male | $(CAG)_n$ | 29/52 | NL/FM | Nil | 29/52 |
| NA13536 | SCA1 | *ATXN1* | Female | $(CAG)_n$ | 31/43 | NL/FM | Nil | 31/43 |
| NA13537 | SCA1 | *ATXN1* | Male | $(CAG)_n$ | 32/60 | NL/FM | Nil | 32/60 |
| NA06151 | SCA3 | *ATXN3* | Male | $(CAG)_n$ | 24/74 | NL/FM | Nil | 24/74 |
| NA23709 | SBMA | *AR* | Male | $(CAG)_n$ | 51 | FM | Nil | 51 |

DM1: Myotonic Dystrophy Type 1; DRPLA: Dentatorubral-pallidoluysian atrophy; FRDA: Friedreich Ataxia; FXS: Fragile X Syndrome; HD: Huntington Disease; SCA1: Spinocerebellar Ataxia Type 1; SCA3: Spinocerebellar Ataxia Type 3; SMBA: Spinal and bulbar muscular atrophy
*DMPK*: Dystrophia Myotonica Protein Kinase; *ATN1*: Atrophin 1; *FXN*: Frataxin; *FMR1*: Fragile X Mental Retardation 1; *ATXN1*: Ataxin 1; *ATXN3*: Ataxin 3; *AR*: Androgen Receptor
NL: Normal; IM: Intermediate; PM: Premutation; FM: Full-mutation
*Samples that exhibit mosaicism
SB: Southern blot analysis; LCL: Lymphoblastoid Cell Line; PCR: Polymerase Chain Reaction
#Unstable expansion of a 118-repeat PM during cell culture

**SUPPLEMENTARY TABLE 1b.** Summary of expanded STR alleles simulated using ART. Chromosome 9 open reading frame 72 (*C9orf72*) and fragile X mental retardation 2 (*FMR2*) genes were targeted for expanded allele insertions.

| Gene | Associated Health Condition | Expanded Allele Size (# of repeats) |
|---|---|---|
| *C9ORF72* | Amyotrophic lateral sclerosis (ALS) | 60, 500, 1000 |
| *FMR2* | Fragile XE syndrome (FRAXE) | 200, 500, 1000 |

**SUPPLEMENTARY TABLE 2.** ES data summary.

| **Number of exomes** | **Sequencing service provider** | **Library preparation** | **Sequencing platform** | **Read length** |
|---|---|---|---|---|
| 69 | Ambry Genetics (Aliso Viejo, United States) | xGen Exome Research Panel v1.0 with the xGen Hybridization Kit (Integrated DNA Technologies (IDT), Coralville, US) or SeqCap EZ Hybridization Kit (Roche, Basel, Switzerland) | Illumina NextSeq 500 | 2 x 150 bp |
| 47 | Centogene (Rostock, Germany) | Nextera Rapid Capture Exomes library preparation and exome enrichment kits (Illumina, San Diego, US) | Illumina HiSeq 4000 | 2 x 150 bp |
| 29 | BC Cancer Agency Genome Sciences Centre (Vancouver, Canada) | SureSelect Human All Exon V5+UTRs target enrichment technology (Agilent Technologies, Santa Clara, US) | Illumina HiSeq X | 2 x 125 bp |

**SUPPLEMENTARY TABLE 3.** Commands and parameters used for running lobSTR, RepeatSeq, HipSTR, TREDPARSE, ExpansionHunter, STRetch, exSTRa, and GangSTR.

| **STR Caller** | **Command** |
|---|---|
| lobSTR | allelotype \ <br>--command classify \ <br>--bam sample.bam \ <br>--noise_model NOISEMODELPREFIX \ <br>--out OUTPUT_PREFIX \ <br>--haploid chrX \ <br>--strinfo STRINFOFILE \ <br>--index-prefix PATH_TO_INDEX/lobSTR_ \ <br>--max-diff-ref 3000 \ <br>--filter-mapq0 \ <br>--filter-clipped \ <br>--max-repeats-in-ends 3 \ <br>--min-read-end-match 10 \ <br>--report-nocalls<br><br>(https://github.com/mgymrek/lobstr-code) |
| RepeatSeq | repeatseq -calls -repeatseq -haploid \ <br>sample.bam \ <br>hg19_reference.fasta \ <br>in.regions<br><br>(https://github.com/adaptivegenome/repeatseq) |
| HipSTR | ./HipSTR \ <br>--bams sample.bam \ <br>--fasta hg19_reference.fasta \ <br>--regions str_regions.bed<br>--str-vcf str_calls.vcf.gz \ <br>--bam-samps list_of_read_groups \ <br>--bam-libs list_of_read_groups \ <br>--log log.txt \ <br>--haploid-chrs chrX* \ <br>--min-reads 1 \ <br>--max-str-len 150 \ <br>--output-filters<br>*altered based on the sex of the analysed sample<br><br>(https://github.com/tfwillems/HipSTR) |

| STR Caller | Command |
|---|---|
| TREDPARSE | python tred.pyc CSV \ <br>--ref hg19 \ <br>--haploid chrX* \ <br>--cpus 3 \ <br>--workdir work<br><br>python tredreport.pyc work/*.json \ <br>--ref hg19 \ <br>--cpus 3 \ <br>--tsv work/Coriell.tsv<br>*altered based on the sex of the analysed sample<br><br>(https://github.com/humanlongevity/tredparse) |
| Expansion Hunter_v2 | ExpansionHunter \ <br>--bam sample.bam \ <br>--ref-fasta hg19_reference.fasta \ <br>--repeat-specs PATH_TO_JSONdirectory \ <br>--sex male* \ <br>--vcf OUTPUT_PREFIX.vcf \ <br>--json OUTPUT_PREFIX.json \ <br>--log OUTPUT_PREFIX.log<br>*altered based on the sex of the analysed sample<br><br>(https://github.com/Illumina/ExpansionHunter) |
| Expansion Hunter_v2 Exome Analysis | ExpansionHunter \ <br>--bam sample.bam \ <br>--ref-fasta hg19_reference.fasta \ <br>--repeat-specs PATH_TO_JSONdirectory \ <br>--sex male* \ <br>--read-depth <float> \ <br>--vcf OUTPUT_PREFIX.vcf \ <br>--json OUTPUT_PREFIX.json \ <br>--log OUTPUT_PREFIX.log<br>*altered based on the sex of the analysed sample<br><br>(https://github.com/Illumina/ExpansionHunter) |

| STR Caller | Command |
|---|---|
| Expansion Hunter_v3 | ExpansionHunter \ <br> --reads sample.bam \ <br> --reference hg19_reference.fasta \ <br> --variant-catalog variant_catalog.json \ <br> --output-prefix OUTPUT_PREFIX \ <br> --sex male* <br> *altered based on the sex of the analysed sample |
| Expansion Hunter_v3 Exome Analysis | ExpansionHunter \ <br> --reads sample.bam \ <br> --reference hg19_reference.fasta \ <br> --variant-catalog variant_catalog.json \ <br> --output-prefix OUTPUT_PREFIX \ <br> --sex male* <br> *altered based on the sex of the analysed sample |
| STRetch | STRetch/tools/bin/bpipe run \ <br> -p input_regions=hg19.simpleRepeat_period1-6_dedup.sorted.bed \ <br> STRetch/pipelines/STRetch_wgs_bam_pipeline.groovy \ <br> sample.bam <br><br> (https://github.com/Oshlack/STRetch) |
| STRetch Exome Pipeline | STRetch/tools/bin/bpipe run \ <br> -p input_regions=hg19.simpleRepeat_period1-6_dedup.sorted.bed \ <br> STRetch/pipelines/STRetch_exome_bam_pipeline.groovy \ <br> sample.bam <br><br> In pipeline_config.groovy file, uncomment and set: <br> EXOME_TARGET=exome_targeted_regions.bed <br><br> (https://github.com/Oshlack/STRetch) |

| STR Caller | Command |
|---|---|
| exSTRa | perl Bio-STR-exSTRa/bin/exSTRa_score.pl \ <br>hg19_reference.fasta \ <br>repeat_database.txt \ <br>sample.bam > OUTPUT_PREFIX.txt<br><br>exSTRa R package:<br>str_score <- read_score (file = file.path("/path”, “to”, “perl_output.txt” database = file.path("/path", “to”, "repeat_expansion_disorders.txt"), groups.regex = c(control = "regular expression matching control names", case = ""))<br>( str_score <- str_score[c("SBMA", "DRPLA", "SCA1", "SCA3", "DM1", "FRAXA", "FRDA","HD","HDL2","FRAXE","SCA2","SCA6","SCA7","SCA17","DM2","FTDALS1","SCA36","SCA10","EPM1A","SCA12","SCA8")] )<br>( tsum <- tsum_test(str_score, case_control = TRUE[#], trim = *) )<br>p_values(tsum, correction = c("loci"), alpha = 0.05, only.signif = TRUE)<br>*Set to 0.15 for analysis with controls and to 0.30 for analysis without controls<br>[#]Used for analysis with controls<br><br>(https://bahlolab.github.io/exSTRa/doc/exSTRa.html) |
| GangSTR | GangSTR \ <br>--bam sample.bam \ <br>--ref hg19_reference.fasta \ <br>--regions regions.bed \ <br>--out OUTPUT_PREFIX \ <br>--output-readinfo<br><br>(https://github.com/gymreklab/GangSTR) |
| GangSTR Exome Analysis | GangSTR \ <br>--bam sample.bam \ <br>--coverage <float> \ <br>--ref hg19_reference.fasta \ <br>--regions regions.bed \ <br>--out OUTPUT_PREFIX \ <br>--output-readinfo \ <br>--targeted \ <br>--nonuniform<br><br>(https://github.com/gymreklab/GangSTR) |

**SUPPLEMENTARY TABLE 4.** STR catalog. STR loci in the default variant catalogs of STRetch, GangSTR, ExpansionHunter versions 2 and 3 (EH_v2 and EH_v3), and exSTRa and our in-house disease STR variant catalog are also indicated. Allele sizes are all from GeneReviews[14], except *CBL*, which was found in Jones *et al.* (1995)[15], and *FMR2*, which was found in Youings *et al.* (2000)[16].

* *CSTB*: HipSTR, lobSTR, and RepeatSeq default variant catalogs have indicated the repeat motif as CGGGG

** Four genes included in this table contain STR expansions with compound motifs. Therefore, the default variant catalogs of certain tools may indicate motifs different than in the "Repeat Motif" column. *RAPGEF2*: STRetch, RepeatSeq, HipSTR, and lobSTR default variant catalogs indicate TTTTA as the repeat motif. *RFC1*: STRetch, GangSTR, RepeatSeq, HipSTR, and lobSTR default variant catalogs indicate wild-type repeat motif AAAAG instead of pathogenic repeat motif AAGGG. *SAMD12*: STRetch, RepeatSeq, HipSTR, and LobSTR default variant catalogs indicate AAATA as the repeat motif. *TNRC6A*: STRetch, RepeatSeq, HipSTR, and lobSTR default variant catalog indicates AAAAT as the repeat motif.

| Gene | Disorder | Chrom | Start | End | Repeat Motif | Normal Allele | Intermediate/ Premutation Allele | Reduced Penetrance Allele | Full Penetrance Allele | STRetch | GangSTR | EH_v2 | EH_v3 | exSTRa | RepeatSeq | HipSTR | lobSTR | TREDPARSE | Used in study (EH_v2) | Common to EH_v2 and EH_v3, STRetch, and exSTRa | Pathogenic Threshold Used |
|---|---|---|---|---|---|---|---|---|---|---|---|---|---|---|---|---|---|---|---|---|---|
| ***AR*** | Spinal and bulbar muscular atrophy | X | 66765159 | 66765225 | GCA | ≤34 | | 36-37 | ≥38 | ✓ | | ✓ | ✓ | ✓ | ✓ | ✓ | ✓ | ✓ | ✓ | ✓ | 37 |
| ***ARX*** | Early-infantile epileptic encephalopathy; Partington syndrome | X | 25031779 | 25031808 | GCG | ≤16 | | | 17-27 | ✓ | ✓ | | | | ✓ | ✓ | ✓ | ✓ | ✓ | | |
| ***ATN1*** | Dentatorubral-Pallidoluysian atrophy | 12 | 7045891 | 7045936 | CAG | 6-35 | 20-35 | | ≥48 | ✓ | | ✓ | ✓ | ✓ | ✓ | ✓ | ✓ | ✓ | ✓ | ✓ | 47 |
| ***ATXN1*** | Spinocerebellar ataxia type 1 | 6 | 16327866 | 16327953 | CTG | 6-35<br>36-44 [with CAT interruptions] | 36-38 [without CAT interruptions] | 44 [with CAT interruptions] | ≥39 | ✓ | ✓ | ✓ | ✓ | ✓ | ✓ | ✓ | ✓ | ✓ | ✓ | ✓ | 38 |
| ***ATXN2*** | Spinocerebellar ataxia type 2 | 12 | 112036754 | 112036823 | CTG | ≤31 | | 33-34 [dominant alleles] | 31/31 [recessive alleles]; ≥37 [dominant alleles] | ✓ | ✓ | | ✓ | ✓ | ✓ | ✓ | ✓ | ✓ | ✓ | ✓ | 36 |
| ***ATXN3*** | Spinocerebellar ataxia type 3 | 14 | 92537354 | 92537378 | CTG | 12-44 | 45-59 | | 60-87 | ✓ | ✓ | ✓ | ✓ | ✓ | ✓ | ✓ | ✓ | ✓ | ✓ | ✓ | 59 |
| ***ATXN7*** | Spinocerebellar ataxia type 7 | 3 | 63898361 | 63898391 | CAG | ≤19 | 28-33 | 34-36 | ≥36 | ✓ | ✓ | | ✓ | ✓ | ✓ | ✓ | ✓ | ✓ | ✓ | ✓ | 35 |
| ***ATXN8*** | | | | | CAG | ~80 | | | Unknown | | | | | | | | | | | ✓ | |
| ***ATXN8OS*** | Spinocerebellar ataxia type 8 | 13 | 70713515 | 70713560 | CTG | 15-50 (CTA·TAG)n(CTG·CAG)n repeats | | Reduced penetrance: (CTA·TAG)n(CTG·CAG)n repeats of all sizes;<br>Higher penetrance: 80-250 (CTA·TAG)n(CTG·CAG)n repeats | NA (penetrance is <100%) | ✓ | ✓ | | ✓ | ✓ | ✓ | ✓ | ✓ | ✓ | ✓ | | 75 |
| ***ATXN10*** | Spinocerebellar ataxia type 10 | 22 | 46191234 | 46191304 | ATTCT | 10-32 | | 33-850 | 800-4500 | ✓ | ✓ | | ✓ | ✓ | | ✓ | ✓ | ✓ | ✓ | ✓ | 33 |
| ***C9orf72*** | C9orf72-related amyotrophic lateral sclerosis and frontotemporal dementia | 9 | 27573526 | 27573544 | GGCCCC | <25 | | | >60 | ✓ | ✓ | ✓ | ✓ | ✓ | ✓ | ✓ | ✓ | ✓ | ✓ | ✓ | 60 |
| ***CACNA1A*** | Spinocerebellar ataxia type 6 | 19 | 13318672 | 13318711 | CTG | ≤18 | | | 20-33 | ✓ | ✓ | | ✓ | ✓ | ✓ | ✓ | ✓ | ✓ | ✓ | ✓ | 19 |
| ***CBL*** | FRA11B (Jacobsen syndrome) | 11 | 119076999 | 119077032 | CGG | 11 | 80-100 | | >100 | ✓ | ✓ | | ✓ | | ✓ | ✓ | ✓ | | ✓ | | |
| ***CNBP*** | Myotonic dystrophy type 2 | 3 | 128891419 | 128891499 | CAGG | ≤26 | | | >75 | ✓ | ✓ | | ✓ | ✓ | ✓ | | ✓ | ✓ | ✓ | ✓ | 75 |
| ***CSTB **** | Unverricht-Lundborg disease | 21 | 45196324 | 45196360 | CGCGGGGC GGGG | 2-3 | | | ≥30 | | ✓ | | ✓ | ✓ | ✓ | ✓ | ✓ | ✓ | ✓ | | |
| ***DIP2B*** | Mental retardation, FRA12A type | 12 | 50898784 | 50898805 | GGC | 6-23 | | | >350 | | ✓ | | ✓ | | ✓ | ✓ | ✓ | | | | |
| ***DMPK*** | Myotonic dystrophy type 1 | 19 | 46273462 | 46273522 | CAG | 5-34 | 35-49 | | >50 | ✓ | ✓ | ✓ | ✓ | ✓ | ✓ | ✓ | ✓ | ✓ | ✓ | ✓ | 50 |
| ***FMR1*** | FMR1-related disorders | X | 146993568 | 146993628 | CGG | 5-44 | 45-200 | | >200 | ✓ | ✓ | ✓ | ✓ | ✓ | ✓ | ✓ | ✓ | ✓ | ✓ | ✓ | 54 |
| ***FMR2*** | Fragile X syndrome, FRAXE type | X | 147582157 | 147582202 | GCC | 11-30 | 31-200 | | >200 | ✓ | | | ✓ | ✓ | ✓ | ✓ | ✓ | ✓ | ✓ | ✓ | 60 |
| ***FOXL2*** | Blepharophimosis, ptosis, and epicanthus inversus | 3 | 138664862 | 138664904 | NGC | 14 | | | 15-24 | | | | | | ✓ | | | ✓ | ✓ | | |
| ***FXN*** | Friedreich ataxia | 9 | 71652202 | 71652220 | GAA | 5-33 | 34-65 | | ≥66 | ✓ | | ✓ | ✓ | ✓ | ✓ | | ✓ | ✓ | ✓ | ✓ | 65 |
| ***HOXA13*** | Hand-foot-genital syndrome | 7 | 27239444 | 27239480 | NGC | ≤18 | | | >18 | | | | | | ✓ | ✓ | ✓ | ✓ | ✓ | | |
| ***HOXD13*** | Syndactyly type V | 2 | 176957786 | 176957831 | GCN | 15 | | | ≥22 | ✓ | | | | | ✓ | ✓ | ✓ | ✓ | ✓ | | |
| ***HTT*** | Huntington disease | 4 | 3076603 | 3076660 | CAG | ≤26 | 27-35 | 36-39 | ≥40 | ✓ | ✓ | ✓ | ✓ | ✓ | ✓ | ✓ | ✓ | ✓ | ✓ | ✓ | 39 |
| ***JPH3*** | Huntington disease-like 2 | 16 | 87637895 | 87637934 | GCT | 6-28 | 33-35 | | ≥40 | ✓ | ✓ | | ✓ | ✓ | ✓ | ✓ | ✓ | ✓ | ✓ | ✓ | 39 |
| ***LRP12*** | Oculopharyngodistal myopathy | 8 | 105601200 | 105601227 | CCG | 13-45 | | | Unknown | ✓ | | | | | ✓ | ✓ | ✓ | | | | |
| ***NOP56*** | Spinocerebellar ataxia type 36 | 20 | 2633379 | 2633403 | GGCCTG | 3-14 | | | ≥650 | ✓ | ✓ | | ✓ | ✓ | ✓ | ✓ | ✓ | ✓ | ✓ | ✓ | 14 |
| ***NOTCH2NLC*** | Neuronal intranuclear inclusion disease | 1 | 145209323 | 145209344 | GGC | <38 | | | ≥66 | | | | | | | | | | | | |
| ***PABPN1*** | Oculopharyngeal muscular dystrophy | 14 | 23790681 | 23790711 | GCN | 10 | | | 11/11 [recessive alleles]; 12-17 [dominant alleles] | | | | | | ✓ | ✓ | ✓ | ✓ | ✓ | | |
| ***PHOX2B*** | Congenital central hypoventilation syndrome | 4 | 41747988 | 41748048 | NGC | ≤20 | | 24-25 | 26-33 | ✓ | | | ✓ | | ✓ | ✓ | ✓ | ✓ | ✓ | | |
| ***PPP2R2B*** | Spinocerebellar ataxia type 12 | 5 | 146258290 | 146258320 | GCT | 7-31 | | | 51-78 | ✓ | ✓ | | ✓ | ✓ | ✓ | ✓ | ✓ | ✓ | ✓ | ✓ | 50 |
| ***RAPGEF2 ***** | Familial adult myoclonic epilepsy type 7 | 4 | 160263678 | 160263768 | TTTCA | 0 | | | Unknown | ✓ | | | | ✓ | ✓ | ✓ | ✓ | | | | |
| ***RFC1 ***** | Cerebellar ataxia, neuropathy, and vestibular areflexia syndrome | 4 | 39350044 | 39350099 | AAGGG | AAAAG(11) | | | ≥400 | ✓ | ✓ | | | ✓ | ✓ | ✓ | ✓ | | | | |
| ***RUNX2*** | Cleidocranial dysplasia spectrum disorder | 6 | 45390487 | 45390538 | GCN | 17 | | | 20-27 | ✓ | ✓ | | | | ✓ | | ✓ | ✓ | ✓ | ✓ | |
| ***SAMD12 ***** | Familial adult myoclonic epilepsy type 1 | 8 | 119379054 | 119379157 | TGAAA | 0 | | | ≥105 | ✓ | | | | ✓ | ✓ | ✓ | ✓ | | | | |
| ***SOX3*** | Panhypopituitarism and intellectual disability with growth hormone deficiency | X | 139586481 | 139586526 | NGC | 15 | | | 22-26 | | | | | | ✓ | ✓ | ✓ | ✓ | ✓ | | |
| ***TBP*** | Spinocerebellar ataxia type 17 | 6 | 170870995 | 170871109 | CAN | 25-40 | | 41-48 | ≥49 | ✓ | ✓ | | ✓ | ✓ | ✓ | ✓ | ✓ | ✓ | ✓ | ✓ | 48 |
| ***TCF4*** | Fuchs endothelial corneal dystrophy | 18 | 53253386 | 53253458 | CAG | <40 | | >80 | NA (penetrance is <100%) | ✓ | | | ✓ | ✓ | ✓ | ✓ | ✓ | | | | |
| ***TNRC6A ***** | Familial adult myoclonic epilepsy type 6 | 16 | 24624760 | 24624850 | TTTCA | 0 | | | 29 | ✓ | | | | ✓ | ✓ | ✓ | ✓ | | | | |
| ***XYLT1*** | Baratela-Scott syndrome (Desbuquois dysplasia type 2) | 16 | 17564764 | 17564779 | GCC | 9-20 | | | ~>72 | | ✓ | | | | ✓ | ✓ | ✓ | | | | |
| ***ZIC2*** | Holoprosencephaly type 5 | 13 | 100637702 | 100637747 | GCN | 15 | | | 25 | ✓ | | | | | ✓ | ✓ | ✓ | ✓ | ✓ | | |

**SUPPLEMENTARY TABLE 5.** Allelic calls from ExpansionHunter versions 2 and 3 (EH_v2 and EH_v3), GangSTR, TREDPARSE (TRED), lobSTR, HipSTR, RepeatSeq, STRetch, and exSTRa STR detection tools for the EGA and simulated GS data (with Isaac alignment). For each tool, a1 and a2 indicate the calls for allele 1 and allele 2, respectively, while CI indicates the confidence interval. For status, "EXPANDED" indicates at least one allele was reported to be over the full-mutation threshold, while "normal" indicates that both alleles were reported to be under the threshold. For STRetch, both the p-value and genotyped repeat counts (repeatUnit) are reported. The p-status for STRetch and exSTRa are marked as "EXPANDED" for those samples with p-values <0.05, and "normal" for those samples with p-values >0.05. For all tools except STRetch and exSTRa, empty cells indicate a call was not made for this sample in the corresponding STR locus.

| sampleID:gene | EH_v2:a1 | EH_v2:a2 | EH_v2:CI-a1 | EH_v2:CI-a2 | EH_v2:status | EH_v3:a1 | EH_v3:a2 | EH_v3:CI-a1 | EH_v3:CI-a2 | EH_v3:status | GangSTR:a1 | GangSTR:a2 | GangSTR:CI-a1 | GangSTR:CI-a2 | GangSTR:status | TRED:a1 |
|---|---|---|---|---|---|---|---|---|---|---|---|---|---|---|---|---|
| NA06075:DMPK | 12 | 71 | 12-12 | 55-84 | EXPANDED | 12 | 64 | 12-12 | 50-75 | EXPANDED | 12 | 62 | 12-12 | 53-74 | EXPANDED | 12 |
| NA04567:DMPK | 21 | 149 | 21-21 | 122-175 | EXPANDED | 21 | 137 | 21-21 | 110-162 | EXPANDED | 21 | 137 | 21-21 | 112-162 | EXPANDED | 21 |
| NA05164:DMPK | 21 | 121 | 21-21 | 96-143 | EXPANDED | 21 | 121 | 21-21 | 95-144 | EXPANDED | 21 | 152 | 21-21 | 130-173 | EXPANDED | 21 |
| NA04648:DMPK | 5 | 159 | 5-5 | 130-185 | EXPANDED | 5 | 135 | 5-5 | 109-159 | EXPANDED | 5 | 140 | 5-5 | 123-157 | EXPANDED | 5 |
| NA05152:DMPK | 5 | 148 | 5-5 | 122-172 | EXPANDED | 5 | 119 | 5-5 | 97-140 | EXPANDED | 5 | 235 | 5-5 | 214-256 | EXPANDED | 5 |
| NA23378:DMPK | 22 | 126 | 22-22 | 103-147 | EXPANDED | 22 | 132 | 22-22 | 103-159 | EXPANDED | 22 | 105 | 22-24 | 84-123 | EXPANDED | 22 |
| NA23374:DMPK | 5 | 138 | 5-5 | 111-163 | EXPANDED | 5 | 138 | 5-5 | 110-166 | EXPANDED | 5 | 119 | 5-5 | 102-134 | EXPANDED | 5 |
| NA23300:DMPK | 5 | 130 | 5-5 | 106-154 | EXPANDED | 5 | 135 | 5-5 | 107-162 | EXPANDED | 5 | 25 | 5-5 | 16-28 | normal | 5 |
| NA03986:DMPK | 12 | 116 | 12-12 | 96-137 | EXPANDED | 12 | 116 | 12-12 | 92-137 | EXPANDED | 12 | 103 | 12-12 | 86-121 | EXPANDED | 12 |
| NA03989:DMPK | 13 | 154 | 13-13 | 126-180 | EXPANDED | 13 | 133 | 13-13 | 107-158 | EXPANDED | 13 | 186 | 13-13 | 165-210 | EXPANDED | 13 |
| NA03990:DMPK | 14 | 100 | 14-14 | 80-117 | EXPANDED | 14 | 94 | 14-14 | 74-112 | EXPANDED | 14 | 72 | 14-14 | 61-86 | EXPANDED | 14 |
| NA03696:DMPK | 12 | 138 | 12-12 | 112-161 | EXPANDED | 12 | 136 | 12-12 | 108-161 | EXPANDED | 12 | 125 | 12-20 | 105-146 | EXPANDED | 12 |
| NA03759:DMPK | 14 | 117 | 14-14 | 94-138 | EXPANDED | 14 | 123 | 14-14 | 97-148 | EXPANDED | 14 | 157 | 14-20 | 142-178 | EXPANDED | 14 |
| NA04034:DMPK | 12 | 117 | 12-12 | 95-139 | EXPANDED | 12 | 117 | 12-12 | 90-142 | EXPANDED | 12 | 102 | 12-12 | 85-118 | EXPANDED | 12 |
| NA03697:DMPK | 12 | 137 | 12-12 | 112-162 | EXPANDED | 12 | 134 | 12-12 | 108-161 | EXPANDED | 12 | 122 | 12-12 | 102-140 | EXPANDED | 12 |
| NA03132:DMPK | 5 | 113 | 5-5 | 91-133 | EXPANDED | 5 | 108 | 5-5 | 86-129 | EXPANDED | 5 | 149 | 5-5 | 131-172 | EXPANDED | 5 |
| NA03756:DMPK | 13 | 141 | 13-13 | 116-166 | EXPANDED | 13 | 131 | 13-13 | 105-156 | EXPANDED | 13 | 129 | 13-13 | 108-148 | EXPANDED | 13 |
| NA13716:ATN1 | 20 | 60 | 20-20 | 52-69 | EXPANDED | 20 | 59 | 20-20 | 50-68 | EXPANDED | 16 | 55 | 16-16 | 41-62 | EXPANDED | 16 |
| NA13717:ATN1 | 19 | 64 | 19-19 | 54-72 | EXPANDED | 19 | 58 | 19-19 | 50-65 | EXPANDED | 15 | 56 | 15-15 | 40-65 | EXPANDED | 15 |
| NA03816:FXN | 90 | 133 | 50-107 | 73-157 | EXPANDED | 83 | 125 | 50-116 | 74-176 | EXPANDED | 6 | 187 | 6-6 | 173-196 | EXPANDED | 51 |
| NA04079:FXN | 99 | 148 | 50-115 | 81-173 | EXPANDED | 94 | 152 | 50-139 | 86-218 | EXPANDED | 6 | 40 | 6-6 | 16-44 | normal | 46 |
| NA14519:FXN | 9 | 83 | 9-9 | 69-98 | EXPANDED | 9 | 90 | 9-9 | 71-109 | EXPANDED | 6 | 6 | 4-6 | 4-6 | normal | 9 |
| NA15850:FXN | 41 | 169 | 41-41 | 140-196 | EXPANDED | 90 | 143 | 50-130 | 86-200 | EXPANDED | 6 | 233 | 6-8 | 215-247 | EXPANDED | 41 |
| NA15847:FXN | 9 | 111 | 9-9 | 89-131 | EXPANDED | 9 | 122 | 9-9 | 92-150 | EXPANDED | 7 | 8 | 5-10 | 5-10 | normal | 9 |
| NA15848:FXN | 8 | 91 | 8-8 | 74-107 | EXPANDED | 8 | 106 | 8-8 | 82-128 | EXPANDED | 8 | 158 | 8-8 | 140-176 | EXPANDED | 8 |
| NA16197:FXN | 41 | 164 | 41-41 | 138-191 | EXPANDED | 90 | 143 | 50-130 | 86-201 | EXPANDED | 6 | 201 | 6-6 | 189-213 | EXPANDED | 46 |
| NA16200:FXN | 9 | 97 | 9-9 | 79-114 | EXPANDED | 9 | 95 | 9-9 | 74-114 | EXPANDED | 6 | 20 | 6-11 | 6-21 | normal | 9 |
| NA16202:FXN | 8 | 76 | 8-8 | 62-89 | EXPANDED | 8 | 99 | 8-8 | 78-120 | EXPANDED | 8 | 8 | 8-8 | 8-8 | normal | 8 |
| NA16203:FXN | 22 | 143 | 22-22 | 119-167 | EXPANDED | 95 | 155 | 50-140 | 92-219 | EXPANDED | 6 | 304 | 6-6 | 290-323 | EXPANDED | 22 |
| NA16205:FXN | 92 | 135 | 50-108 | 76-159 | EXPANDED | 90 | 143 | 50-131 | 84-203 | EXPANDED | 6 | 174 | 6-6 | 162-190 | EXPANDED | 45 |
| NA16209:FXN | 98 | 148 | 50-114 | 80-172 | EXPANDED | 29 | 178 | 29-29 | 141-213 | EXPANDED | 6 | 232 | 6-6 | 218-248 | EXPANDED | 46 |
| NA16210:FXN | 109 | 168 | 50-127 | 89-196 | EXPANDED | 94 | 153 | 50-139 | 91-216 | EXPANDED | 6 | 235 | 6-6 | 222-248 | EXPANDED | 30 |
| NA16212:FXN | 8 | 111 | 8-8 | 91-132 | EXPANDED | 8 | 121 | 8-8 | 96-145 | EXPANDED | 6 | 8 | 6-8 | 7-9 | normal | 8 |
| NA16216:FXN | 41 | 184 | 41-41 | 152-215 | EXPANDED | 21 | 188 | 21-21 | 150-227 | EXPANDED | 10 | 192 | 7-10 | 181-203 | EXPANDED | 35 |
| NA16213:FXN | 8 | 95 | 8-8 | 77-113 | EXPANDED | 8 | 105 | 8-8 | 80-127 | EXPANDED | 8 | 144 | 8-8 | 124-162 | EXPANDED | 8 |
| NA16215:FXN | 79 | 109 | 50-93 | 63-129 | EXPANDED | 27 | 135 | 27-27 | 102-166 | EXPANDED | 6 | 209 | 6-27 | 187-225 | EXPANDED | 27 |
| NA16214:FXN | 98 | 146 | 50-116 | 78-172 | EXPANDED | 90 | 142 | 50-131 | 84-201 | EXPANDED | 6 | 192 | 6-7 | 173-213 | EXPANDED | 38 |
| NA16227:FXN | 107 | 166 | 50-127 | 86-195 | EXPANDED | 95 | 157 | 50-141 | 92-222 | EXPANDED | 6 | 274 | 6-6 | 250-295 | EXPANDED | 44 |
| NA16229:FXN | 19 | 94 | 19-19 | 76-112 | EXPANDED | 19 | 93 | 19-19 | 71-112 | EXPANDED | 6 | 21 | 6-6 | 14-23 | normal | 19 |
| NA16228:FXN | 6 | 180 | 6-6 | 149-210 | EXPANDED | 105 | 180 | 50-160 | 102-258 | EXPANDED | 6 | 220 | 6-6 | 205-233 | EXPANDED | 46 |
| NA16237:FXN | 8 | 80 | 8-8 | 64-94 | EXPANDED | 8 | 75 | 8-8 | 57-90 | EXPANDED | 8 | 19 | 8-8 | 16-20 | normal | 8 |
| NA16243:FXN | 114 | 181 | 50-135 | 92-211 | EXPANDED | 33 | 202 | 33-33 | 162-241 | EXPANDED | 6 | 192 | 6-6 | 180-207 | EXPANDED | 33 |
| NA16240:FXN | 9 | 110 | 9-9 | 89-130 | EXPANDED | 9 | 104 | 9-9 | 81-124 | EXPANDED | 6 | 23 | 6-9 | 9-23 | normal | 9 |
| NA16207:FXN | 8 | 176 | 8-8 | 148-205 | EXPANDED | 94 | 153 | 50-138 | 90-217 | EXPANDED | 6 | 214 | 6-6 | 197-226 | EXPANDED | 45 |
| NA06895:FMR1 | 23 | 0 | 23-23 | 0-0 | normal | 23 | 0 | 23-23 | 0-0 | normal | 21 | 0 | 14-23 | 0-0 | normal | 23 |
| NA04025:FMR1 | 133 | 0 | 108-155 | 0-0 | normal | 84 | 0 | 67-99 | 0-0 | normal | 21 | 0 | 8-21 | 0-0 | normal | 64 |
| NA04926:FMR1 | 127 | 0 | 105-150 | 0-0 | normal | 107 | 0 | 83-132 | 0-0 | normal | 17 | 0 | 12-17 | 0-0 | normal | 60 |
| NA05131:FMR1 | 207 | 0 | 173-240 | 0-0 | EXPANDED | 88 | 0 | 64-105 | 0-0 | normal | 39 | 0 | 38-39 | 0-0 | normal | 98 |
| NA05185:FMR1 | 127 | 0 | 102-149 | 0-0 | normal | 69 | 0 | 53-83 | 0-0 | normal | 26 | 0 | 19-26 | 0-0 | normal | 61 |
| NA09145:FMR1 | 135 | 0 | 109-158 | 0-0 | normal | 85 | 0 | 65-103 | 0-0 | normal | 17 | 0 | 17-20 | 0-0 | normal | 70 |
| NA09237:FMR1 | 131 | 0 | 108-154 | 0-0 | normal | 87 | 0 | 67-104 | 0-0 | normal | 24 | 0 | 11-24 | 0-0 | normal | 67 |
| NA07063:FMR1 | 32 | 141 | 32-32 | 123-159 | normal | 33 | 79 | 33-33 | 65-91 | normal | 33 | 62 | 31-57 | 56-69 | normal | 32 |
| NA07539:FMR1 | 23 | 0 | 23-23 | 0-0 | normal | 23 | 0 | 23-23 | 0-0 | normal | 23 | 0 | 20-23 | 0-0 | normal | 23 |
| NA06890:FMR1 | 30 | 0 | 30-30 | 0-0 | normal | 30 | 0 | 30-30 | 0-0 | normal | 30 | 0 | 27-31 | 0-0 | normal | 30 |
| NA06905:FMR1 | 23 | 56 | 23-23 | 50-63 | normal | 23 | 60 | 23-23 | 50-68 | normal | 23 | 30 | 17-31 | 18-31 | normal | 39 |
| NA07536:FMR1 | 23 | 0 | 23-23 | 0-0 | normal | 23 | 0 | 23-23 | 0-0 | normal | 23 | 0 | 22-24 | 0-0 | normal | 23 |
| NA07540:FMR1 | 23 | 29 | 23-23 | 29-29 | normal | 23 | 29 | 23-23 | 29-29 | normal | 23 | 29 | 20-29 | 29-29 | normal | 23 |
| NA07542:FMR1 | 23 | 0 | 23-23 | 0-0 | normal | 23 | 0 | 23-23 | 0-0 | normal | 23 | 0 | 23-23 | 0-0 | normal | 23 |
| NA06910:FMR1 | 30 | 78 | 30-30 | 63-91 | normal | 30 | 77 | 30-30 | 59-92 | normal | 30 | 55 | 30-34 | 30-61 | normal | 30 |
| NA06894:FMR1 | 30 | 64 | 30-30 | 52-73 | normal | 31 | 63 | 31-31 | 50-74 | normal | 27 | 27 | 27-31 | 27-31 | normal | 26 |
| NA07541:FMR1 | 29 | 31 | 29-29 | 31-31 | normal | 29 | 52 | 29-29 | 50-55 | normal | 29 | 31 | 14-31 | 14-31 | normal | 29 |
| NA07175:FMR1 | 23 | 30 | 23-23 | 30-30 | normal | 23 | 45 | 23-23 | 23-45 | normal | 23 | 24 | 20-24 | 20-24 | normal | 23 |
| NA06889:FMR1 | 23 | 30 | 23-23 | 30-30 | normal | 23 | 30 | 23-23 | 30-30 | normal | 30 | 30 | 21-30 | 21-30 | normal | 23 |
| NA06893:FMR1 | 23 | 30 | 23-23 | 30-30 | normal | 23 | 30 | 23-23 | 30-30 | normal | 23 | 30 | 8-23 | 8-30 | normal | 23 |
| NA06896:FMR1 | 23 | 89 | 23-23 | 71-104 | normal | 23 | 75 | 23-23 | 57-90 | normal | 23 | 24 | 23-23 | 23-26 | normal | 23 |
| NA07538:FMR1 | 29 | 29 | 29-29 | 29-29 | normal | 29 | 52 | 29-29 | 50-55 | normal | 27 | 30 | 26-31 | 26-31 | normal | 29 |
| NA07537:FMR1 | 29 | 84 | 29-29 | 69-98 | normal | 29 | 70 | 29-29 | 56-81 | normal | 29 | 57 | 28-38 | 46-65 | normal | 29 |
| NA06897:FMR1 | 102 | 0 | 80-121 | 0-0 | normal | 68 | 0 | 52-81 | 0-0 | normal | 18 | 0 | 17-20 | 0-0 | normal | 60 |
| NA07174:FMR1 | 30 | 0 | 30-30 | 0-0 | normal | 52 | 0 | 50-55 | 0-0 | normal | 33 | 0 | 29-33 | 0-0 | normal | 31 |
| NA06903:FMR1 | 24 | 57 | 24-24 | 50-65 | normal | 24 | 61 | 24-24 | 50-72 | normal | 24 | 53 | 20-24 | 20-59 | normal | 24 |
| NA07543:FMR1 | 20 | 29 | 20-20 | 29-29 | normal | 20 | 29 | 20-20 | 29-29 | normal | 20 | 20 | 20-29 | 20-29 | normal | 20 |
| NA06852:FMR1 | 100 | 0 | 80-120 | 0-0 | normal | 68 | 0 | 54-80 | 0-0 | normal | 20 | 0 | 20-75 | 0-0 | normal | 59 |
| NA06891:FMR1 | 85 | 0 | 68-101 | 0-0 | normal | 79 | 0 | 60-95 | 0-0 | normal | 17 | 0 | 17-20 | 0-0 | normal | 54 |
| NA06907:FMR1 | 29 | 75 | 29-29 | 61-88 | normal | 29 | 71 | 29-29 | 52-86 | normal | 32 | 55 | 28-55 | 29-62 | normal | 18 |
| NA06906:FMR1 | 67 | 0 | 56-78 | 0-0 | normal | 67 | 0 | 52-80 | 0-0 | normal | 31 | 0 | 20-32 | 0-0 | normal | 57 |
| NA06892:FMR1 | 73 | 0 | 61-86 | 0-0 | normal | 65 | 0 | 52-76 | 0-0 | normal | 21 | 0 | 20-78 | 0-0 | normal | 61 |
| NA06904:FMR1 | 24 | 29 | 24-24 | 29-29 | normal | 29 | 46 | 29-29 | 29-46 | normal | 29 | 29 | 28-31 | 28-31 | normal | 29 |
| NA06968:FMR1 | 33 | 80 | 33-33 | 65-93 | normal | 33 | 87 | 33-33 | 67-104 | normal | 20 | 57 | 20-20 | 32-64 | normal | 33 |
| NA07294:FMR1 | 132 | 0 | 108-156 | 0-0 | normal | 86 | 0 | 63-106 | 0-0 | normal | 24 | 0 | 6-24 | 0-0 | normal | 65 |
| NA09316:FMR1 | 94 | 0 | 77-111 | 0-0 | normal | 82 | 0 | 61-100 | 0-0 | normal | 27 | 0 | 20-27 | 0-0 | normal | 61 |
| NA09317:FMR1 | 78 | 0 | 63-91 | 0-0 | normal | 64 | 0 | 50-75 | 0-0 | normal | 63 | 0 | 41-63 | 0-0 | normal | 42 |
| NA09497:FMR1 | 110 | 0 | 89-129 | 0-0 | normal | 69 | 0 | 52-83 | 0-0 | normal | 28 | 0 | 14-28 | 0-0 | normal | 53 |
| NA07730:FMR1 | 117 | 0 | 96-138 | 0-0 | normal | 88 | 0 | 69-105 | 0-0 | normal | 16 | 0 | 8-16 | 0-0 | normal | 64 |
| NA03200:FMR1 | 185 | 0 | 154-216 | 0-0 | normal | 121 | 0 | 95-145 | 0-0 | normal | 18 | 0 | 10-18 | 0-0 | normal | 89 |
| NA20235:FMR1 | 29 | 40 | 29-29 | 40-40 | normal | 29 | 40 | 29-29 | 29-40 | normal | 29 | 29 | 28-32 | 28-32 | normal | 29 |
| NA20238:FMR1 | 29 | 30 | 29-29 | 30-30 | normal | 29 | 52 | 29-29 | 50-54 | normal | 29 | 29 | 28-29 | 28-29 | normal | 29 |
| NA20237:FMR1 | 74 | 0 | 60-87 | 0-0 | normal | 69 | 0 | 56-79 | 0-0 | normal | 62 | 0 | 20-64 | 0-0 | normal | 32 |
| NA20239:FMR1 | 20 | 81 | 20-20 | 65-94 | normal | 20 | 73 | 20-20 | 58-86 | normal | 20 | 65 | 20-20 | 57-75 | normal | 20 |
| NA20242:FMR1 | 30 | 68 | 30-30 | 56-80 | normal | 30 | 63 | 30-30 | 50-74 | normal | 30 | 30 | 29-32 | 29-32 | normal | 30 |
| NA20243:FMR1 | 29 | 29 | 29-29 | 29-29 | normal | 29 | 31 | 29-29 | 29-38 | normal | 29 | 29 | 17-30 | 17-30 | normal | 29 |
| NA20230:FMR1 | 58 | 0 | 50-65 | 0-0 | normal | 59 | 0 | 50-66 | 0-0 | normal | 55 | 0 | 23-60 | 0-0 | normal | 52 |
| NA20232:FMR1 | 65 | 0 | 54-73 | 0-0 | normal | 55 | 0 | 50-60 | 0-0 | normal | 22 | 0 | 20-30 | 0-0 | normal | 39 |
| NA20233:FMR1 | 88 | 0 | 71-103 | 0-0 | normal | 82 | 0 | 61-99 | 0-0 | normal | 118 | 0 | 20-118 | 0-0 | normal | 33 |
| NA20234:FMR1 | 31 | 54 | 31-31 | 50-58 | normal | 31 | 43 | 31-31 | 31-45 | normal | 29 | 29 | 28-31 | 28-31 | normal | 31 |
| NA20236:FMR1 | 31 | 54 | 31-31 | 50-58 | normal | 52 | 55 | 50-55 | 50-60 | normal | 23 | 27 | 20-30 | 20-30 | normal | 37 |
| NA20231:FMR1 | 56 | 0 | 50-63 | 0-0 | normal | 69 | 0 | 53-81 | 0-0 | normal | 33 | 0 | 24-35 | 0-0 | normal | 57 |
| NA20240:FMR1 | 66 | 84 | 50-75 | 54-101 | normal | 64 | 79 | 50-79 | 53-106 | normal | 28 | 60 | 4-57 | 53-68 | normal | 34 |
| NA20241:FMR1 | 29 | 73 | 29-29 | 59-84 | normal | 29 | 71 | 29-29 | 53-86 | normal | 29 | 29 | 29-29 | 29-29 | normal | 29 |
| NA20244:FMR1 | 35 | 0 | 35-35 | 0-0 | normal | 52 | 0 | 50-55 | 0-0 | normal | 36 | 0 | 30-37 | 0-0 | normal | 37 |
| NA07862:FMR1 | 117 | 0 | 96-139 | 0-0 | normal | 87 | 0 | 67-105 | 0-0 | normal | 20 | 0 | 20-20 | 0-0 | normal | 68 |
| NA13509:HTT | 15 | 84 | 15-15 | 69-99 | EXPANDED | 15 | 83 | 15-15 | 63-100 | EXPANDED | 18 | 75 | 18-18 | 64-88 | EXPANDED | 15 |
| NA13515:HTT | 16 | 82 | 16-16 | 68-97 | EXPANDED | 16 | 76 | 16-16 | 60-88 | EXPANDED | 16 | 67 | 16-16 | 56-78 | EXPANDED | 16 |
| NA13507:HTT | 15 | 58 | 15-15 | 50-66 | EXPANDED | 15 | 57 | 15-15 | 50-64 | EXPANDED | 15 | 40 | 15-15 | 29-41 | EXPANDED | 15 |
| NA13508:HTT | 22 | 63 | 22-22 | 52-71 | EXPANDED | 22 | 57 | 22-22 | 50-65 | EXPANDED | 22 | 62 | 22-22 | 52-73 | EXPANDED | 22 |
| NA13510:HTT | 15 | 58 | 15-15 | 50-65 | EXPANDED | 15 | 56 | 15-15 | 50-62 | EXPANDED | 15 | 33 | 15-19 | 19-36 | normal | 15 |
| NA13511:HTT | 43 | 56 | 43-43 | 50-62 | EXPANDED | 50 | 52 | 50-50 | 50-54 | EXPANDED | 19 | 46 | 5-19 | 42-49 | EXPANDED | 5 |
| NA13512:HTT | 16 | 52 | 16-16 | 50-54 | EXPANDED | 16 | 43 | 16-16 | 16-50 | EXPANDED | 16 | 41 | 16-16 | 35-41 | EXPANDED | 16 |
| NA13513:HTT | 15 | 62 | 15-15 | 52-70 | EXPANDED | 15 | 52 | 15-15 | 50-54 | EXPANDED | 15 | 59 | 15-15 | 52-69 | EXPANDED | 15 |
| NA13514:HTT | 15 | 56 | 15-15 | 50-62 | EXPANDED | 15 | 52 | 15-15 | 50-54 | EXPANDED | 18 | 66 | 15-18 | 54-77 | EXPANDED | 15 |
| NA13503:HTT | 17 | 43 | 17-17 | 43-48 | EXPANDED | 17 | 43 | 17-17 | 17-50 | EXPANDED | 20 | 73 | 19-20 | 60-88 | EXPANDED | 17 |
| NA13504:HTT | 16 | 52 | 16-16 | 50-54 | EXPANDED | 16 | 52 | 16-16 | 50-55 | EXPANDED | 16 | 39 | 16-19 | 35-39 | normal | 16 |
| NA13505:HTT | 22 | 60 | 22-22 | 52-68 | EXPANDED | 22 | 52 | 22-22 | 50-54 | EXPANDED | 22 | 54 | 22-22 | 33-61 | EXPANDED | 22 |
| NA13506:HTT | 17 | 59 | 17-17 | 50-66 | EXPANDED | 17 | 46 | 17-17 | 17-50 | EXPANDED | 19 | 53 | 18-19 | 45-57 | EXPANDED | 17 |
| NA06926:ATXN1 | 30 | 61 | 30-30 | 52-70 | EXPANDED | 30 | 53 | 30-30 | 50-56 | EXPANDED | 33 | 62 | 28-59 | 52-73 | EXPANDED | 29 |
| NA13536:ATXN1 | 32 | 38 | 32-32 | 35-42 | normal | 32 | 43 | 32-32 | 32-50 | EXPANDED | 31 | 36 | 31-31 | 33-38 | normal | 31 |
| NA13537:ATXN1 | 33 | 69 | 33-33 | 56-80 | EXPANDED | 54 | 57 | 50-58 | 50-65 | EXPANDED | 31 | 61 | 31-58 | 52-71 | EXPANDED | 43 |
| NA06151:ATXN3 | 54 | 61 | 50-59 | 50-71 | EXPANDED | 21 | 57 | 21-21 | 50-64 | normal | 18 | 58 | 18-18 | 29-68 | normal | 18 |
| NA23709:AR | 52 | 0 | 50-54 | 0-0 | EXPANDED | 46 | 0 | 46-48 | 0-0 | EXPANDED | 28 | 0 | 24-48 | 0-0 | normal | 46 |
| hg19C9ORF721000:C9ORF72 | 3 | 1052 | 3-3 | 976-1128 | EXPANDED | 3 | 78 | 3-3 | 60-97 | EXPANDED | 3 | 3 | 3-3 | 3-3 | normal | 3 |
| hg19C9ORF72500:C9ORF72 | 3 | 474 | 3-3 | 422-524 | EXPANDED | 3 | 93 | 3-3 | 70-114 | EXPANDED | 3 | 3 | 3-3 | 3-3 | normal | 3 |
| hg19C9ORF7260:C9ORF72 | 3 | 70 | 3-3 | 53-85 | EXPANDED | 3 | 70 | 3-3 | 52-85 | EXPANDED | 3 | 3 | 3-3 | 3-3 | normal | 3 |
| hg19FMR21000:FMR2 | 15 | 163 | 15-15 | 126-196 | normal | 20 | 161 | 20-20 | 123-196 | normal | 15 | 15 | 15-15 | 15-15 | normal | 15 |
| hg19FMR2200:FMR2 | 15 | 160 | 15-15 | 123-193 | normal | 20 | 162 | 20-20 | 125-196 | normal | 15 | 15 | 12-15 | 12-15 | normal | 15 |
| hg19FMR2500:FMR2 | 15 | 150 | 15-15 | 113-183 | normal | 20 | 153 | 20-20 | 116-187 | normal | 15 | 15 | 15-15 | 15-15 | normal | 15 |

| sampleID:gene | TRED:a2 | TRED:CI-a1 | TRED:CI-a2 | TRED:status | lobSTR:a1 | lobSTR:a2 | lobSTR:status | HipSTR:a1 | HipSTR:a2 | HipSTR:status | RepeatSeq:a1 | RepeatSeq:a2 | RepeatSeq:status | STRetch:p-value | STRetch:repeatUnit | STRetch:pstatus | exSTRa:p-value | exSTRa:pstatus |
|---|---|---|---|---|---|---|---|---|---|---|---|---|---|---|---|---|---|---|
| NA06075:DMPK | 67 | 12-12 | 58-85 | EXPANDED | 12 | 12 | normal | . | . | . | 12 | 0 | normal | 3.7277720557467967e-41 | 108.34164576556 | EXPANDED | 3.229009e-05 | EXPANDED |
| NA04567:DMPK | 98 | 21-21 | 83-121 | EXPANDED | 21 | 21 | normal | 20 | 20 | normal | 21 | 0 | normal | 7.892057562359336e-47 | 131.09690768412807 | EXPANDED | 8.072524e-06 | EXPANDED |
| NA05164:DMPK | 81 | 21-21 | 71-99 | EXPANDED | 21 | 21 | normal | . | . | . | 21 | 20 | normal | 3.544050479922964e-35 | 88.81836861417233 | EXPANDED | 8.072524e-06 | EXPANDED |
| NA04648:DMPK | 89 | 5-5 | 76-106 | EXPANDED | 5 | 5 | normal | . | . | . | 5 | 0 | normal | 3.5049462133950315e-49 | 142.5357219746143 | EXPANDED | 8.072524e-06 | EXPANDED |
| NA05152:DMPK | 88 | 5-5 | 77-105 | EXPANDED | 5 | 5 | normal | . | . | . | 5 | 0 | normal | 3.678098712788432e-42 | 112.76768911699213 | EXPANDED | 8.072524e-06 | EXPANDED |
| NA23378:DMPK | 93 | 22-22 | 80-111 | EXPANDED | 22 | 22 | normal | . | . | . | . | . | . | 1.1734388904983037e-41 | 110.86114291198635 | EXPANDED | 8.072524e-06 | EXPANDED |
| NA23374:DMPK | 86 | 5-5 | 74-103 | EXPANDED | 5 | 5 | normal | . | . | . | 5 | 0 | normal | 1.3769131465391494e-42 | 113.246610958481 | EXPANDED | 8.072524e-06 | EXPANDED |
| NA23300:DMPK | 102 | 5-5 | 83-132 | EXPANDED | 5 | 5 | normal | 19 | 4 | normal | 5 | 0 | normal | 5.512189571954157e-36 | 90.77661613272883 | EXPANDED | 8.072524e-06 | EXPANDED |
| NA03986:DMPK | 86 | 12-12 | 74-105 | EXPANDED | 12 | 12 | normal | . | . | . | 12 | 0 | normal | 2.3628129568746877e-35 | 88.81836861417233 | EXPANDED | 8.072524e-06 | EXPANDED |
| NA03989:DMPK | 121 | 13-13 | 98-154 | EXPANDED | 13 | 13 | normal | . | . | . | 13 | 0 | normal | 4.970730518608572e-43 | 116.12357307491801 | EXPANDED | 8.072524e-06 | EXPANDED |
| NA03990:DMPK | 70 | 14-14 | 61-85 | EXPANDED | 14 | 14 | normal | . | . | . | 14 | 0 | normal | 3.353893516338058e-46 | 129.13981546383863 | EXPANDED | 8.072524e-06 | EXPANDED |
| NA03696:DMPK | 102 | 12-12 | 86-129 | EXPANDED | 12 | 12 | normal | . | . | . | 12 | 0 | normal | 9.630077613467448e-35 | 86.96491278295703 | EXPANDED | 8.072524e-06 | EXPANDED |
| NA03759:DMPK | 102 | 14-14 | 85-127 | EXPANDED | 14 | 14 | normal | . | . | . | 14 | 0 | normal | 2.8804922967648865e-40 | 105.7526764442294 | EXPANDED | 8.072524e-06 | EXPANDED |
| NA04034:DMPK | 84 | 12-12 | 71-109 | EXPANDED | 12 | 12 | normal | . | . | . | 12 | 0 | normal | 1.2054130082764893e-37 | 97.69835776214023 | EXPANDED | 8.072524e-06 | EXPANDED |
| NA03697:DMPK | 97 | 12-12 | 84-117 | EXPANDED | 12 | 12 | normal | 19 | 11 | normal | 12 | 0 | normal | 5.467350112003751e-44 | 118.72233913960766 | EXPANDED | 8.072524e-06 | EXPANDED |
| NA03132:DMPK | 95 | 5-5 | 80-118 | EXPANDED | 5 | 5 | normal | . | . | . | 5 | 0 | normal | 1.3845178527636742e-38 | 99.85652117365396 | EXPANDED | 8.072524e-06 | EXPANDED |
| NA03756:DMPK | 103 | 13-13 | 87-126 | EXPANDED | 13 | 13 | normal | . | . | . | 13 | 0 | normal | 1.8310962160517065e-36 | 92.84874494897238 | EXPANDED | 8.072524e-06 | EXPANDED |
| NA13716:ATN1 | 57 | 16-16 | 52-70 | EXPANDED | 20 | 20 | normal | . | . | . | 16 | 0 | normal | 1.6582902531602304e-22 | 54.70200868905826 | EXPANDED | 8.072524e-06 | EXPANDED |
| NA13717:ATN1 | 57 | 15-15 | 52-66 | EXPANDED | 19 | 19 | normal | . | . | . | 15 | 0 | normal | 7.401094261799903e-25 | 60.27639850519684 | EXPANDED | 8.072524e-06 | EXPANDED |
| NA03816:FXN | 187 | 45-128 | 114-220 | EXPANDED | . | . | . | . | . | . | . | . | . | 2.9832369461813087e-05 | 15.578422974583995 | EXPANDED | . | . |
| NA04079:FXN | 216 | 48-142 | 129-248 | EXPANDED | . | . | . | . | . | . | . | . | . | 1.1833946335987388e-05 | 16.198337727475767 | EXPANDED | . | . |
| NA14519:FXN | 86 | 9-9 | 70-108 | EXPANDED | 10 | 10 | normal | . | . | . | . | . | . | 0.7401573219255145 | 10.407802243504225 | normal | . | . |
| NA15850:FXN | 208 | 39-41 | 171-258 | EXPANDED | . | . | . | . | . | . | . | . | . | 0.7401099148180085 | 10.407802243504225 | normal | . | . |
| NA15847:FXN | 121 | 9-9 | 98-167 | EXPANDED | 10 | 10 | normal | 21 | 21 | normal | . | . | . | 8.995220715831744e-07 | 17.372663713190747 | EXPANDED | . | . |
| NA15848:FXN | 97 | 8-8 | 80-124 | EXPANDED | 8 | 8 | normal | . | . | . | . | . | . | 7.12961035007329e-06 | 16.468954638639772 | EXPANDED | . | . |
| NA16197:FXN | 205 | 46-72 | 159-250 | EXPANDED | . | . | . | . | . | . | . | . | . | 0.00041235062995444654 | 14.53741359552382 | EXPANDED | . | . |
| NA16200:FXN | 86 | 9-9 | 72-110 | EXPANDED | 10 | 10 | normal | . | . | . | . | . | . | 0.0009865965957953724 | 14.096233139666829 | EXPANDED | . | . |
| NA16202:FXN | 90 | 8-8 | 75-117 | EXPANDED | . | . | . | 18 | 18 | normal | . | . | . | 2.003949716253737e-07 | 18.315063337850766 | EXPANDED | . | . |
| NA16203:FXN | 222 | 22-22 | 184-271 | EXPANDED | . | . | . | . | . | . | . | . | . | 0.20362351863908823 | 11.860705206113849 | normal | . | . |
| NA16205:FXN | 184 | 45-71 | 141-227 | EXPANDED | . | . | . | . | . | . | . | . | . | 0.0003829426595419 | 14.53741359552382 | EXPANDED | 8.072524e-06 | EXPANDED |
| NA16209:FXN | 194 | 46-92 | 135-237 | EXPANDED | . | . | . | . | . | . | 6 | 0 | normal | 0.000324955901415001 | 14.53741359552382 | EXPANDED | . | . |
| NA16210:FXN | 225 | 30-30 | 186-274 | EXPANDED | . | . | . | . | . | . | . | . | . | 0.09558598632894726 | 12.012331779984216 | normal | . | . |
| NA16212:FXN | 122 | 8-8 | 101-158 | EXPANDED | . | . | . | . | . | . | . | . | . | 0.04898181202316149 | 12.339956149206124 | EXPANDED | . | . |
| NA16216:FXN | 213 | 35-35 | 176-273 | EXPANDED | . | . | . | . | . | . | 6 | 0 | normal | 5.549538395112172e-13 | 25.039922789984015 | EXPANDED | 4.601338e-04 | EXPANDED |
| NA16213:FXN | 91 | 8-8 | 75-121 | EXPANDED | 8 | 8 | normal | . | . | . | . | . | . | 0.026968005133449918 | 12.704644354116141 | EXPANDED | . | . |
| NA16215:FXN | 131 | 27-27 | 101-180 | EXPANDED | . | . | . | . | . | . | . | . | . | 0.001931481583624665 | 13.573372575142738 | EXPANDED | 8.072524e-06 | EXPANDED |
| NA16214:FXN | 195 | 38-38 | 160-243 | EXPANDED | . | . | . | . | . | . | . | . | . | 0.03477864273660233 | 12.517287231386582 | EXPANDED | 6.458019e-04 | EXPANDED |
| NA16227:FXN | 201 | 44-74 | 154-249 | EXPANDED | . | . | . | . | . | . | 6 | 0 | normal | 4.832109811887195e-09 | 19.73312766164753 | EXPANDED | . | . |
| NA16229:FXN | 100 | 19-19 | 79-125 | EXPANDED | . | . | . | . | . | . | 6 | 0 | normal | 0.7837364129909655 | 10.168047022663163 | normal | 1.614505e-05 | EXPANDED |
| NA16228:FXN | 257 | 48-162 | 149-283 | EXPANDED | . | . | . | . | . | . | 6 | 0 | normal | 1.9338354731690887e-08 | 19.349363937747043 | EXPANDED | . | . |
| NA16237:FXN | 83 | 8-8 | 69-109 | EXPANDED | 8 | 8 | normal | . | . | . | . | . | . | 1.3475567782712614e-05 | 15.87882609688239 | EXPANDED | . | . |
| NA16243:FXN | 229 | 33-33 | 187-279 | EXPANDED | . | . | . | . | . | . | . | . | . | 1.784802306823897e-08 | 19.349363937747043 | EXPANDED | . | . |
| NA16240:FXN | 93 | 9-9 | 79-116 | EXPANDED | 10 | 10 | normal | . | . | . | . | . | . | 1.9304783525437763e-06 | 17.054861688424708 | EXPANDED | . | . |
| NA16207:FXN | 240 | 45-61 | 196-289 | EXPANDED | . | . | . | . | . | . | . | . | . | 9.581449663113758e-11 | 22.10723736424742 | EXPANDED | . | . |
| NA06895:FMR1 | 23 | 23-23 | 23-23 | normal | 28 | 28 | normal | . | . | . | . | . | . | . | . | . | . | . |
| NA04025:FMR1 | 64 | 56-83 | 56-83 | normal | . | . | . | . | . | . | . | . | . | 0.5902410425783222 | 39.82730357396629 | normal | . | . |
| NA04926:FMR1 | 60 | 53-78 | 53-78 | normal | . | . | . | . | . | . | . | . | . | 0.22010201737603807 | 43.732265042149834 | normal | . | . |
| NA05131:FMR1 | 98 | 77-133 | 77-133 | normal | . | . | . | . | . | . | 20 | 0 | normal | 0.015216551830441167 | 52.27418861391992 | EXPANDED | . | . |
| NA05185:FMR1 | 61 | 53-83 | 53-83 | normal | . | . | . | . | . | . | 20 | 20 | normal | . | . | . | . | . |
| NA09145:FMR1 | 70 | 59-93 | 59-93 | normal | . | . | . | . | . | . | 20 | 0 | normal | 0.03163493537181473 | 48.98344508707048 | EXPANDED | . | . |
| NA09237:FMR1 | 67 | 58-90 | 58-90 | normal | . | . | . | . | . | . | 20 | 0 | normal | 0.7564862689737221 | 33.77620843200008 | normal | . | . |
| NA07063:FMR1 | 59 | 32-32 | 54-69 | normal | . | . | . | . | . | . | . | . | . | 0.9988193918748451 | 31.590980402100367 | normal | . | . |
| NA07539:FMR1 | 23 | 23-23 | 23-23 | normal | . | . | . | . | . | . | . | . | . | . | . | . | . | . |
| NA06890:FMR1 | 30 | 30-30 | 30-30 | normal | 35 | 35 | normal | . | . | . | . | . | . | 0.5077802907884078 | 37.846232390972816 | normal | . | . |
| NA06905:FMR1 | 56 | 39-55 | 41-71 | normal | . | . | . | . | . | . | . | . | . | 0.7078028638195785 | 29.538138645989306 | normal | . | . |
| NA07536:FMR1 | 23 | 23-23 | 23-23 | normal | 28 | 28 | normal | . | . | . | 23 | 0 | normal | 0.7556262468044473 | 31.517287231386582 | normal | . | . |
| NA07540:FMR1 | 29 | 23-23 | 29-29 | normal | 28 | 34 | normal | 23 | 29 | normal | 29 | 0 | normal | . | . | . | . | . |
| NA07542:FMR1 | 23 | 23-23 | 23-23 | normal | 28 | 28 | normal | . | . | . | . | . | . | . | . | . | . | . |
| NA06910:FMR1 | 38 | 30-30 | 38-69 | normal | 35 | 35 | normal | 30 | 30 | normal | 30 | 0 | normal | 0.6689549129386576 | 29.822937443502347 | normal | . | . |
| NA06894:FMR1 | 30 | 26-30 | 30-61 | normal | 35 | 35 | normal | . | . | . | 20 | 30 | normal | 0.013941195829359163 | 52.27418861391992 | EXPANDED | . | . |
| NA07541:FMR1 | 31 | 29-29 | 31-31 | normal | 12 | 34 | normal | . | . | . | 29 | 0 | normal | . | . | . | . | . |
| NA07175:FMR1 | 30 | 23-23 | 30-31 | normal | 28 | 35 | normal | 23 | 30 | normal | 30 | 20 | normal | . | . | . | . | . |
| NA06889:FMR1 | 30 | 23-23 | 30-30 | normal | 28 | 35 | normal | 23 | 30 | normal | 30 | 20 | normal | . | . | . | . | . |
| NA06893:FMR1 | 30 | 23-23 | 30-30 | normal | 28 | 35 | normal | . | . | . | 30 | 0 | normal | . | . | . | . | . |
| NA06896:FMR1 | 59 | 23-23 | 52-76 | normal | 28 | 28 | normal | 20 | 23 | normal | . | . | . | . | . | . | . | . |
| NA07538:FMR1 | 29 | 24-29 | 29-29 | normal | 34 | 34 | normal | 29 | 29 | normal | . | . | . | . | . | . | . | . |
| NA07537:FMR1 | 41 | 29-29 | 41-60 | normal | 34 | 34 | normal | 29 | 29 | normal | 29 | 0 | normal | 0.8163119020450283 | 37.64184839737077 | normal | . | . |
| NA06897:FMR1 | 60 | 54-80 | 54-80 | normal | . | . | . | . | . | . | . | . | . | 0.41415199905733374 | 37.846232390972816 | normal | . | . |
| NA07174:FMR1 | 31 | 31-54 | 31-54 | normal | . | . | . | . | . | . | 30 | 20 | normal | . | . | . | . | . |
| NA06903:FMR1 | 45 | 24-24 | 45-58 | normal | . | . | . | 20 | 24 | normal | 20 | 0 | normal | 0.5587675979859662 | 33.09623313966683 | normal | . | . |
| NA07543:FMR1 | 29 | 20-20 | 20-29 | normal | 25 | 34 | normal | 20 | 20 | normal | 20 | 29 | normal | 0.41215555482648236 | 41.107237364247425 | normal | . | . |
| NA06852:FMR1 | 59 | 53-77 | 53-77 | normal | . | . | . | . | . | . | . | . | . | . | . | . | . | . |
| NA06891:FMR1 | 54 | 37-70 | 37-70 | normal | . | . | . | . | . | . | . | . | . | 0.03218152699008422 | 48.595907134202605 | EXPANDED | . | . |
| NA06907:FMR1 | 29 | 18-29 | 29-66 | normal | 23 | 23 | normal | 20 | 29 | normal | 29 | 0 | normal | 0.7379405686343744 | 36.37266371319075 | normal | . | . |
| NA06906:FMR1 | 57 | 35-74 | 35-74 | normal | . | . | . | . | . | . | . | . | . | 0.7087317570123631 | 29.538138645989306 | normal | . | . |
| NA06892:FMR1 | 61 | 54-75 | 54-75 | normal | . | . | . | . | . | . | . | . | . | 0.7375458024466894 | 36.37266371319075 | normal | . | . |
| NA06904:FMR1 | 29 | 29-29 | 29-29 | normal | . | . | . | . | . | . | 29 | 20 | normal | . | . | . | . | . |
| NA06968:FMR1 | 56 | 33-33 | 45-69 | normal | . | . | . | . | . | . | . | . | . | 0.7084976341740767 | 29.538138645989306 | normal | . | . |
| NA07294:FMR1 | 65 | 56-88 | 56-88 | normal | . | . | . | . | . | . | 20 | 0 | normal | 0.7559035339336784 | 31.517287231386582 | normal | . | . |
| NA09316:FMR1 | 61 | 53-83 | 53-83 | normal | . | . | . | . | . | . | . | . | . | 0.7395014071948224 | 36.37266371319075 | normal | . | . |
| NA09317:FMR1 | 42 | 42-58 | 42-58 | normal | . | . | . | . | . | . | . | . | . | 0.7833549061692991 | 29.168047022663163 | normal | 6.861645e-04 | EXPANDED |
| NA09497:FMR1 | 53 | 42-67 | 42-67 | normal | . | . | . | . | . | . | . | . | . | 0.7846427392786426 | 35.75406620144316 | normal | . | . |
| NA07730:FMR1 | 64 | 56-83 | 56-83 | normal | . | . | . | . | . | . | . | . | . | 0.7837620406312714 | 29.168047022663163 | normal | . | . |
| NA03200:FMR1 | 89 | 73-116 | 73-116 | normal | . | . | . | . | . | . | 20 | 0 | normal | 0.707116475336365 | 39.82730357396629 | normal | . | . |
| NA20235:FMR1 | 35 | 29-29 | 35-35 | normal | 34 | 34 | normal | 20 | 29 | normal | 29 | 30 | normal | 0.07192203936589792 | 46.506946748334926 | normal | . | . |
| NA20238:FMR1 | 30 | 29-29 | 30-30 | normal | 34 | 34 | normal | 29 | 30 | normal | 29 | 0 | normal | . | . | . | . | . |
| NA20237:FMR1 | 32 | 32-60 | 32-60 | normal | . | . | . | . | . | . | . | . | . | 0.6903067804358576 | 34.578422974583994 | normal | . | . |
| NA20239:FMR1 | 58 | 20-20 | 52-75 | normal | . | . | . | 20 | 20 | normal | 20 | 0 | normal | 0.7845494239378982 | 31.339956149206124 | normal | . | . |
| NA20242:FMR1 | 52 | 30-30 | 45-62 | normal | 35 | 35 | normal | . | . | . | 30 | 20 | normal | 0.7390321520072598 | 36.37266371319075 | normal | . | . |
| NA20243:FMR1 | 31 | 29-29 | 31-59 | normal | . | . | . | . | . | . | 29 | 0 | normal | . | . | . | . | . |
| NA20230:FMR1 | 52 | 46-62 | 46-62 | normal | . | . | . | . | . | . | 20 | 0 | normal | 0.7824676871531359 | 29.168047022663163 | normal | . | . |
| NA20232:FMR1 | 39 | 39-56 | 39-56 | normal | . | . | . | . | . | . | 20 | 0 | normal | 0.7393573059201957 | 31.70464435411614 | normal | 8.072524e-06 | EXPANDED |
| NA20233:FMR1 | 33 | 33-33 | 33-33 | normal | . | . | . | . | . | . | 20 | 0 | normal | 0.6900587874390874 | 29.67627776425815 | normal | . | . |
| NA20234:FMR1 | 55 | 31-31 | 45-66 | normal | 36 | 36 | normal | 31 | 31 | normal | 31 | 0 | normal | 0.7564499749711414 | 29.28462894716124 | normal | . | . |
| NA20236:FMR1 | 37 | 37-37 | 37-37 | normal | . | . | . | . | . | . | 30 | 0 | normal | 0.739991385567443 | 29.407802243504225 | normal | . | . |
| NA20231:FMR1 | 57 | 34-74 | 34-74 | normal | . | . | . | . | . | . | . | . | . | 0.7839152240863776 | 33.537413595523816 | normal | . | . |
| NA20240:FMR1 | 64 | 34-61 | 55-85 | normal | . | . | . | . | . | . | . | . | . | 0.6898122238452001 | 29.67627776425815 | normal | . | . |
| NA20241:FMR1 | 29 | 29-29 | 29-57 | normal | 34 | 34 | normal | 29 | 29 | normal | 29 | 0 | normal | . | . | . | . | . |
| NA20244:FMR1 | 37 | 37-55 | 37-55 | normal | . | . | . | . | . | . | . | . | . | 0.7092864134341791 | 29.538138645989306 | normal | . | . |
| NA07862:FMR1 | 68 | 59-90 | 59-90 | normal | . | . | . | . | . | . | 20 | 0 | normal | 0.7387311220150747 | 38.73312766164753 | normal | . | . |
| NA13509:HTT | 79 | 15-15 | 66-99 | EXPANDED | . | . | . | . | . | . | . | . | . | 1.848827406791674e-12 | 39.79309175778684 | EXPANDED | 4.843514e-05 | EXPANDED |
| NA13515:HTT | 68 | 16-16 | 60-83 | EXPANDED | 18 | 18 | normal | 30 | 27 | normal | 16 | 0 | normal | 1.7725548029144713e-23 | 59.179412621242406 | EXPANDED | 1.695230e-04 | EXPANDED |
| NA13507:HTT | 46 | 15-15 | 46-55 | EXPANDED | 17 | 17 | normal | . | . | . | 15 | 0 | normal | 7.010672527459866e-11 | 37.883519932305724 | EXPANDED | 8.072524e-06 | EXPANDED |
| NA13508:HTT | 61 | 22-22 | 54-78 | EXPANDED | 24 | 24 | normal | 33 | 33 | normal | 22 | 0 | normal | 1.1859624604436796e-06 | 32.67266371319075 | EXPANDED | 8.072524e-06 | EXPANDED |
| NA13510:HTT | 44 | 15-15 | 44-57 | EXPANDED | 17 | 17 | normal | . | . | . | 15 | 0 | normal | 2.4361265326230777e-11 | 38.32357016992598 | EXPANDED | 8.072524e-06 | EXPANDED |
| NA13511:HTT | 47 | 5-5 | 47-56 | EXPANDED | . | . | . | . | . | . | . | . | . | 4.678760756472997e-23 | 58.12089595957269 | EXPANDED | 8.072524e-06 | EXPANDED |
| NA13512:HTT | 44 | 16-16 | 44-54 | EXPANDED | 18 | 18 | normal | . | . | . | 16 | 0 | normal | 6.294465213895462e-06 | 31.76895463863977 | EXPANDED | 8.072524e-06 | EXPANDED |
| NA13513:HTT | 59 | 15-15 | 54-71 | EXPANDED | 17 | 17 | normal | . | . | . | 15 | 0 | normal | 1.77253367768293e-23 | 59.179412621242406 | EXPANDED | 8.072524e-06 | EXPANDED |
| NA13514:HTT | 56 | 15-15 | 51-68 | EXPANDED | . | . | . | . | . | . | . | . | . | 2.0138340846723294e-08 | 34.64936393774704 | EXPANDED | 9.364127e-04 | EXPANDED |
| NA13503:HTT | 43 | 17-17 | 43-53 | EXPANDED | 22 | 22 | normal | . | . | . | . | . | . | 4.654634605179165e-15 | 43.461609572066756 | EXPANDED | 4.036262e-05 | EXPANDED |
| NA13504:HTT | 40 | 16-16 | 40-64 | EXPANDED | 18 | 18 | normal | 30 | 27 | normal | 16 | 0 | normal | 0.00034028101454248994 | 29.83741359552382 | EXPANDED | 8.072524e-06 | EXPANDED |
| NA13505:HTT | 52 | 22-22 | 44-60 | EXPANDED | 24 | 24 | normal | 33 | 33 | normal | . | . | . | 3.2194201976482876e-10 | 36.957130619087295 | EXPANDED | 8.072524e-06 | EXPANDED |
| NA13506:HTT | 46 | 17-17 | 46-55 | EXPANDED | 20 | 20 | normal | . | . | . | . | . | . | 1.8611334182995864e-13 | 41.53480631786409 | EXPANDED | 2.421757e-05 | EXPANDED |
| NA06926:ATXN1 | 36 | 29-29 | 36-68 | normal | 30 | 30 | normal | . | . | . | . | . | . | 2.6723432691580394e-16 | 55.28991375976906 | EXPANDED | 8.072524e-06 | EXPANDED |
| NA13536:ATXN1 | 37 | 31-31 | 37-37 | normal | 32 | 32 | normal | 32 | 32 | normal | 31 | 0 | normal | 6.65557331165976e-11 | 46.883519932305724 | EXPANDED | 2.139219e-03 | EXPANDED |
| NA13537:ATXN1 | 56 | 43-54 | 52-66 | EXPANDED | . | . | . | 30 | 30 | normal | . | . | . | 1.8805157226315128e-19 | 60.6707761522461 | EXPANDED | 8.072524e-06 | EXPANDED |
| NA06151:ATXN3 | 55 | 18-18 | 44-65 | normal | 24 | 24 | normal | 23 | 29 | normal | 18 | 0 | normal | 2.268972274384911e-11 | 49.00200868905826 | EXPANDED | . | . |
| NA23709:AR | 46 | 46-54 | 46-54 | EXPANDED | . | . | . | . | . | . | . | . | . | 0.02496645229833929 | 40.00464435411614 | EXPANDED | 2.421757e-04 | EXPANDED |
| hg19C9ORF721000:C9ORF72 | 99 | 3-3 | 80-124 | EXPANDED | 10 | 10 | normal | 10 | 10 | normal | . | . | . | 0.00072 | 32.2 | EXPANDED | 4.848328e-04 | EXPANDED |
| hg19C9ORF72500:C9ORF72 | 119 | 3-3 | 97-148 | EXPANDED | 10 | 10 | normal | . | . | . | . | . | . | 4.2e-05 | 39.6 | EXPANDED | 4.848328e-04 | EXPANDED |
| hg19C9ORF7260:C9ORF72 | 71 | 3-3 | 60-94 | EXPANDED | 10 | 10 | normal | . | . | . | . | . | . | 6.7e-09 | 65.2 | EXPANDED | 5.817994e-04 | EXPANDED |
| hg19FMR21000:FMR2 | 15 | 15-15 | 15-15 | normal | . | . | . | . | . | . | . | . | . | 3e-32 | 153.1 | EXPANDED | 8.072524e-06 | EXPANDED |
| hg19FMR2200:FMR2 | 15 | 15-15 | 15-15 | normal | . | . | . | . | . | . | . | . | . | 8.6e-33 | 156.1 | EXPANDED | 8.072524e-06 | EXPANDED |
| hg19FMR2500:FMR2 | 15 | 15-15 | 15-15 | normal | . | . | . | . | . | . | . | . | . | 3e-32 | 153.1 | EXPANDED | 8.072524e-06 | EXPANDED |

**SUPPLEMENTARY TABLE 6.** Allelic calls from ExpansionHunter versions 2 and 3 (EH_v2 and EH_v3), GangSTR, TREDPARSE (TRED), lobSTR, HipSTR, RepeatSeq, STRetch, and exSTRa with (wctrls) and without controls (woctrls) STR detection tools for the EGA and simulated GS data (with BWA alignment). For each tool, "a1" and "a2" indicate the calls for allele 1 and allele 2, respectively, while CI indicates the confidence interval. For status, "EXPANDED" indicates at least one allele was reported to be over the full mutation threshold, while "normal" indicates that both alleles were reported to be under the threshold. For STRetch, both the p-value and genotyped repeat counts (repeatUnit) are reported. The p-status for STRetch and exSTRa are marked as "EXPANDED" for those samples with p-values <0.05, and "normal" for those samples with p-values >0.05. For all tools except STRetch and exSTRa, empty cells indicate a call was not made for this sample in the corresponding STR locus.

| sampleID:gene | EH_v2:a1 | EH_v2:a2 | EH_v2:CI-a1 | EH_v2:CI-a2 | EH_v2:status | EH_v3:a1 | EH_v3:a2 | EH_v3:CI-a1 | EH_v3:CI-a2 | EH_v3:status | GangSTR:a1 | GangSTR:a2 | GangSTR:CI-a1 | GangSTR:CI-a2 | GangSTR:status | TRED:a1 | TRED:a2 | TRED:CI-a1 | TRED:CI-a2 | TRED:status | lobSTR:a1 | lobSTR:a2 | lobSTR:status | HipSTR:a1 | HipSTR:a2 | HipSTR:status | RepeatSeq:a1 | RepeatSeq:a2 | RepeatSeq:status | STRetch:p-value | STRetch:repeatUnit | STRetch:pstatus | exSTRa(wctrls):p-value | exSTRa(wctrls):pstatus | exSTRa(woctrls):p-value | exSTRa(woctrls):pstatus |
|---|---|---|---|---|---|---|---|---|---|---|---|---|---|---|---|---|---|---|---|---|---|---|---|---|---|---|---|---|---|---|---|---|---|---|---|---|
| NA06075:DMPK | 12 | 70 | 12-12 | 57-83 | EXPANDED | 12 | 61 | 12-12 | 50-70 | EXPANDED | 12 | 64 | 12-12 | 53-78 | EXPANDED | 12 | 65 | 12-12 | 56-76 | EXPANDED | 12 | 12 | normal | . | . | . | 12 | 0 | normal | 3.7277720557467967e-41 | 108.34164576556 | EXPANDED | 9.443316e-06 | EXPANDED | 8.138153e-06 | EXPANDED |
| NA04567:DMPK | 21 | 139 | 21-21 | 114-163 | EXPANDED | 21 | 135 | 21-21 | 108-160 | EXPANDED | 21 | 237 | 21-21 | 212-261 | EXPANDED | 21 | 93 | 21-21 | 80-116 | EXPANDED | 21 | 21 | normal | 20 | 20 | normal | 21 | 0 | normal | 7.892057562359336e-47 | 131.09690768412807 | EXPANDED | . | . | . | . |
| NA05164:DMPK | 21 | 121 | 21-21 | 97-142 | EXPANDED | 21 | 106 | 21-21 | 82-127 | EXPANDED | 21 | 203 | 21-21 | 186-227 | EXPANDED | 21 | 83 | 21-21 | 72-103 | EXPANDED | 21 | 21 | normal | 20 | 20 | normal | 21 | 20 | normal | 3.544050479922964e-35 | 88.81836861417233 | EXPANDED | 9.443316e-06 | EXPANDED | 8.138153e-06 | EXPANDED |
| NA04648:DMPK | 5 | 163 | 5-5 | 136-190 | EXPANDED | 5 | 139 | 5-5 | 113-162 | EXPANDED | 5 | 340 | 5-5 | 312-359 | EXPANDED | 5 | 100 | 5-5 | 85-125 | EXPANDED | 5 | 5 | normal | . | . | . | 5 | 0 | normal | 3.5049462133950315e-49 | 142.5357219746143 | EXPANDED | 9.443316e-06 | EXPANDED | 8.138153e-06 | EXPANDED |
| NA05152:DMPK | 5 | 162 | 5-5 | 120-204 | EXPANDED | 5 | 141 | 5-5 | 104-178 | EXPANDED | 666 | 850 | 666-666 | 784-926 | EXPANDED | 5 | 94 | 5-5 | 74-132 | EXPANDED | 5 | 5 | normal | . | . | . | . | . | . | 3.678098712788432e-42 | 112.76768911699213 | EXPANDED | 9.443316e-06 | EXPANDED | 8.138153e-06 | EXPANDED |
| NA23378:DMPK | 22 | 125 | 22-22 | 103-148 | EXPANDED | 22 | 131 | 22-22 | 102-157 | EXPANDED | 22 | 110 | 21-30 | 88-129 | EXPANDED | 22 | 94 | 22-22 | 80-116 | EXPANDED | 22 | 22 | normal | . | . | . | 22 | 0 | normal | 1.1734388904983037e-41 | 110.86114291198635 | EXPANDED | 9.443316e-06 | EXPANDED | 8.138153e-06 | EXPANDED |
| NA23374:DMPK | 5 | 132 | 5-5 | 108-156 | EXPANDED | 5 | 128 | 5-5 | 100-153 | EXPANDED | 5 | 188 | 5-5 | 167-210 | EXPANDED | 5 | 97 | 5-5 | 78-117 | EXPANDED | 5 | 5 | normal | . | . | . | 5 | 0 | normal | 1.3769131465391494e-42 | 113.246610958481 | EXPANDED | 9.443316e-06 | EXPANDED | 8.138153e-06 | EXPANDED |
| NA23300:DMPK | 5 | 129 | 5-5 | 106-152 | EXPANDED | 5 | 135 | 5-5 | 107-161 | EXPANDED | 5 | 27 | 5-5 | 16-32 | normal | 5 | 93 | 5-5 | 78-118 | EXPANDED | 5 | 5 | normal | 19 | 4 | normal | 5 | 0 | normal | 5.512189571954157e-36 | 90.77661613272883 | EXPANDED | 9.443316e-06 | EXPANDED | 8.138153e-06 | EXPANDED |
| NA03986:DMPK | 12 | 110 | 12-12 | 90-130 | EXPANDED | 12 | 111 | 12-12 | 88-133 | EXPANDED | 12 | 208 | 12-12 | 180-228 | EXPANDED | 12 | 80 | 12-12 | 70-99 | EXPANDED | 12 | 12 | normal | . | . | . | 12 | 0 | normal | 2.3628129568746877e-35 | 88.81836861417233 | EXPANDED | 9.443316e-06 | EXPANDED | 8.138153e-06 | EXPANDED |
| NA03989:DMPK | 13 | 153 | 13-13 | 126-178 | EXPANDED | 13 | 130 | 13-13 | 104-154 | EXPANDED | 13 | 1103 | 13-13 | 1059-1152 | EXPANDED | 13 | 113 | 13-13 | 94-140 | EXPANDED | 13 | 13 | normal | . | . | . | 13 | 0 | normal | 4.970730518608572e-43 | 116.12357307491801 | EXPANDED | 9.443316e-06 | EXPANDED | 8.138153e-06 | EXPANDED |
| NA03990:DMPK | 14 | 100 | 14-14 | 81-119 | EXPANDED | 14 | 85 | 14-14 | 67-100 | EXPANDED | 14 | 72 | 14-14 | 61-84 | EXPANDED | 14 | 67 | 14-14 | 59-80 | EXPANDED | 14 | 14 | normal | . | . | . | . | . | . | 3.353893516338058e-46 | 129.13981546383863 | EXPANDED | 9.443316e-06 | EXPANDED | 8.138153e-06 | EXPANDED |
| NA03696:DMPK | 12 | 131 | 12-12 | 106-153 | EXPANDED | 12 | 134 | 12-12 | 108-159 | EXPANDED | 12 | 258 | 12-12 | 237-281 | EXPANDED | 12 | 101 | 12-12 | 85-125 | EXPANDED | 12 | 12 | normal | 11 | 24 | normal | 12 | 0 | normal | 9.630077613467448e-35 | 86.96491278295703 | EXPANDED | 9.443316e-06 | EXPANDED | 8.138153e-06 | EXPANDED |
| NA03759:DMPK | 14 | 118 | 14-14 | 96-138 | EXPANDED | 14 | 116 | 14-14 | 92-138 | EXPANDED | 14 | 666 | 14-14 | 666-666 | EXPANDED | 14 | 95 | 14-14 | 81-125 | EXPANDED | 14 | 14 | normal | 19 | 13 | normal | 14 | 0 | normal | 2.8804922967648865e-40 | 105.7526764442294 | EXPANDED | 9.443316e-06 | EXPANDED | 8.138153e-06 | EXPANDED |
| NA04034:DMPK | 12 | 113 | 12-12 | 92-134 | EXPANDED | 12 | 111 | 12-12 | 84-135 | EXPANDED | 12 | 260 | 12-12 | 238-282 | EXPANDED | 12 | 88 | 12-12 | 73-115 | EXPANDED | 12 | 12 | normal | . | . | . | 12 | 0 | normal | 1.2054130082764893e-37 | 97.69835776214023 | EXPANDED | 9.443316e-06 | EXPANDED | 8.138153e-06 | EXPANDED |
| NA03697:DMPK | 12 | 134 | 12-12 | 110-156 | EXPANDED | 12 | 131 | 12-12 | 105-155 | EXPANDED | 12 | 204 | 12-12 | 186-227 | EXPANDED | 12 | 111 | 12-12 | 92-139 | EXPANDED | 12 | 12 | normal | 19 | 11 | normal | 12 | 0 | normal | 5.467350112003751e-44 | 118.72233913960766 | EXPANDED | 9.443316e-06 | EXPANDED | 8.138153e-06 | EXPANDED |
| NA03132:DMPK | 5 | 115 | 5-5 | 93-135 | EXPANDED | 5 | 90 | 5-5 | 71-106 | EXPANDED | 5 | 666 | 5-37 | 646-666 | EXPANDED | 5 | 90 | 5-5 | 77-114 | EXPANDED | 5 | 5 | normal | . | . | . | 5 | 0 | normal | 1.3845178527636742e-38 | 99.85652117365396 | EXPANDED | 9.443316e-06 | EXPANDED | 8.138153e-06 | EXPANDED |
| NA03756:DMPK | 13 | 134 | 13-13 | 110-156 | EXPANDED | 13 | 126 | 13-13 | 102-151 | EXPANDED | 13 | 200 | 13-13 | 180-223 | EXPANDED | 13 | 94 | 13-13 | 81-116 | EXPANDED | 13 | 13 | normal | . | . | . | 13 | 0 | normal | 1.8310962160517065e-36 | 92.84874494897238 | EXPANDED | 9.443316e-06 | EXPANDED | 8.138153e-06 | EXPANDED |
| NA13716:ATN1 | 20 | 62 | 20-20 | 52-71 | EXPANDED | 20 | 58 | 20-20 | 50-67 | EXPANDED | 16 | 62 | 16-16 | 55-72 | EXPANDED | 16 | 58 | 16-16 | 53-70 | EXPANDED | 20 | 20 | normal | . | . | . | 16 | 0 | normal | 1.6582902531602304e-22 | 54.70200868905826 | EXPANDED | 9.443316e-06 | EXPANDED | 8.072524e-06 | EXPANDED |
| NA13717:ATN1 | 19 | 67 | 19-19 | 56-77 | EXPANDED | 19 | 56 | 19-19 | 50-62 | EXPANDED | 15 | 61 | 15-15 | 52-71 | EXPANDED | 15 | 58 | 15-15 | 53-67 | EXPANDED | 19 | 19 | normal | . | . | . | 15 | 0 | normal | 7.401094261799903e-25 | 60.27639850519684 | EXPANDED | 9.443316e-06 | EXPANDED | 8.072524e-06 | EXPANDED |
| NA03816:FXN | 96 | 144 | 50-112 | 79-169 | EXPANDED | 84 | 128 | 50-118 | 77-180 | EXPANDED | 1 | 1 | 1-1 | 1-1 | normal | 43 | 158 | 43-111 | 101-184 | EXPANDED | . | . | . | . | . | . | . | . | . | 2.9832369461813087e-05 | 15.578422974583995 | EXPANDED | 9.443316e-06 | EXPANDED | 8.072524e-06 | EXPANDED |
| NA04079:FXN | 97 | 144 | 50-113 | 79-168 | EXPANDED | 93 | 148 | 50-136 | 86-210 | EXPANDED | 16 | 16 | 12-16 | 12-16 | normal | 46 | 187 | 47-126 | 115-215 | EXPANDED | . | . | . | . | . | . | . | . | . | 1.1833946335987388e-05 | 16.198337727475767 | EXPANDED | 9.443316e-06 | EXPANDED | 8.072524e-06 | EXPANDED |
| NA14519:FXN | 9 | 84 | 9-9 | 70-98 | EXPANDED | 9 | 82 | 9-9 | 64-99 | EXPANDED | 1 | 1 | 1-5 | 1-5 | normal | 9 | 77 | 9-9 | 66-97 | EXPANDED | 10 | 10 | normal | . | . | . | . | . | . | 0.7401573219255145 | 10.407802243504225 | normal | 9.443316e-06 | EXPANDED | 8.072524e-06 | EXPANDED |
| NA15850:FXN | 109 | 171 | 50-127 | 90-199 | EXPANDED | 87 | 135 | 50-124 | 81-189 | EXPANDED | 25 | 25 | 23-25 | 23-25 | normal | 45 | 170 | 46-116 | 107-194 | EXPANDED | . | . | . | . | . | . | . | . | . | 0.7401099148180085 | 10.407802243504225 | normal | 9.443316e-06 | EXPANDED | 8.072524e-06 | EXPANDED |
| NA15847:FXN | 9 | 109 | 9-9 | 88-128 | EXPANDED | 9 | 120 | 9-9 | 91-147 | EXPANDED | 8 | 11 | 3-12 | 3-12 | normal | 9 | 120 | 9-9 | 95-157 | EXPANDED | 10 | 10 | normal | . | . | . | 9 | 6 | normal | 8.995220715831744e-07 | 17.372663713190747 | EXPANDED | 9.443316e-06 | EXPANDED | 8.072524e-06 | EXPANDED |
| NA15848:FXN | 8 | 101 | 8-8 | 81-119 | EXPANDED | 8 | 106 | 8-8 | 82-128 | EXPANDED | 8 | 18 | 8-8 | 8-21 | normal | 8 | 94 | 8-8 | 79-118 | EXPANDED | 8 | 8 | normal | 20 | 20 | normal | . | . | . | 7.12961035007329e-06 | 16.468954638639772 | EXPANDED | 9.443316e-06 | EXPANDED | 8.072524e-06 | EXPANDED |
| NA16197:FXN | 110 | 173 | 50-130 | 91-200 | EXPANDED | 90 | 144 | 50-131 | 87-202 | EXPANDED | 1 | 1 | 1-1 | 1-1 | normal | 51 | 187 | 47-125 | 116-212 | EXPANDED | . | . | . | . | . | . | . | . | . | 0.00041235062995444654 | 14.53741359552382 | EXPANDED | 9.443316e-06 | EXPANDED | 8.072524e-06 | EXPANDED |
| NA16200:FXN | 9 | 100 | 9-9 | 82-118 | EXPANDED | 9 | 95 | 9-9 | 73-114 | EXPANDED | 7 | 20 | 1-20 | 2-21 | normal | 9 | 86 | 9-9 | 73-109 | EXPANDED | 10 | 10 | normal | 21 | 21 | normal | . | . | . | 0.0009865965957953724 | 14.096233139666829 | EXPANDED | 9.443316e-06 | EXPANDED | 8.072524e-06 | EXPANDED |
| NA16202:FXN | 8 | 89 | 8-8 | 74-105 | EXPANDED | 8 | 101 | 8-8 | 80-122 | EXPANDED | 8 | 8 | 5-8 | 5-8 | normal | 8 | 84 | 8-8 | 72-107 | EXPANDED | . | . | . | 18 | 20 | normal | . | . | . | 2.003949716253737e-07 | 18.315063337850766 | EXPANDED | 9.443316e-06 | EXPANDED | 8.072524e-06 | EXPANDED |
| NA16203:FXN | 22 | 166 | 22-22 | 139-193 | EXPANDED | 96 | 158 | 50-142 | 93-224 | EXPANDED | 19 | 19 | 1-19 | 1-19 | normal | 22 | 218 | 22-22 | 182-264 | EXPANDED | . | . | . | . | . | . | . | . | . | 0.20362351863908823 | 11.860705206113849 | normal | 9.443316e-06 | EXPANDED | 8.072524e-06 | EXPANDED |
| NA16205:FXN | 96 | 143 | 50-112 | 79-166 | EXPANDED | 90 | 140 | 50-130 | 83-198 | EXPANDED | 25 | 25 | 17-25 | 17-25 | normal | 51 | 168 | 46-116 | 106-194 | EXPANDED | . | . | . | . | . | . | . | . | . | 0.0003829426595419 | 14.53741359552382 | EXPANDED | 9.443316e-06 | EXPANDED | 8.072524e-06 | EXPANDED |
| NA16209:FXN | 102 | 155 | 50-120 | 85-180 | EXPANDED | 91 | 146 | 50-133 | 86-207 | EXPANDED | 6 | 6 | 4-6 | 4-6 | normal | 46 | 176 | 47-120 | 110-201 | EXPANDED | . | . | . | . | . | . | 6 | 0 | normal | 0.000324955901415001 | 14.53741359552382 | EXPANDED | 9.443316e-06 | EXPANDED | 8.072524e-06 | EXPANDED |
| NA16210:FXN | 111 | 172 | 50-130 | 91-199 | EXPANDED | 93 | 149 | 50-136 | 88-211 | EXPANDED | 8 | 8 | 1-8 | 1-8 | normal | 47 | 195 | 48-129 | 119-219 | EXPANDED | . | . | . | . | . | . | . | . | . | 0.09558598632894726 | 12.012331779984216 | normal | 9.443316e-06 | EXPANDED | 8.072524e-06 | EXPANDED |
| NA16212:FXN | 8 | 112 | 8-8 | 92-132 | EXPANDED | 8 | 123 | 8-8 | 96-147 | EXPANDED | 8 | 8 | 5-8 | 5-8 | normal | 8 | 110 | 8-8 | 92-138 | EXPANDED | 8 | 9 | normal | . | . | . | . | . | . | 0.04898181202316149 | 12.339956149206124 | EXPANDED | 9.443316e-06 | EXPANDED | 8.072524e-06 | EXPANDED |
| NA16216:FXN | 114 | 181 | 50-135 | 92-211 | EXPANDED | 21 | 182 | 21-21 | 144-217 | EXPANDED | 15 | 15 | 14-15 | 14-15 | normal | 51 | 208 | 48-137 | 126-237 | EXPANDED | . | . | . | . | . | . | 6 | 0 | normal | 5.549538395112172e-13 | 25.039922789984015 | EXPANDED | 9.443316e-06 | EXPANDED | 8.072524e-06 | EXPANDED |
| NA16213:FXN | 8 | 93 | 8-8 | 75-110 | EXPANDED | 8 | 106 | 8-8 | 82-131 | EXPANDED | 8 | 21 | 8-8 | 14-25 | normal | 8 | 90 | 8-8 | 75-118 | EXPANDED | 8 | 8 | normal | . | . | . | 6 | 8 | normal | 0.026968005133449918 | 12.704644354116141 | EXPANDED | 9.443316e-06 | EXPANDED | 8.072524e-06 | EXPANDED |
| NA16215:FXN | 78 | 107 | 50-92 | 63-127 | EXPANDED | 27 | 134 | 27-27 | 101-164 | EXPANDED | 28 | 28 | 23-28 | 23-28 | normal | 27 | 116 | 27-27 | 91-151 | EXPANDED | . | . | . | . | . | . | . | . | . | 0.001931481583624665 | 13.573372575142738 | EXPANDED | 9.443316e-06 | EXPANDED | 8.072524e-06 | EXPANDED |
| NA16214:FXN | 98 | 147 | 50-117 | 79-172 | EXPANDED | 89 | 138 | 50-128 | 82-194 | EXPANDED | 25 | 25 | 15-25 | 15-25 | normal | 45 | 177 | 46-120 | 110-203 | EXPANDED | . | . | . | . | . | . | . | . | . | 0.03477864273660233 | 12.517287231386582 | EXPANDED | 9.443316e-06 | EXPANDED | 8.072524e-06 | EXPANDED |
| NA16227:FXN | 107 | 164 | 50-126 | 85-192 | EXPANDED | 94 | 153 | 50-138 | 88-218 | EXPANDED | 25 | 25 | 16-25 | 16-25 | normal | 44 | 178 | 45-121 | 111-205 | EXPANDED | . | . | . | . | . | . | 6 | 0 | normal | 4.832109811887195e-09 | 19.73312766164753 | EXPANDED | 9.443316e-06 | EXPANDED | 8.072524e-06 | EXPANDED |
| NA16229:FXN | 19 | 95 | 19-19 | 78-112 | EXPANDED | 19 | 90 | 19-19 | 71-109 | EXPANDED | 8 | 21 | 1-21 | 8-23 | normal | 19 | 84 | 19-19 | 72-108 | EXPANDED | 20 | 20 | normal | . | . | . | 6 | 6 | normal | 0.7837364129909655 | 10.168047022663163 | normal | 9.443316e-06 | EXPANDED | 8.072524e-06 | EXPANDED |
| NA16228:FXN | 6 | 188 | 6-6 | 157-220 | EXPANDED | 103 | 174 | 50-156 | 99-249 | EXPANDED | 22 | 22 | 18-22 | 18-22 | normal | 51 | 220 | 48-143 | 132-248 | EXPANDED | . | . | . | . | . | . | 6 | 0 | normal | 1.9338354731690887e-08 | 19.349363937747043 | EXPANDED | 9.443316e-06 | EXPANDED | 8.072524e-06 | EXPANDED |
| NA16237:FXN | 8 | 83 | 8-8 | 68-99 | EXPANDED | 8 | 76 | 8-8 | 59-91 | EXPANDED | 8 | 19 | 8-8 | 16-21 | normal | 8 | 79 | 8-8 | 66-100 | EXPANDED | 8 | 8 | normal | . | . | . | 8 | 6 | normal | 1.3475567782712614e-05 | 15.87882609688239 | EXPANDED | 9.443316e-06 | EXPANDED | 8.072524e-06 | EXPANDED |
| NA16243:FXN | 114 | 178 | 50-135 | 93-207 | EXPANDED | 98 | 160 | 50-147 | 93-228 | EXPANDED | 20 | 20 | 1-20 | 1-20 | normal | 47 | 207 | 49-136 | 125-233 | EXPANDED | . | . | . | . | . | . | . | . | . | 1.784802306823897e-08 | 19.349363937747043 | EXPANDED | 9.443316e-06 | EXPANDED | 8.072524e-06 | EXPANDED |
| NA16240:FXN | 9 | 110 | 9-9 | 89-131 | EXPANDED | 9 | 102 | 9-9 | 79-122 | EXPANDED | 6 | 11 | 6-12 | 6-12 | normal | 9 | 90 | 9-9 | 77-112 | EXPANDED | 9 | 10 | normal | 21 | 21 | normal | . | . | . | 1.9304783525437763e-06 | 17.054861688424708 | EXPANDED | 9.443316e-06 | EXPANDED | 8.072524e-06 | EXPANDED |
| NA16207:FXN | 8 | 180 | 8-8 | 151-208 | EXPANDED | 95 | 155 | 50-140 | 93-218 | EXPANDED | 1 | 1 | 1-1 | 1-1 | normal | 51 | 203 | 47-133 | 124-229 | EXPANDED | . | . | . | . | . | . | . | . | . | 9.581449663113758e-11 | 22.10723736424742 | EXPANDED | 9.443316e-06 | EXPANDED | 8.072524e-06 | EXPANDED |
| NA06895:FMR1 | 23 | 0 | 23-23 | 0-0 | normal | 23 | 0 | 23-23 | 0-0 | normal | 21 | 0 | 12-22 | 0-0 | normal | 23 | 23 | 23-23 | 23-23 | normal | 28 | 28 | normal | 23 | 23 | normal | . | . | . | . | . | . | . | . | . | . |
| NA04025:FMR1 | 101 | 0 | 83-118 | 0-0 | normal | 79 | 0 | 62-94 | 0-0 | normal | 23 | 0 | 10-23 | 0-0 | normal | 51 | 76 | 44-72 | 61-104 | normal | . | . | . | . | . | . | . | . | . | 0.5902410425783222 | 39.82730357396629 | normal | 9.443316e-06 | EXPANDED | . | . |
| NA04926:FMR1 | 117 | 0 | 95-137 | 0-0 | normal | 110 | 0 | 85-134 | 0-0 | normal | 17 | 0 | 9-17 | 0-0 | normal | 38 | 77 | 38-73 | 61-110 | normal | . | . | . | . | . | . | . | . | . | 0.22010201737603807 | 43.732265042149834 | normal | 9.443316e-06 | EXPANDED | 2.421757e-05 | EXPANDED |
| NA05131:FMR1 | 164 | 0 | 135-192 | 0-0 | normal | 90 | 0 | 67-110 | 0-0 | normal | 39 | 0 | 27-39 | 0-0 | normal | 43 | 149 | 44-114 | 93-195 | normal | . | . | . | . | . | . | . | . | . | 0.015216551830441167 | 52.27418861391992 | EXPANDED | 9.443316e-06 | EXPANDED | 8.072524e-06 | EXPANDED |
| NA05185:FMR1 | 78 | 0 | 63-91 | 0-0 | normal | 74 | 0 | 58-88 | 0-0 | normal | 38 | 0 | 19-38 | 0-0 | normal | 39 | 66 | 39-66 | 56-96 | normal | . | . | . | . | . | . | . | . | . | . | . | . | 9.443316e-06 | EXPANDED | 2.163436e-03 | EXPANDED |
| NA09145:FMR1 | 99 | 0 | 81-116 | 0-0 | normal | 87 | 0 | 67-105 | 0-0 | normal | 34 | 0 | 18-34 | 0-0 | normal | 35 | 86 | 35-76 | 65-121 | normal | . | . | . | . | . | . | . | . | . | 0.03163493537181473 | 48.98344508707048 | EXPANDED | 9.443316e-06 | EXPANDED | 1.614505e-05 | EXPANDED |
| NA09237:FMR1 | 94 | 0 | 76-110 | 0-0 | normal | 86 | 0 | 67-103 | 0-0 | normal | 30 | 0 | 24-30 | 0-0 | normal | 31 | 79 | 31-70 | 62-112 | normal | . | . | . | . | . | . | . | . | . | 0.7564862689737221 | 33.77620843200008 | normal | 9.443316e-06 | EXPANDED | 1.614505e-05 | EXPANDED |
| NA07063:FMR1 | 32 | 88 | 32-32 | 76-100 | normal | 32 | 79 | 32-32 | 66-91 | normal | 32 | 36 | 24-36 | 24-36 | normal | 32 | 59 | 32-32 | 54-68 | normal | . | . | . | . | . | . | . | . | . | 0.9988193918748451 | 31.590980402100367 | normal | 9.443316e-06 | EXPANDED | . | . |
| NA07539:FMR1 | 23 | 0 | 23-23 | 0-0 | normal | 23 | 0 | 23-23 | 0-0 | normal | 23 | 0 | 16-23 | 0-0 | normal | 4 | 23 | 4-4 | 23-23 | normal | . | . | . | 23 | 23 | normal | . | . | . | . | . | . | . | . | . | . |
| NA06890:FMR1 | 30 | 0 | 30-30 | 0-0 | normal | 30 | 0 | 30-30 | 0-0 | normal | 29 | 0 | 22-30 | 0-0 | normal | 30 | 30 | 30-30 | 30-30 | normal | . | . | . | . | . | . | . | . | . | 0.5077802907884078 | 37.846232390972816 | normal | . | . | . | . |
| NA06905:FMR1 | 23 | 56 | 23-23 | 50-62 | normal | 23 | 60 | 23-23 | 50-68 | normal | 23 | 30 | 17-30 | 17-31 | normal | 39 | 55 | 39-54 | 40-67 | normal | 28 | 28 | normal | . | . | . | . | . | . | 0.7078028638195785 | 29.538138645989306 | normal | 7.837953e-04 | EXPANDED | . | . |
| NA07536:FMR1 | 23 | 0 | 23-23 | 0-0 | normal | 23 | 0 | 23-23 | 0-0 | normal | 23 | 0 | 23-24 | 0-0 | normal | 23 | 23 | 23-23 | 23-23 | normal | 28 | 28 | normal | 23 | 23 | normal | 23 | 0 | normal | 0.7556262468044473 | 31.517287231386582 | normal | . | . | . | . |
| NA07540:FMR1 | 23 | 29 | 23-23 | 29-29 | normal | 23 | 29 | 23-23 | 29-29 | normal | 23 | 29 | 22-23 | 29-29 | normal | 23 | 29 | 23-23 | 29-29 | normal | 28 | 34 | normal | 23 | 29 | normal | 29 | 0 | normal | . | . | . | . | . | . | . |
| NA07542:FMR1 | 23 | 0 | 23-23 | 0-0 | normal | 23 | 0 | 23-23 | 0-0 | normal | 23 | 0 | 23-23 | 0-0 | normal | 23 | 23 | 23-23 | 23-23 | normal | 28 | 28 | normal | 23 | 23 | normal | . | . | . | . | . | . | . | . | . | . |
| NA06910:FMR1 | 30 | 71 | 30-30 | 58-81 | normal | 30 | 76 | 30-30 | 58-91 | normal | 32 | 32 | 27-35 | 27-35 | normal | 30 | 38 | 30-30 | 38-62 | normal | . | . | . | . | . | . | . | . | . | 0.6689549129386576 | 29.822937443502347 | normal | 2.832995e-05 | EXPANDED | . | . |
| NA06894:FMR1 | 30 | 54 | 30-30 | 50-58 | normal | 31 | 63 | 31-31 | 50-74 | normal | 27 | 27 | 19-27 | 19-27 | normal | 26 | 30 | 26-30 | 30-56 | normal | . | . | . | . | . | . | . | . | . | 0.013941195829359163 | 52.27418861391992 | EXPANDED | . | . | . | . |
| NA07541:FMR1 | 29 | 31 | 29-29 | 31-31 | normal | 29 | 52 | 29-29 | 50-55 | normal | 29 | 29 | 12-29 | 12-29 | normal | 29 | 31 | 29-29 | 31-31 | normal | 12 | 12 | normal | . | . | . | . | . | . | . | . | . | . | . | . | . |
| NA07175:FMR1 | 23 | 30 | 23-23 | 30-30 | normal | 23 | 45 | 23-23 | 23-45 | normal | 23 | 24 | 20-25 | 20-25 | normal | 23 | 30 | 23-23 | 30-31 | normal | 28 | 28 | normal | 20 | 23 | normal | . | . | . | . | . | . | . | . | . | . |
| NA06889:FMR1 | 23 | 30 | 23-23 | 30-30 | normal | 23 | 30 | 23-23 | 30-30 | normal | 26 | 28 | 21-30 | 21-30 | normal | 23 | 30 | 23-23 | 30-30 | normal | 28 | 35 | normal | 20 | 23 | normal | . | . | . | . | . | . | . | . | . | . |
| NA06893:FMR1 | 23 | 30 | 23-23 | 30-30 | normal | 23 | 30 | 23-23 | 30-30 | normal | 23 | 24 | 23-23 | 23-26 | normal | 23 | 30 | 23-23 | 30-30 | normal | 28 | 28 | normal | . | . | . | . | . | . | . | . | . | . | . | . | . |
| NA06896:FMR1 | 23 | 79 | 23-23 | 64-92 | normal | 23 | 70 | 23-23 | 55-82 | normal | 23 | 38 | 23-23 | 23-40 | normal | 23 | 59 | 23-23 | 54-75 | normal | 28 | 28 | normal | 20 | 23 | normal | 23 | 20 | normal | . | . | . | 9.443316e-06 | EXPANDED | . | . |
| NA07538:FMR1 | 29 | 29 | 29-29 | 29-29 | normal | 29 | 52 | 29-29 | 50-55 | normal | 13 | 23 | 13-23 | 13-24 | normal | 29 | 29 | 24-29 | 29-29 | normal | . | . | . | . | . | . | . | . | . | . | . | . | . | . | . | . |
| NA07537:FMR1 | 29 | 79 | 29-29 | 64-91 | normal | 29 | 70 | 29-29 | 56-81 | normal | 29 | 61 | 29-54 | 54-71 | normal | 29 | 41 | 29-29 | 41-58 | normal | 34 | 34 | normal | 29 | 29 | normal | 29 | 22 | normal | 0.8163119020450283 | 37.64184839737077 | normal | 2.549695e-04 | EXPANDED | . | . |
| NA06897:FMR1 | 77 | 0 | 63-91 | 0-0 | normal | 68 | 0 | 52-81 | 0-0 | normal | 18 | 0 | 4-18 | 0-0 | normal | 51 | 65 | 45-68 | 56-95 | normal | . | . | . | . | . | . | . | . | . | 0.41415199905733374 | 37.846232390972816 | normal | 9.443316e-06 | EXPANDED | . | . |
| NA07174:FMR1 | 30 | 0 | 30-30 | 0-0 | normal | 52 | 0 | 50-55 | 0-0 | normal | 32 | 0 | 27-32 | 0-0 | normal | 31 | 31 | 31-50 | 31-71 | normal | . | . | . | . | . | . | . | . | . | . | . | . | . | . | . | . |
| NA06903:FMR1 | 24 | 59 | 24-24 | 50-67 | normal | 24 | 61 | 24-24 | 50-72 | normal | 27 | 30 | 23-30 | 23-30 | normal | 24 | 45 | 24-24 | 45-56 | normal | . | . | . | . | . | . | . | . | . | 0.5587675979859662 | 33.09623313966683 | normal | . | . | . | . |
| NA07543:FMR1 | 20 | 29 | 20-20 | 29-29 | normal | 20 | 29 | 20-20 | 29-29 | normal | 20 | 20 | 20-20 | 20-20 | normal | 20 | 29 | 20-20 | 20-29 | normal | 25 | 25 | normal | 20 | 20 | normal | 20 | 0 | normal | 0.41215555482648236 | 41.107237364247425 | normal | . | . | . | . |
| NA06852:FMR1 | 83 | 0 | 69-98 | 0-0 | normal | 75 | 0 | 59-88 | 0-0 | normal | 36 | 0 | 31-36 | 0-0 | normal | 42 | 74 | 42-71 | 59-104 | normal | . | . | . | . | . | . | . | . | . | . | . | . | 9.443316e-06 | EXPANDED | . | . |
| NA06891:FMR1 | 75 | 0 | 61-88 | 0-0 | normal | 82 | 0 | 59-101 | 0-0 | normal | 11 | 0 | 5-11 | 0-0 | normal | 37 | 56 | 37-61 | 42-84 | normal | . | . | . | . | . | . | . | . | . | 0.03218152699008422 | 48.595907134202605 | EXPANDED | 9.443316e-06 | EXPANDED | 2.421757e-05 | EXPANDED |
| NA06907:FMR1 | 29 | 65 | 29-29 | 54-74 | normal | 29 | 74 | 29-29 | 56-89 | normal | 29 | 32 | 21-32 | 21-32 | normal | 29 | 61 | 18-29 | 52-78 | normal | 23 | 23 | normal | . | . | . | . | . | . | 0.7379405686343744 | 36.37266371319075 | normal | 2.077530e-04 | EXPANDED | . | . |
| NA06906:FMR1 | 62 | 0 | 51-70 | 0-0 | normal | 64 | 0 | 52-74 | 0-0 | normal | 32 | 0 | 20-33 | 0-0 | normal | 35 | 64 | 35-63 | 55-91 | normal | . | . | . | . | . | . | . | . | . | 0.7087317570123631 | 29.538138645989306 | normal | 9.443316e-06 | EXPANDED | . | . |
| NA06892:FMR1 | 74 | 0 | 62-87 | 0-0 | normal | 65 | 0 | 52-75 | 0-0 | normal | 31 | 0 | 23-32 | 0-0 | normal | 60 | 60 | 43-67 | 58-93 | normal | . | . | . | . | . | . | . | . | . | 0.7375458024466894 | 36.37266371319075 | normal | 9.443316e-06 | EXPANDED | . | . |
| NA06904:FMR1 | 24 | 29 | 24-24 | 29-29 | normal | 29 | 46 | 29-29 | 29-46 | normal | 28 | 28 | 20-31 | 20-31 | normal | 29 | 29 | 29-29 | 29-51 | normal | . | . | . | . | . | . | . | . | . | . | . | . | . | . | . | . |
| NA06968:FMR1 | 33 | 75 | 33-33 | 62-87 | normal | 33 | 84 | 33-33 | 64-101 | normal | 26 | 52 | 24-53 | 33-57 | normal | 33 | 55 | 33-33 | 43-66 | normal | . | . | . | . | . | . | . | . | . | 0.7084976341740767 | 29.538138645989306 | normal | 9.443316e-06 | EXPANDED | . | . |
| NA07294:FMR1 | 97 | 0 | 79-116 | 0-0 | normal | 86 | 0 | 63-105 | 0-0 | normal | 24 | 0 | 6-24 | 0-0 | normal | 31 | 74 | 31-69 | 59-110 | normal | . | . | . | . | . | . | . | . | . | 0.7559035339336784 | 31.517287231386582 | normal | 9.443316e-06 | EXPANDED | . | . |
| NA09316:FMR1 | 90 | 0 | 74-106 | 0-0 | normal | 82 | 0 | 62-99 | 0-0 | normal | 27 | 0 | 20-27 | 0-0 | normal | 27 | 80 | 27-70 | 62-118 | normal | . | . | . | . | . | . | . | . | . | 0.7395014071948224 | 36.37266371319075 | normal | 9.443316e-06 | EXPANDED | 1.614505e-05 | EXPANDED |
| NA09317:FMR1 | 64 | 0 | 54-73 | 0-0 | normal | 63 | 0 | 50-74 | 0-0 | normal | 63 | 0 | 29-63 | 0-0 | normal | 42 | 42 | 42-42 | 42-42 | normal | . | . | . | . | . | . | . | . | . | 0.7833549061692991 | 29.168047022663163 | normal | 5.193824e-04 | EXPANDED | . | . |
| NA09497:FMR1 | 76 | 0 | 61-88 | 0-0 | normal | 71 | 0 | 55-84 | 0-0 | normal | 28 | 0 | 14-28 | 0-0 | normal | 43 | 61 | 43-65 | 53-89 | normal | . | . | . | . | . | . | . | . | . | 0.7846427392786426 | 35.75406620144316 | normal | 9.443316e-06 | EXPANDED | . | . |
| NA07730:FMR1 | 80 | 0 | 66-94 | 0-0 | normal | 86 | 0 | 66-102 | 0-0 | normal | 16 | 0 | 8-16 | 0-0 | normal | 36 | 72 | 36-68 | 59-100 | normal | . | . | . | . | . | . | . | . | . | 0.7837620406312714 | 29.168047022663163 | normal | 9.443316e-06 | EXPANDED | 2.986834e-04 | EXPANDED |
| NA03200:FMR1 | 154 | 0 | 126-180 | 0-0 | normal | 125 | 0 | 99-150 | 0-0 | normal | 18 | 0 | 10-18 | 0-0 | normal | 27 | 126 | 27-96 | 86-169 | normal | . | . | . | . | . | . | . | . | . | 0.707116475336365 | 39.82730357396629 | normal | 9.443316e-06 | EXPANDED | 8.072524e-06 | EXPANDED |
| NA20235:FMR1 | 29 | 40 | 29-29 | 40-40 | normal | 29 | 40 | 29-29 | 29-40 | normal | 29 | 29 | 27-32 | 27-32 | normal | 29 | 35 | 29-29 | 35-35 | normal | 34 | 34 | normal | 20 | 29 | normal | . | . | . | 0.07192203936589792 | 46.506946748334926 | normal | . | . | . | . |
| NA20238:FMR1 | 29 | 30 | 29-29 | 30-30 | normal | 29 | 52 | 29-29 | 50-54 | normal | 21 | 28 | 15-28 | 21-28 | normal | 29 | 30 | 29-29 | 30-30 | normal | . | . | . | 30 | 29 | normal | . | . | . | . | . | . | . | . | . | . |
| NA20237:FMR1 | 67 | 0 | 56-77 | 0-0 | normal | 68 | 0 | 56-79 | 0-0 | normal | 34 | 0 | 27-34 | 0-0 | normal | 32 | 32 | 32-53 | 33-70 | normal | . | . | . | . | . | . | . | . | . | 0.6903067804358576 | 34.578422974583994 | normal | 9.443316e-06 | EXPANDED | 1.453054e-04 | EXPANDED |
| NA20239:FMR1 | 20 | 82 | 20-20 | 67-97 | normal | 20 | 70 | 20-20 | 55-83 | normal | 20 | 57 | 20-20 | 32-67 | normal | 20 | 59 | 20-20 | 54-75 | normal | 25 | 25 | normal | 20 | 20 | normal | 20 | 0 | normal | 0.7845494239378982 | 31.339956149206124 | normal | 1.888663e-05 | EXPANDED | . | . |
| NA20242:FMR1 | 30 | 56 | 30-30 | 50-63 | normal | 30 | 63 | 30-30 | 50-74 | normal | 29 | 31 | 27-32 | 29-32 | normal | 30 | 45 | 30-30 | 45-60 | normal | . | . | . | . | . | . | . | . | . | 0.7390321520072598 | 36.37266371319075 | normal | 1.095425e-03 | EXPANDED | . | . |
| NA20243:FMR1 | 29 | 29 | 29-29 | 29-29 | normal | 29 | 31 | 29-29 | 29-38 | normal | 30 | 30 | 8-30 | 8-30 | normal | 29 | 31 | 29-29 | 31-31 | normal | . | . | . | . | . | . | . | . | . | . | . | . | . | . | . | . |
| NA20230:FMR1 | 58 | 0 | 50-64 | 0-0 | normal | 59 | 0 | 50-66 | 0-0 | normal | 38 | 0 | 35-38 | 0-0 | normal | 50 | 55 | 46-59 | 50-75 | normal | . | . | . | . | . | . | . | . | . | 0.7824676871531359 | 29.168047022663163 | normal | 9.443316e-06 | EXPANDED | 2.986834e-04 | EXPANDED |
| NA20232:FMR1 | 54 | 0 | 50-58 | 0-0 | normal | 55 | 0 | 50-60 | 0-0 | normal | 34 | 0 | 30-35 | 0-0 | normal | 39 | 55 | 39-59 | 42-79 | normal | . | . | . | . | . | . | . | . | . | 0.7393573059201957 | 31.70464435411614 | normal | 9.443316e-06 | EXPANDED | . | . |
| NA20233:FMR1 | 70 | 0 | 58-80 | 0-0 | normal | 82 | 0 | 61-99 | 0-0 | normal | 32 | 0 | 12-32 | 0-0 | normal | 9 | 73 | 9-9 | 59-109 | normal | . | . | . | . | . | . | . | . | . | 0.6900587874390874 | 29.67627776425815 | normal | 9.443316e-06 | EXPANDED | 1.243169e-03 | EXPANDED |
| NA20234:FMR1 | 31 | 54 | 31-31 | 50-58 | normal | 31 | 43 | 31-31 | 31-45 | normal | 19 | 19 | 18-22 | 18-22 | normal | 31 | 54 | 31-31 | 44-65 | normal | . | . | . | . | . | . | . | . | . | 0.7564499749711414 | 29.28462894716124 | normal | . | . | . | . |
| NA20236:FMR1 | 31 | 52 | 31-31 | 50-54 | normal | 52 | 55 | 50-55 | 50-60 | normal | 24 | 29 | 20-30 | 20-30 | normal | 37 | 37 | 37-37 | 37-37 | normal | . | . | . | . | . | . | . | . | . | 0.739991385567443 | 29.407802243504225 | normal | . | . | . | . |
| NA20231:FMR1 | 56 | 0 | 50-62 | 0-0 | normal | 69 | 0 | 53-81 | 0-0 | normal | 42 | 0 | 33-42 | 0-0 | normal | 40 | 62 | 40-65 | 54-93 | normal | . | . | . | . | . | . | . | . | . | 0.7839152240863776 | 33.537413595523816 | normal | . | . | . | . |
| NA20240:FMR1 | 58 | 70 | 50-65 | 50-81 | normal | 64 | 78 | 50-79 | 52-105 | normal | 30 | 30 | 12-30 | 12-30 | normal | 34 | 64 | 34-60 | 56-83 | normal | . | . | . | . | . | . | . | . | . | 0.6898122238452001 | 29.67627776425815 | normal | 9.443316e-06 | EXPANDED | . | . |
| NA20241:FMR1 | 29 | 74 | 29-29 | 61-87 | normal | 29 | 76 | 29-29 | 58-91 | normal | 32 | 32 | 28-32 | 28-32 | normal | 29 | 58 | 29-29 | 52-74 | normal | 34 | 34 | normal | 29 | 29 | normal | . | . | . | . | . | . | . | . | . | . |
| NA20244:FMR1 | 35 | 0 | 35-35 | 0-0 | normal | 52 | 0 | 50-55 | 0-0 | normal | 37 | 0 | 28-37 | 0-0 | normal | 37 | 37 | 37-37 | 37-37 | normal | . | . | . | . | . | . | . | . | . | 0.7092864134341791 | 29.538138645989306 | normal | . | . | . | . |
| NA07862:FMR1 | 82 | 0 | 68-97 | 0-0 | normal | 87 | 0 | 67-105 | 0-0 | normal | 20 | 0 | 10-20 | 0-0 | normal | 32 | 79 | 32-70 | 63-112 | normal | . | . | . | . | . | . | . | . | . | 0.7387311220150747 | 38.73312766164753 | normal | 9.443316e-06 | EXPANDED | 3.874811e-04 | EXPANDED |
| NA13509:HTT | 15 | 85 | 15-15 | 68-99 | EXPANDED | 15 | 79 | 15-15 | 60-95 | EXPANDED | 15 | 77 | 15-18 | 61-92 | EXPANDED | 15 | 77 | 15-15 | 65-94 | EXPANDED | . | . | . | . | . | . | 15 | 0 | normal | 1.848827406791674e-12 | 39.79309175778684 | EXPANDED | 9.443316e-06 | EXPANDED | 8.072524e-06 | EXPANDED |
| NA13515:HTT | 16 | 81 | 16-16 | 67-95 | EXPANDED | 16 | 75 | 16-16 | 60-87 | EXPANDED | 16 | 67 | 16-16 | 56-77 | EXPANDED | 16 | 66 | 16-16 | 58-78 | EXPANDED | 18 | 18 | normal | . | . | . | 16 | 0 | normal | 1.7725548029144713e-23 | 59.179412621242406 | EXPANDED | 9.443316e-06 | EXPANDED | 8.072524e-06 | EXPANDED |
| NA13507:HTT | 15 | 60 | 15-15 | 50-68 | EXPANDED | 15 | 56 | 15-15 | 50-63 | EXPANDED | 15 | 29 | 15-15 | 23-31 | normal | 15 | 46 | 15-15 | 46-54 | EXPANDED | 17 | 17 | normal | . | . | . | 15 | 0 | normal | 7.010672527459866e-11 | 37.883519932305724 | EXPANDED | 9.443316e-06 | EXPANDED | 8.072524e-06 | EXPANDED |
| NA13508:HTT | 22 | 62 | 22-22 | 52-71 | EXPANDED | 22 | 57 | 22-22 | 50-64 | EXPANDED | 22 | 61 | 22-22 | 54-71 | EXPANDED | 22 | 55 | 22-22 | 45-65 | EXPANDED | 24 | 24 | normal | 33 | 33 | normal | . | . | . | 1.1859624604436796e-06 | 32.67266371319075 | EXPANDED | 9.443316e-06 | EXPANDED | 8.072524e-06 | EXPANDED |
| NA13510:HTT | 15 | 58 | 15-15 | 50-64 | EXPANDED | 15 | 56 | 15-15 | 50-62 | EXPANDED | 15 | 33 | 15-15 | 26-36 | normal | 15 | 48 | 15-15 | 44-57 | EXPANDED | 17 | 17 | normal | . | . | . | 15 | 0 | normal | 2.4361265326230777e-11 | 38.32357016992598 | EXPANDED | 9.443316e-06 | EXPANDED | 8.072524e-06 | EXPANDED |
| NA13511:HTT | 43 | 56 | 43-43 | 50-62 | EXPANDED | 50 | 52 | 50-50 | 50-54 | EXPANDED | 42 | 42 | 5-43 | 5-43 | EXPANDED | 5 | 47 | 5-5 | 47-55 | EXPANDED | . | . | . | . | . | . | . | . | . | 4.678760756472997e-23 | 58.12089595957269 | EXPANDED | 9.443316e-06 | EXPANDED | 8.072524e-06 | EXPANDED |
| NA13512:HTT | 16 | 52 | 16-16 | 50-54 | EXPANDED | 16 | 43 | 16-16 | 16-50 | EXPANDED | 16 | 41 | 16-16 | 37-41 | EXPANDED | 16 | 44 | 16-16 | 44-52 | EXPANDED | 18 | 18 | normal | . | . | . | 16 | 0 | normal | 6.294465213895462e-06 | 31.76895463863977 | EXPANDED | 9.443316e-06 | EXPANDED | 8.072524e-06 | EXPANDED |
| NA13513:HTT | 15 | 62 | 15-15 | 52-70 | EXPANDED | 15 | 51 | 15-15 | 50-53 | EXPANDED | 15 | 58 | 15-15 | 52-65 | EXPANDED | 15 | 57 | 15-15 | 53-66 | EXPANDED | 17 | 17 | normal | . | . | . | 15 | 0 | normal | 1.77253367768293e-23 | 59.179412621242406 | EXPANDED | 9.443316e-06 | EXPANDED | 8.072524e-06 | EXPANDED |
| NA13514:HTT | 15 | 58 | 15-15 | 50-64 | EXPANDED | 15 | 52 | 15-15 | 50-54 | EXPANDED | 15 | 74 | 15-15 | 61-84 | EXPANDED | 15 | 55 | 15-15 | 46-64 | EXPANDED | 17 | 17 | normal | . | . | . | 15 | 0 | normal | 2.0138340846723294e-08 | 34.64936393774704 | EXPANDED | 9.443316e-06 | EXPANDED | 1.130153e-04 | EXPANDED |
| NA13503:HTT | 17 | 43 | 17-17 | 43-48 | EXPANDED | 17 | 43 | 17-17 | 17-50 | EXPANDED | 17 | 76 | 17-20 | 61-92 | EXPANDED | 17 | 43 | 17-17 | 43-52 | EXPANDED | 19 | 22 | normal | . | . | . | 17 | 0 | normal | 4.654634605179165e-15 | 43.461609572066756 | EXPANDED | 9.443316e-06 | EXPANDED | 8.072524e-06 | EXPANDED |
| NA13504:HTT | 16 | 52 | 16-16 | 50-54 | EXPANDED | 16 | 52 | 16-16 | 50-55 | EXPANDED | 16 | 39 | 16-16 | 38-39 | normal | 16 | 40 | 16-16 | 40-63 | EXPANDED | 18 | 18 | normal | 30 | 27 | normal | 16 | 14 | normal | 0.00034028101454248994 | 29.83741359552382 | EXPANDED | 9.443316e-06 | EXPANDED | 1.291604e-04 | EXPANDED |
| NA13505:HTT | 22 | 59 | 22-22 | 52-67 | EXPANDED | 22 | 52 | 22-22 | 50-54 | EXPANDED | 22 | 56 | 22-22 | 30-62 | EXPANDED | 22 | 44 | 22-22 | 44-57 | EXPANDED | 24 | 24 | normal | 33 | 33 | normal | . | . | . | 3.2194201976482876e-10 | 36.957130619087295 | EXPANDED | 9.443316e-06 | EXPANDED | 8.072524e-06 | EXPANDED |
| NA13506:HTT | 17 | 59 | 17-17 | 50-65 | EXPANDED | 17 | 46 | 17-17 | 17-50 | EXPANDED | 17 | 52 | 17-17 | 41-57 | EXPANDED | 17 | 46 | 17-17 | 46-54 | EXPANDED | 19 | 19 | normal | 30 | 29 | normal | 17 | 0 | normal | 1.8611334182995864e-13 | 41.53480631786409 | EXPANDED | 9.443316e-06 | EXPANDED | 8.072524e-06 | EXPANDED |
| NA06926:ATXN1 | 30 | 60 | 30-30 | 52-69 | EXPANDED | 30 | 53 | 30-30 | 50-56 | EXPANDED | 33 | 62 | 30-58 | 52-71 | EXPANDED | 29 | 36 | 29-29 | 36-65 | normal | 30 | 30 | normal | . | . | . | . | . | . | 2.6723432691580394e-16 | 55.28991375976906 | EXPANDED | 6.610322e-05 | EXPANDED | 8.072524e-06 | EXPANDED |
| NA13536:ATXN1 | 32 | 38 | 32-32 | 35-41 | normal | 32 | 43 | 32-32 | 32-50 | EXPANDED | 31 | 36 | 31-31 | 33-38 | normal | 31 | 37 | 31-31 | 37-37 | normal | 32 | 32 | normal | 32 | 32 | normal | 31 | 0 | normal | 6.65557331165976e-11 | 46.883519932305724 | EXPANDED | 2.020870e-03 | EXPANDED | 2.906108e-04 | EXPANDED |
| NA13537:ATXN1 | 33 | 68 | 33-33 | 56-79 | EXPANDED | 54 | 57 | 50-58 | 50-65 | EXPANDED | 32 | 68 | 24-62 | 59-81 | EXPANDED | 43 | 55 | 43-53 | 52-63 | EXPANDED | . | . | . | . | . | . | . | . | . | 1.8805157226315128e-19 | 60.6707761522461 | EXPANDED | 9.443316e-06 | EXPANDED | 8.072524e-06 | EXPANDED |
| NA06151:ATXN3 | 56 | 63 | 50-63 | 50-72 | EXPANDED | 21 | 57 | 21-21 | 50-64 | EXPANDED | 18 | 63 | 18-18 | 55-71 | EXPANDED | 18 | 55 | 18-18 | 51-64 | normal | 24 | 24 | normal | . | . | . | . | . | . | 2.268972274384911e-11 | 49.00200868905826 | EXPANDED | 3.777327e-05 | EXPANDED | 8.072524e-06 | EXPANDED |
| NA23709:AR | 52 | 0 | 50-54 | 0-0 | EXPANDED | 46 | 0 | 46-46 | 0-0 | EXPANDED | 52 | 0 | 28-62 | 0-0 | EXPANDED | 46 | 46 | 46-53 | 46-62 | EXPANDED | . | . | . | . | . | . | . | . | . | 0.0249664522983929 | 40.00464435411614 | EXPANDED | 9.443316e-06 | EXPANDED | 8.072524e-06 | EXPANDED |
| hg19C9ORF721000:C9ORF72 | 3 | 1058 | 3-3 | 981-1137 | EXPANDED | 3 | 82 | 3-3 | 62-100 | EXPANDED | 3 | 856 | 3-3 | 834-884 | EXPANDED | 3 | 87 | 3-3 | 70-111 | EXPANDED | 10 | 10 | normal | . | . | . | 3 | 0 | normal | 0.00072 | 32.2 | EXPANDED | . | . | 5.817994e-04 | EXPANDED |
| hg19C9ORF72500:C9ORF72 | 3 | 486 | 3-3 | 435-538 | EXPANDED | 3 | 94 | 3-3 | 72-115 | EXPANDED | 3 | 3 | 3-3 | 3-3 | normal | 3 | 96 | 3-3 | 78-122 | EXPANDED | 10 | 10 | normal | . | . | . | 3 | 0 | normal | 4.2e-05 | 39.6 | EXPANDED | . | . | 5.817994e-04 | EXPANDED |
| hg19C9ORF7260:C9ORF72 | 3 | 70 | 3-3 | 53-85 | EXPANDED | 3 | 73 | 3-3 | 52-85 | EXPANDED | 3 | 3 | 3-3 | 3-3 | normal | 3 | 63 | 3-3 | 55-74 | EXPANDED | 10 | 10 | normal | . | . | . | 3 | 0 | normal | 6.7e-09 | 65.2 | EXPANDED | . | . | 6.464438e-04 | EXPANDED |
| hg19FMR21000:FMR2 | 15 | 156 | 15-15 | 120-190 | normal | 20 | 164 | 20-20 | 125-198 | normal | 15 | 15 | 7-15 | 7-15 | normal | 15 | 15 | 15-15 | 15-15 | normal | . | . | . | . | . | . | 15 | 0 | normal | 3e-32 | 153.1 | EXPANDED | 9.443316e-06 | EXPANDED | 8.138153e-06 | EXPANDED |
| hg19FMR2200:FMR2 | 15 | 160 | 15-15 | 123-193 | normal | 20 | 161 | 20-20 | 124-196 | normal | 15 | 15 | 7-15 | 7-15 | normal | 15 | 15 | 15-15 | 15-15 | normal | . | . | . | . | . | . | 15 | 0 | normal | 8.6e-33 | 156.1 | EXPANDED | 9.443316e-06 | EXPANDED | 8.138153e-06 | EXPANDED |
| hg19FMR2500:FMR2 | 15 | 150 | 15-15 | 113-183 | normal | 20 | 156 | 20-20 | 119-192 | normal | 15 | 15 | 7-15 | 7-15 | normal | 15 | 15 | 15-15 | 15-15 | normal | . | . | . | . | . | . | 15 | 0 | normal | 3e-32 | 153.1 | EXPANDED | 9.443316e-06 | EXPANDED | 8.138153e-06 | EXPANDED |

**SUPPLEMENTARY TABLE 7.** Summary of full-mutation (FM) alleles of the EGA and simulated genomes (with Isaac, a, and BWA-MEM, b, alignment) detected by ExpansionHunter versions 2 and 3 (EH_v2 and EH_v3), GangSTR, and TREDPARSE for *AR*, *ATN1*, *ATXN1*, *ATXN3*, *C9ORF72*, *DMPK*, *FMR1*, *FMR2*, *FXN*, and *HTT* genes based on genotyped repeat size. The allelic classification determined by each tool is sorted into normal (NL), intermediate (IM), premutation (PM), reduced penetrance (RP), or FM bins. The FM threshold is the non-inclusive pathogenic lower bound for each STR locus. The true number (n) of known FM alleles is indicated in parenthesis along with the gene. False positives are also indicated for each STR detection tool.

a.

| | *AR* (n=1) | | *ATN1* (n=2) | *ATXN1* (n=3) | | *ATXN3* (n=1) | | *C9orf72* (n=3) | | *DMPK* (n=17) | | *FMR1* (n=18) | | | | *FMR2* (n=3) | | | *FXN* NL/FM (n=11) | | | *FXN* FM/FM (n=14) | | | | | *HTT* NL/FM (n=12) | | | *HTT* FM/FM (n=1) | False-positives | Total FM detected | Total FM known | Sensitivity |
|---|---|---|---|---|---|---|---|---|---|---|---|---|---|---|---|---|---|---|---|---|---|---|---|---|---|---|---|---|---|---|---|---|---|---|
| FM Threshold | 37 repeats | | 47 repeats | 38 repeats | | 59 repeats | | 60 repeats | | 50 repeats | | 200 repeats | | | | 200 repeats | | | 65 repeats | | | | | | | | 39 repeats | | | | | | | |
| Allelic classification | FM | NL | FM | FM | IM | FM | IM | FM | NL | FM | NL | FM | PM | IM | NL | FM | PM | NL | FM/FM | NL/FM | NL/NL | FM/FM | PM/FM | NL/FM | NL/PM | NL/NL | NL/FM | NL/RP | NL/IM | FM/FM | | | | |
| EH_v2 | 1 | . | 2 | 2 | 1 | 1 | . | 3 | . | 17 | . | 1 | 17 | . | . | 0 | 3 | . | 1 | 10 | . | 8 | 3 | 3 | . | . | 12 | . | . | 1 | 7 | 74 | 101 | 0.7326733 |
| EH_v3 | 1 | . | 2 | 3 | . | 0 | 1 | 3 | . | 17 | . | 0 | 18 | . | . | 0 | 3 | . | . | 11 | . | 11 | . | 3 | . | . | 12 | . | . | 1 | 5 | 76 | 101 | 0.7524752 |
| GangSTR | 0 | 1 | 2 | 2 | 1 | 0 | 1 | 0 | 3 | 16 | 1 | 0 | 4 | . | 14 | 0 | . | 3 | . | 3 | 8 | 0 | . | 13 | 1 | . | 10 | 1 | 1 | 1 | 8 | 48 | 101 | 0.4752475 |
| TREDPARSE | 1 | . | 2 | 1 | 2 | 0 | 1 | 3 | . | 17 | . | 0 | 15 | 1 | 2 | 0 | . | 3 | . | 11 | . | 0 | 11 | 3 | . | . | 12 | . | . | 1 | 3 | 63 | 101 | 0.6237624 |

b.

| | *AR* (n=1) | *ATN1* (n=2) | *ATXN1* (n=3) | | *ATXN3* (n=1) | | *C9orf72* (n=3) | | *DMPK* (n=17) | | *FMR1* (n=18) | | | | *FMR2* (n=3) | | | *FXN* NL/FM (n=11) | | | *FXN* FM/FM (n=14) | | | | *HTT* NL/FM (n=12) | | | *HTT* FM/FM (n=1) | | | | |
|---|---|---|---|---|---|---|---|---|---|---|---|---|---|---|---|---|---|---|---|---|---|---|---|---|---|---|---|---|---|---|---|---|
| FM Threshold | 37 repeats | 47 repeats | 38 repeats | | 59 repeats | | 60 repeats | | 50 repeats | | 200 repeats | | | | 200 repeats | | | 65 repeats | | | | | | | 39 repeats | | | | | Total FM | Total FM | |
| Allelic classification | FM | FM | FM | IM | FM | IM | FM | NL | FM | NL | FM | PM | IM | NL | FM | PM | NL | FM/FM | NL/FM | NL/NL | FM/FM | PM/FM | NL/FM | NL/NL | NL/FM | NL/RP | NL/IM | FM/FM | False-positives | detected | known | Sensitivity |
| EH_v2 | 1 | 2 | 2 | 1 | 1 | . | 3 | . | 17 | . | 0 | 18 | . | . | 0 | 3 | . | 1 | 10 | . | 11 | . | 3 | . | 12 | . | . | 1 | 7 | 76 | 101 | 0.752475248 |
| EH_v3 | 1 | 2 | 3 | . | 0 | 1 | 3 | . | 17 | . | 0 | 18 | . | . | 0 | 3 | . | . | 11 | . | 13 | . | 1 | . | 12 | . | . | 1 | 5 | 78 | 101 | 0.772277228 |
| GangSTR | 1 | 2 | 2 | 1 | 1 | . | 1 | 2 | 16 | 1 | 0 | 3 | . | 15 | 0 | . | 3 | . | 0 | 11 | 0 | . | . | 14 | 9 | 1 | 2 | 1 | 8 | 34 | 101 | 0.336633663 |
| TREDPARSE | 1 | 2 | 1 | 2 | 0 | 1 | 3 | . | 17 | . | 0 | 16 | . | 2 | 0 | . | 3 | . | 11 | . | 0 | 13 | 1 | . | 12 | . | . | 1 | 10 | 63 | 101 | 0.623762376 |

**SUPPLEMENTARY TABLE 8.** Genotyping of known normal (NL) (a) and intermediate/premutation (IM/PM) (b) alleles in the Isaac-aligned EGA and simulated genomes using ExpansionHunter versions 2 and 3 (EH_v2 and EH_v3), GangSTR, TREDPARSE, lobSTR, HipSTR, and RepeatSeq. Based on the genotyped calls, the alleles were sorted into NL, IM, PM, or FM bins. The true number (n) of known NL, IM, or PM alleles in each of the analysed disease STR locus is indicated in parenthesis.

a.

| | *ATN1* (n=2) | *ATXN1* (n=3) | | *ATXN3* (n=1) | | *C9ORF72* (n=3) | *DMPK* (n=17) | *FMR1* (n=42) | | | *FMR2* (n=3) | *FXN* (n=11) | | *HTT* (n=12) | | Total NL detected | Total NL present |
|---|---|---|---|---|---|---|---|---|---|---|---|---|---|---|---|---|---|
| | NL | NL | FM | NL | IM | NL | NL | NL | IM | PM | NL | NL | FM | NL | IM | | |
| EH_v2 | 2 | 3 | . | 0 | 1 | 3 | 17 | 41 | . | 1 | 3 | 10 | 1 | 12 | . | 91 | 94 |
| EH_v3 | 2 | 2 | 1 | 1 | . | 3 | 17 | 33 | 8 | 1 | 3 | 11 | . | 12 | . | 84 | 94 |
| GangSTR | 2 | 3 | . | 1 | . | 3 | 17 | 42 | . | . | 3 | 11 | . | 12 | . | 94 | 94 |
| TREDPARSE | 2 | 2 | 1 | 1 | . | 3 | 17 | 42 | . | . | 3 | 11 | . | 12 | . | 93 | 94 |
| lobSTR | 2 | 2 | . | 1 | . | 3 | 17 | 29 | . | . | 0 | 7 | . | 10 | . | 71 | 94 |
| HipSTR | 0 | 2 | . | 1 | . | 1 | 3 | 21 | . | . | 0 | 2 | . | 0 | 4 | 30 | 94 |
| RepeatSeq | 2 | 1 | . | 1 | . | 0 | 16 | 26 | . | . | 0 | 1 | . | 7 | . | 54 | 94 |

b.

| | TRUE IMs | | | TRUE PMs | | | Total detected IM/PM | Total known IM/PM |
|---|---|---|---|---|---|---|---|---|
| | *FMR1* (n=5) | | | *FMR1* (n=16) | | | | |
| | NL | IM | PM | NL | IM | PM | | |
| EH_v2 | 1 | 2 | 2 | . | . | 16 | 18 | 21 |
| EH_v3 | 2 | 0 | 3 | . | . | 16 | 16 | 21 |
| GangSTR | 4 | 0 | 1 | 9 | 1 | 6 | 6 | 21 |
| TREDPARSE | 3 | 1 | 1 | 6 | 3 | 7 | 8 | 21 |
| lobSTR | 2 | 0 | . | 6 | . | 0 | 0 | 21 |
| HipSTR | 2 | 0 | . | 5 | . | 0 | 0 | 21 |
| RepeatSeq | 3 | 0 | . | 3 | . | 0 | 0 | 21 |

**SUPPLEMENTARY TABLE 9.** Genotyping of known normal (NL) (a) and intermediate/premutation (IM/PM) (b) alleles in the BWA-aligned EGA and simulated genomes using ExpansionHunter versions 2 and 3 (EH_v2 and EH_v3), GangSTR, TREDPARSE, lobSTR, HipSTR, and RepeatSeq. Based on the genotyped calls, the alleles were sorted into NL, IM, PM, or FM bins. The true number (n) of known NL, IM, or PM alleles in each of the analysed disease STR locus is indicated in parenthesis.

a.

| | *ATN1* (n=2) | *ATXN1* (n=3) | | *ATXN3* (n=1) | | *C9ORF72* (n=3) | *DMPK* (n=17) | *FMR1* (n=42) | | | *FMR2* (n=3) | *FXN* (n=11) | | *HTT* (n=12) | | Total NL detected | Total NL present |
|---|---|---|---|---|---|---|---|---|---|---|---|---|---|---|---|---|---|
| | NL | NL | FM | NL | IM | NL | NL | NL | IM | PM | NL | NL | FM | NL | IM | | |
| EH_v2 | 2 | 3 | | 0 | 1 | 3 | 17 | 41 | . | 1 | 3 | 10 | 1 | 12 | . | 91 | 94 |
| EH_v3 | 2 | 2 | 1 | 1 | . | 3 | 17 | 33 | 8 | 1 | 3 | 11 | . | 12 | . | 84 | 94 |
| GangSTR | 2 | 3 | . | 1 | . | 3 | 16 | 42 | . | . | 3 | 11 | . | 12 | . | 93 | 94 |
| TREDPARSE | 2 | 2 | 1 | 1 | . | 3 | 17 | 42 | . | . | 3 | 11 | . | 12 | . | 93 | 94 |
| lobSTR | 2 | 2 | . | 1 | . | 3 | 17 | 22 | . | . | 0 | 9 | . | 11 | . | 67 | 94 |
| HipSTR | 0 | 1 | . | 0 | . | 0 | 6 | 19 | . | . | 0 | 4 | . | 0 | 4 | 30 | 94 |
| RepeatSeq | 2 | 1 | . | 0 | . | 3 | 15 | 6 | . | . | 3 | 4 | . | 10 | . | 44 | 94 |

b.

| | TRUE IMs | | | TRUE PMs | | | Total detected IM/PM | Total known IM/PM |
|---|---|---|---|---|---|---|---|---|
| | *FMR1* (n=5) | | | *FMR1* (n=16) | | | | |
| | NL | IM | PM | NL | IM | PM | | |
| EH_v2 | 1 | 3 | 1 | . | 1 | 15 | 18 | 21 |
| EH_v3 | 2 | 0 | 3 | . | . | 16 | 16 | 21 |
| GangSTR | 5 | 0 | . | 15 | 1 | 0 | 0 | 21 |
| TREDPARSE | 2 | 3 | . | 3 | 6 | 7 | 10 | 21 |
| lobSTR | 1 | 0 | . | 4 | . | 0 | 0 | 21 |
| HipSTR | 1 | 0 | . | 2 | . | 0 | 0 | 21 |
| RepeatSeq | . | 0 | . | 1 | . | 0 | 0 | 21 |

**SUPPLEMENTARY TABLE 10.** Summary of full-mutation (FM) alleles of the EGA and simulated genomes (with Isaac, a, and BWA-MEM, b, alignment) detected by ExpansionHunter version 2 (EH_v2) (default; without off-target sites, no OT; and with off-target sites, w OT) and GangSTR (no OT and w OT) for *C9ORF72*, *DMPK*, *FMR1*, *FMR2*, and *FXN* genes. The allelic classification determined by each tool is sorted into normal (NL), intermediate (IM), premutation (PM), or FM bins. The true number (n) of known FM alleles is indicated in parenthesis along with the gene. The FM threshold is the non-inclusive pathogenic lower bound for each STR locus. False positives are totalled for each STR detection tool in the second last column of the table.

a.

| | *C9orf72* (n=3) | | *DMPK* (n=17) | | *FMR1* (n=18) | | | *FMR2* (n=3) | | | *FXN* NL/FM (n=11) | | | *FXN* FM/FM (n=14) | | | | False-positives | Total FM detected | Total FM known | Sensitivity |
|---|---|---|---|---|---|---|---|---|---|---|---|---|---|---|---|---|---|---|---|---|---|
| FM Threshold | 60 repeats | | 50 repeats | | 200 repeats | | | 200 repeats | | | 65 repeats | | | | | | | | | | |
| Allelic classification | FM | NL | FM | NL | FM | PM | NL | FM | PM | NL | FM/FM | NL/FM | NL/NL | FM/FM | PM/FM | NL/FM | NL/PM | | | | |
| EH_v2 default | 3 | . | 17 | . | 1 | 17 | . | 0 | 3 | . | 1 | 10 | . | 8 | 3 | 3 | . | 4 | 54 | 80 | 0.675 |
| EH_v2 no OT | 3 | . | 17 | . | 0 | 18 | . | 0 | 3 | . | 1 | 10 | . | 8 | 3 | 3 | . | 2 | 53 | 80 | 0.6625 |
| EH_v2 w OT | 3 | . | 17 | . | 1 | 17 | . | 2 | 1 | . | 1 | 10 | . | 7 | 4 | 3 | . | 4 | 55 | 80 | 0.6875 |
| GangSTR no OT | 0 | 3 | 16 | 1 | 0 | 4 | 14 | 0 | . | 3 | . | 3 | 8 | 0 | . | 13 | 1 | 0 | 32 | 80 | 0.4 |
| GangSTR w OT | 0 | 3 | 16 | 1 | 0 | 4 | 14 | 0 | . | 3 | . | 3 | 8 | 0 | . | 13 | 1 | 0 | 32 | 80 | 0.4 |

b.

| | *C9orf72* (n=3) | | *DMPK* (n=17) | | | *FMR1* (n=18) | | | *FMR2* (n=3) | | | *FXN* NL/FM (n=11) | | | *FXN* FM/FM (n=14) | | | | | | | |
|---|---|---|---|---|---|---|---|---|---|---|---|---|---|---|---|---|---|---|---|---|---|---|
| FM Threshold | 60 repeats | | 50 repeats | | | 200 repeats | | | 200 repeats | | | 65 repeats | | | | | | | | Total FM | Total FM | |
| Allelic classification | FM | NL | FM | PM | NL | FM | PM | NL | FM | PM | NL | FM/FM | NL/FM | NL/NL | FM/FM | PM/FM | NL/FM | NL/NL | False-positives | detected | known | Sensitivity |
| EH_v2 default | 3 | . | 17 | . | . | 0 | 18 | . | 0 | 3 | . | 1 | 10 | . | 11 | . | 3 | . | 3 | 56 | 80 | 0.7 |
| EH_v2 no OT | 3 | . | 17 | . | . | 0 | 18 | . | 0 | 3 | . | 1 | 10 | . | 11 | . | 3 | . | 2 | 56 | 80 | 0.7 |
| EH_v2 w OT | 3 | . | 17 | . | . | 0 | 18 | . | 2 | 1 | . | 1 | 10 | . | 5 | 7 | 2 | . | 3 | 52 | 80 | 0.65 |
| GangSTR no OT | 0 | 3 | 8 | 5 | 4 | 0 | . | 18 | 0 | . | 3 | . | 0 | 11 | 0 | . | . | 14 | 0 | 8 | 80 | 0.1 |
| GangSTR w OT | 1 | 2 | 16 | . | 1 | 0 | 3 | 15 | 0 | . | 3 | . | 0 | 11 | 0 | . | . | 14 | 0 | 17 | 80 | 0.2125 |

**SUPPLEMENTARY TABLE 11**. Allelic calls from ExpansionHunter versions 2 and 3 (EH_v2 and EH_v3), STRetch, and exSTRa STR detection tools for the CAUSES and IMAGINE genomes and exomes. For the Sample ID, "-P" indicates the proband, "-S" indicates the sibling, "-M" indicates the mother, and "-F" indicates the father. For EH_v2 and EH_v3, a1 and a2 indicate the calls for allele 1 and allele 2, respectively, while CI indicates the confidence interval. In certain cases, "borderline" is used to indicate an allele between a reduced penetrance full-mutation (FM) and a full-penetrance FM. For STRetch, the detected number of base pairs inserted (bpInsertion) and genotyped repeat counts (repeatUnits) are both reported in addition to the p-value (pval). The p-status for STRetch and exSTRa are marked as "EXPANDED" for those samples with p-values <0.05, and "normal" for those samples with p-values >0.05. For all tools except STRetch and exSTRa, empty cells indicate a call was not made for this sample in the corresponding STR locus. The column "No. of support" reports the number of tools that support a positive call for the corresponding patient or parent, while the tools themselves are listed in the next column. Finally, the molecular findings for each sample is either reported or marked as "pending" for those currently being validated.

| Sample ID | Gene | FM_threshold | EH_v2:a1 | EH_v2:a2 | EH_v2:CI-a1 | EH_v2:CI-a2 | EH_v2:status | EH_v3:a1 | EH_v3:a2 | EH_v3:CI-a1 | EH_v3:CI-a2 | EH_v3:status | STRetch:pval | STRetch:bpInsertion | STRetch:repeatUnits | STRetch:status | exSTRa:pval | exSTRa:status | No. of support | Tools supporting the STR Finding | Molecular Findings |
|---|---|---|---|---|---|---|---|---|---|---|---|---|---|---|---|---|---|---|---|---|---|
| 1901-P | *AR* | 37 | 22 | 37 | 22-22 | 37-37 | Full-mutation | 23 | 38 | 23-23 | 38-38 | Full-mutation | 0.051 | 20.7 | 40.2 | normal | 9.605975e-04 | EXPANDED | 2 | EH_v2, EH_v3 & exSTRa | Reduced Penetrance Full-mutation: Allele 1: 22 ± 1 repeats; Allele 2: 37±1 repeats |
| 1901-F | *AR* | 37 | 37 | 0 | 37-37 | 0-0 | Full-mutation | 38 | 0 | 38-38 | 0-0 | Full-mutation | 1.3e-08 | 42.4 | 47.4 | EXPANDED | . | . | 2 | EH_v2, EH_v3 & STRetch | Reduced Penetrance Full-mutation: 37±1 repeats |
| 890-P | *ATXN1* | 38 | 30 | 39 | 30-30 | 39-39 | Full-mutation | 31 | 39 | 31-31 | 31-39 | Full-mutation | . | . | . | . | . | . | 2 | EH_v2 & EH_v3 | Pending |
| 532-M | *ATXN1* | 38 | 38 | 52 | 38-38 | 50-55 | Full-mutation | 30 | 39 | 30-30 | 39-39 | Full-mutation | 0.027 | 17.8 | 36.2 | EXPANDED | 4.170820e-06 | EXPANDED | 4 | EH_v2, EH_v3, STRetch, & exSTRa | Pending |
| 2560-M | *ATXN1* | 38 | 28 | 41 | 28-28 | 41-41 | Full-mutation | 29 | 42 | 29-29 | 42-42 | Full-mutation | 0.45 | 8.8 | 33.2 | normal | . | . | 2 | EH_v2 & EH_v3 | Pending |
| 1411-F | *ATXN1* | 38 | 30 | 52 | 30-30 | 50-54 | Full-mutation | 31 | 41 | 31-31 | 31-41 | Full-mutation | 0.31 | 10.3 | 33.7 | normal | . | . | 2 | EH_v2 & EH_v3 | Pending |
| 821-P | *ATXN2* | 36 | 22 | 35 | 22-22 | 35-35 | borderline | 22 | 35 | 22-22 | 35-35 | borderline | 1.2e-07 | 74.0 | 48.0 | EXPANDED | 4.205869e-06 | EXPANDED | 2 | STRetch & exSTRa | Pending |
| 821-M | *ATXN2* | 36 | 37 | 37 | 37-37 | 37-37 | Full-mutation | 22 | 35 | 22-22 | 35-35 | borderline | 4.8e-10 | 106.0 | 58.6 | EXPANDED | . | . | 3 | EH_v2 & STRetch | Pending |
| 1099-P | *ATXN8* | 75 | 20 | 79 | 20-20 | 67-91 | Full-mutation | . | . | . | . | . | 4.8e-21 | 471.5 | 172.5 | EXPANDED | 8.704281e-06 | EXPANDED | 4 | EH_v2, STRetch, & exSTRa | Pending |
| 235-P | *ATXN8* | 75 | 8 | 111 | 8-8 | 87-132 | Full-mutation | 9 | 87 | 9-9 | 68-104 | Full-mutation | 3.9e-42 | 283.9 | 109.9 | EXPANDED | . | . | 3 | EH_v2, EH_v3, & STRetch | Pending |
| 235-M | *ATXN8* | 75 | 17 | 95 | 17-17 | 73-113 | Full-mutation | 18 | 74 | 18-18 | 57-88 | normal | 5.8e-33 | 190.1 | 78.7 | EXPANDED | . | . | 2 | EH_v2 & STRetch | Pending |
| 2010-P | *DMPK* | 49 | 11 | 52 | 11-11 | 50-54 | Full-mutation | 11 | 56 | 11-11 | 51-62 | Full-mutation | 0.3 | 10.9 | 24.3 | normal | 1.334653e-03 | EXPANDED | 3 | EH_v2, EH_v3, & exSTRa | Positive: 150 repeats |
| 2010-M | *DMPK* | 49 | 52 | 56 | 51-54 | 50-61 | Full-mutation | 51 | 57 | 51-51 | 51-63 | Full-mutation | . | . | . | . | 6.903376e-04 | EXPANDED | 3 | EH_v2, EH_v3, & exSTRa | Positive: 430 repeats |
| 699-M | *FMR1* | 54 | 29 | 55 | 29-29 | 50-60 | Premutation | 29 | 55 | 29-29 | 50-60 | Premutation | 0.4 | 27.1 | 34.0 | normal | . | . | 2 | EH_v2 & EH_v3 | Pending |
| 148-M | *FMR1* | 54 | 20 | 63 | 20-20 | 52-73 | Premutation | 20 | 57 | 20-20 | 50-64 | Premutation | 0.0012 | 100.0 | 58.3 | EXPANDED | . | . | 3 | EH_v2, EH_v3, & STRetch | Pending |
| 800-F | *FMR1* | 54 | 56 | 0 | 50-62 | 0-0 | Premutation | 53 | 0 | 51-55 | 0-0 | Intermediate | 2.6e-06 | 36.0 | 37.0 | EXPANDED | . | . | 2 | EH_v2 & STRetch | Pending |
| 800-P | *FMR1* | 54 | 31 | 55 | 31-31 | 50-60 | Premutation | 31 | 54 | 31-31 | 51-58 | Intermediate | 0.97 | 29.9 | 35.0 | normal | . | . | 1 | EH_v2 | Pending |
| 480-P | *FMR1* | 54 | 52 | 0 | 50-55 | 0-0 | Intermediate | 60 | 0 | 50-68 | 0-0 | Premutation | . | . | . | . | 1.805216e-03 | EXPANDED | 2 | EH_v3 & exSTRa | Pending |
| 712-M | *FMR1* | 54 | 31 | 52 | 31-31 | 50-54 | Intermediate | 31 | 57 | 31-31 | 50-64 | Premutation | 0.072 | 53.4 | 42.8 | normal | 2.566658e-05 | EXPANDED | 2 | EH_v3 & exSTRa | Pending |
| 925-P | *FMR1* | 54 | 38 | 0 | 38-38 | 0-0 | normal | 56 | 0 | 50-62 | 0-0 | Premutation | 0.4 | 36.5 | 37.2 | normal | 2.177382e-03 | EXPANDED | 2 | EH_v3 & exSTRa | Negative |
| 925-S | *FMR1* | 54 | 50 | 0 | 50-55 | 0-0 | Intermediate | 53 | 0 | 50-56 | 0-0 | Intermediate | 0.13 | 45.9 | 40.3 | normal | 4.277764e-06 | EXPANDED | 2 | EH_v3 & exSTRa | Pending |
| 925-M | *FMR1* | 54 | 38 | 62 | 38-38 | 52-72 | Premutation | 38 | 57 | 38-38 | 50-65 | Premutation | 0.00064 | 109.4 | 61.5 | EXPANDED | . | . | 3 | EH_v2, EH_v3, & STRetch | Pending |
| 1987-F | *FXN* | 65 | 9 | 52 | 9-9 | 50-54 | Intermediate/ Premutation | 9 | 69 | 9-9 | 57-79 | Full-mutation | . | . | . | . | 4.488774e-06 | EXPANDED | 2 | EH_v3 & exSTRa | Pending |
| 1530-P | *HTT* | 39 | 23 | 37 | 23-23 | 37-37 | Full-mutation | 23 | 37 | 23-23 | 37-37 | Full-mutation | 9.4e-07 | 35.1 | 33.0 | EXPANDED | 4.488774e-06 | EXPANDED | 2 | EH_v2, EH_v3, STRetch, & exSTRa | Reduced Penetrance Full-mutation: Allele 1: 23 ± 1 repeats; Allele 2: 37 ± 1 repeats |
| 1530-F | *HTT* | 39 | 17 | 37 | 17-17 | 37-37 | Full-mutation | 17 | 37 | 17-17 | 37-37 | Full-mutation | 1e-14 | 68.6 | 44.2 | EXPANDED | 4.488774e-06 | EXPANDED | 2 | EH_v2, EH_v3, STRetch, & exSTRa | Reduced Penetrance Full-mutation: Allele 1: 17 ± 1 repeats; Allele 2: 37 ± 1 repeats |
| 1992-M | *TBP* | 48 | 36 | 52 | 36-36 | 50-55 | Full-mutation | 35 | 53 | 35-35 | 51-56 | Full-mutation | 0.67 | 14.5 | 41.8 | normal | . | . | 2 | EH_v2 & EH_v3 | Negative; Allele 1: 36 ± 1 repeats; Allele 2: 37±1 repeats |
| 2990-M | *TBP* | 48 | 37 | 52 | 37-37 | 50-53 | Full-mutation | 36 | 54 | 36-36 | 51-57 | Full-mutation | 0.38 | 9.5 | 40.2 | normal | . | . | 2 | EH_v2 & EH_v3 | Pending |

**SUPPLEMENTARY TABLE 12.** Summary of the allelic distributions of repeat lengths for STR loci in *AR*, *ATN1*, *ATXN1*, *ATXN2*, *ATXN3*, *ATXN7*, *ATXN8/ATXN8OS*, *ATXN10*, *C9ORF72*, *CACNA1A*, *CBL*, *CNBP*, *CSTB*, *DMPK*, *FMR1*, *FMR2*, *FXN*, *HTT*, *JPH3*, *NOP56*, *PHOX2B*, *PPP2R2B*, and *TBP* genes for CAUSES and IMAGINE genomes as genotyped by ExpansionHunter versions 2 and 3 (EH_v2 and EH_v3) and GangSTR. Mode 1, 2 and 3 are the three most common alleles genotyped, while Lowest and Highest indicate the lower and upper bound of the range of alleles detected. The n next to each value indicates the number of alleles at the specified repeat length. The Total n value indicates the total number of alleles genotyped by the tool at the specified STR locus.

| Gene | EH_v2 | | | | | | | | | | | EH_v3 | | | | | | | | | | | GangSTR | | | | | | | | | | | Consensus (Caucasian)[REF] |
|---|---|---|---|---|---|---|---|---|---|---|---|---|---|---|---|---|---|---|---|---|---|---|---|---|---|---|---|---|---|---|---|---|---|---|
| | Mode 1 | n | Mode 2 | n | Mode 3 | n | Lowest | n | Highest | n | Total n | Mode 1 | n | Mode 2 | n | Mode 3 | n | Lowest | n | Highest | n | Total n | Mode 1 | n | Mode 2 | n | Mode 3 | n | Lowest | n | Highest | n | Total n | |
| *AR* | 21 | 115 | 23 | 90 | 20 | 89 | 13 | 1 | 37 | 2 | 689 | 22 | 113 | 24 | 90 | 25 | 89 | 1 | 1 | 38 | 2 | 681 | 21 | 136 | 20 | 109 | 23 | 104 | 11 | 1 | 37 | 2 | 819 | 21[1] |
| *ATN1* | 15 | 292 | 16 | 152 | 10 | 118 | 7 | 9 | 24 | 4 | 934 | 19 | 284 | 20 | 153 | 14 | 118 | 11 | 9 | 28 | 4 | 930 | 15 | 349 | 16 | 181 | 10 | 145 | 7 | 9 | 24 | 6 | 1100 | 15[2] |
| *ATXN1* | 30 | 320 | 29 | 254 | 31 | 95 | 8 | 2 | 52 | 1 | 934 | 31 | 316 | 30 | 250 | 32 | 95 | 8 | 1 | 40 | 1 | 930 | 30 | 380 | 29 | 310 | 31 | 111 | 19 | 1 | 53 | 1 | 1100 | 30[2] |
| *ATXN2* | 22 | 831 | 23 | 55 | 27 | 12 | 15 | 2 | 34 | 1 | 934 | 22 | 813 | 23 | 57 | 27 | 14 | 9 | 3 | 34 | 1 | 930 | 22 | 983 | 23 | 63 | 27 | 13 | 16 | 3 | 34 | 2 | 1100 | 22[2] |
| *ATXN3* | 17 | 249 | 8 | 217 | 21 | 155 | 8 | 217 | 52 | 1 | 934 | 20 | 247 | 11 | 216 | 24 | 152 | 11 | 216 | 44 | 1 | 930 | 17 | 305 | 8 | 266 | 21 | 131 | 1 | 2 | 32 | 2 | 1100 | 23[2] |
| *ATXN7* | 10 | 723 | 12 | 111 | 11 | 39 | 2 | 1 | 23 | 1 | 934 | 10 | 711 | 12 | 112 | 11 | 40 | 2 | 1 | 23 | 1 | 930 | 10 | 843 | 12 | 133 | 11 | 52 | 7 | 8 | 14 | 14 | 1100 | 10[3] |
| *ATXN8* | 12 | 178 | 8 | 174 | 15 | 170 | 5 | 3 | 111 | 1 | 934 | 12 | 204 | 15 | 194 | 9 | 186 | 6 | 6 | 87 | 1 | 930 | 15 | 311 | 13 | 218 | 8 | 206 | 5 | 3 | 41 | 1 | 1100 | 23[4] |
| *ATXN10* | 14 | 276 | 13 | 218 | 15 | 139 | 10 | 3 | 21 | 2 | 934 | 14 | 278 | 13 | 213 | 15 | 138 | 10 | 4 | 31 | 1 | 930 | 14 | 320 | 13 | 261 | 15 | 160 | 10 | 6 | 21 | 1 | 1100 | 13[4] |
| *C9ORF72* | 2 | 481 | 8 | 124 | 5 | 99 | 2 | 481 | 30 | 1 | 934 | 2 | 479 | 8 | 123 | 5 | 96 | 2 | 479 | 28 | 1 | 930 | 2 | 570 | 8 | 139 | 5 | 120 | 2 | 570 | 17 | 2 | 1100 | 2[5] |
| *CACNA1A* | 13 | 321 | 11 | 289 | 12 | 145 | 4 | 6 | 18 | 1 | 934 | 13 | 319 | 11 | 287 | 12 | 145 | 4 | 8 | 18 | 1 | 930 | 13 | 372 | 11 | 340 | 12 | 179 | 4 | 6 | 18 | 1 | 1100 | 13[2] |
| *CBL* | 11 | 634 | 12 | 87 | 14 | 73 | 6 | 1 | 52 | 1 | 934 | 11 | 630 | 12 | 87 | 14 | 72 | 5 | 1 | 44 | 1 | 930 | 11 | 739 | 12 | 106 | 14 | 91 | 7 | 7 | 39 | 1 | 1100 | 11[6] |
| *CNBP* | 15 | 381 | 14 | 175 | 7 | 102 | 3 | 1 | 27 | 1 | 934 | 15 | 536 | 16 | 154 | 7 | 66 | 1 | 2 | 41 | 1 | 930 | 7 | 306 | 15 | 261 | 14 | 165 | 4 | 1 | 42 | 1 | 1100 | 14[7] |
| *CSTB* | 3 | 547 | 2 | 383 | 5 | 1 | 2 | 383 | 19 | 1 | 934 | 3 | 530 | 2 | 382 | 8 | 5 | 2 | 382 | 18 | 1 | 930 | 3 | 640 | 2 | 456 | 5 | 1 | 2 | 456 | 15 | 1 | 1100 | 2-3[8] |
| *DMPK* | 5 | 312 | 13 | 161 | 12 | 139 | 4 | 2 | 33 | 2 | 934 | 5 | 312 | 13 | 159 | 12 | 135 | 4 | 2 | 41 | 2 | 930 | 5 | 388 | 13 | 178 | 11 | 153 | 5 | 388 | 33 | 2 | 1100 | 5[9] |
| *FMR1* | 30 | 244 | 29 | 157 | 31 | 45 | 7 | 2 | 63 | 1 | 681 | 30 | 218 | 29 | 135 | 23 | 38 | 7 | 2 | 58 | 1 | 645 | 29 | 214 | 30 | 169 | 31 | 64 | 2 | 1 | 43 | 1 | 807 | 29-31[10] |
| *FMR2* | 15 | 211 | 18 | 143 | 17 | 62 | 4 | 3 | 52 | 2 | 677 | 20 | 204 | 23 | 128 | 22 | 55 | 7 | 1 | 53 | 1 | 625 | 15 | 259 | 18 | 147 | 17 | 76 | 4 | 3 | 43 | 2 | 772 | 16[11] |
| *FXN* | 9 | 473 | 8 | 295 | 17 | 27 | 2 | 1 | 52 | 1 | 934 | 9 | 513 | 8 | 255 | 18 | 27 | 2 | 2 | 69 | 1 | 930 | 8 | 414 | 9 | 379 | 10 | 50 | 2 | 2 | 27 | 2 | 1100 | 9[12] |
| *HTT* | 17 | 387 | 18 | 131 | 19 | 102 | 4 | 1 | 37 | 2 | 934 | 17 | 395 | 18 | 132 | 19 | 93 | 1 | 1 | 37 | 2 | 930 | 17 | 403 | 18 | 144 | 19 | 119 | 4 | 1 | 37 | 2 | 1100 | 17[13] |
| *JPH3* | 14 | 517 | 16 | 255 | 15 | 88 | 7 | 2 | 26 | 1 | 934 | 14 | 508 | 16 | 255 | 15 | 88 | 7 | 3 | 26 | 1 | 930 | 14 | 612 | 16 | 292 | 15 | 109 | 6 | 1 | 24 | 1 | 1100 | 12[14] |
| *NOP56* | 4 | 151 | 5 | 88 | 6 | 38 | 1 | 6 | 13 | 2 | 376 | 4 | 264 | 7 | 245 | 6 | 203 | 3 | 82 | 14 | 1 | 930 | 9 | 529 | 7 | 235 | 6 | 184 | 5 | 8 | 13 | 2 | 1100 | 9[15] |
| *PHOX2B* | 4 | 102 | 3 | 90 | 6 | 68 | 1 | 2 | 18 | 2 | 478 | 20 | 793 | 52 | 25 | 15 | 12 | 13 | 3 | 55 | 1 | 930 | 20 | 1079 | 15 | 14 | 13 | 3 | 13 | 3 | 25 | 2 | 1100 | 20[16] |
| *PPP2R2B* | 10 | 473 | 13 | 162 | 15 | 107 | 6 | 1 | 27 | 1 | 934 | 10 | 472 | 13 | 160 | 15 | 107 | 6 | 1 | 27 | 1 | 930 | 10 | 563 | 13 | 190 | 15 | 122 | 6 | 1 | 26 | 3 | 1100 | 10[17] |
| *TBP* | 38 | 273 | 36 | 229 | 37 | 228 | 18 | 1 | 52 | 2 | 934 | 37 | 239 | 36 | 197 | 35 | 196 | 1 | 1 | 53 | 2 | 930 | 34 | 268 | 35 | 216 | 36 | 199 | 1 | 4 | 57 | 1 | 1100 | 38[18] |

**Supplementary Figure 1.** exSTRa empirical cumulative distribution function (ECDF) plots of Isaac- (A) and BWA-aligned (without controls, B, and with 100 controls, C) EGA and simulated genomes. ECDF plots represent STR repeats found in each read of a sample as a step function from the smallest number of repeats to the largest. Therefore, a right shifted plot would indicate an STR expansion in the sample. In the figure, a pair of plots are shown for each STR locus (*AR*, *ATN1*, *ATXN1*, *ATXN3*, *C9ORF72*, *DMPK*, *FMR1*, *FMR2*, *FXN*, and *HTT*). Of each pair, the plot on the left highlights samples which are known to have STR expansions at the given locus. The plot on the right shows samples which were called by exSTRa as having repeat expansions at the given locus. The title on top of each plot indicates the STR locus, the number of repeats in the hg19 reference genome, and the pathogenic lowerbound number of repeats of the STR.

### *A. exSTRa ECDF and t-sum plots of Isaac-aligned EGA and simulated genomes*

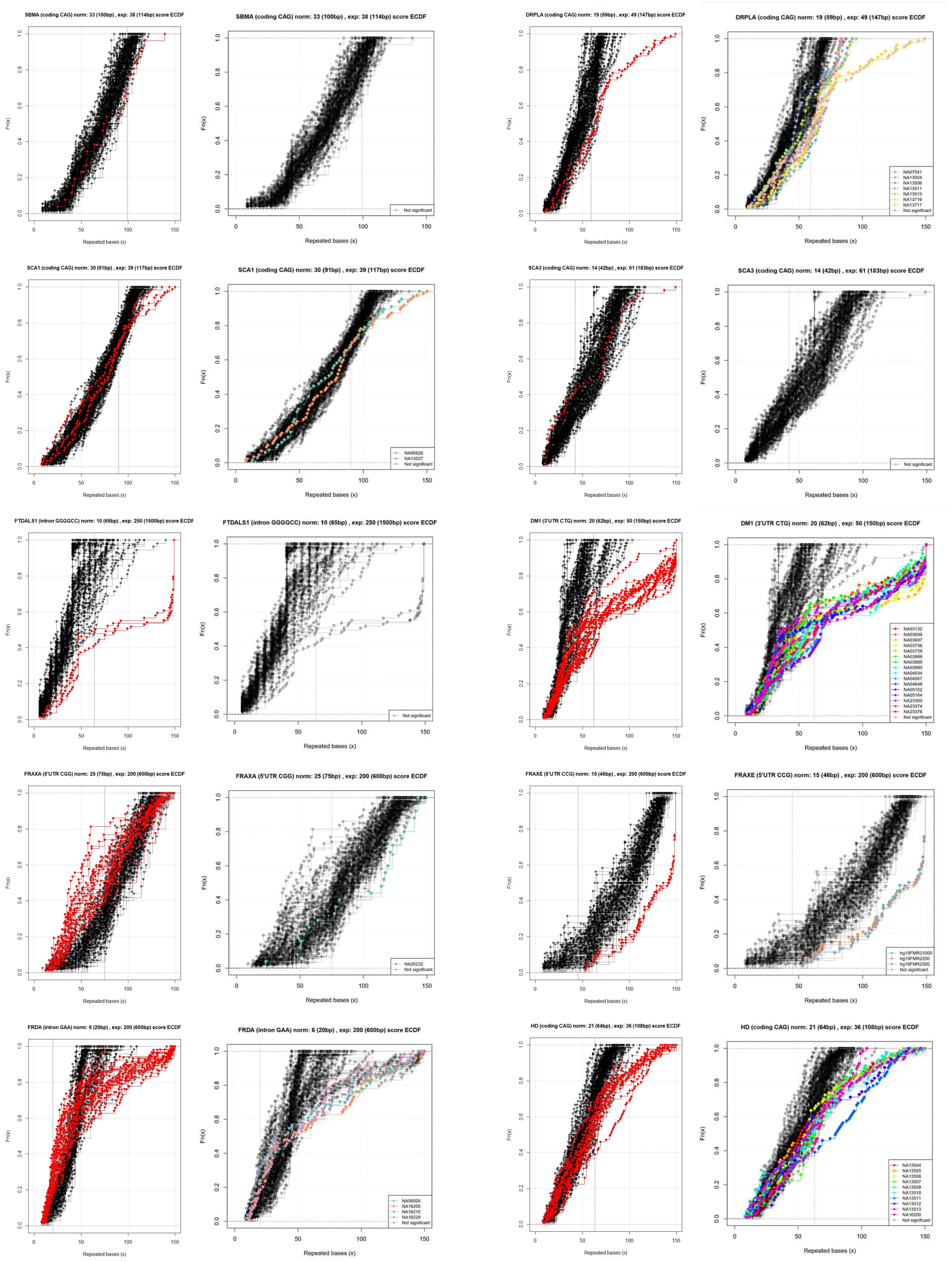

### *B. exSTRa ECDF and t-sum plots of BWA-aligned EGA and simulated genomes*

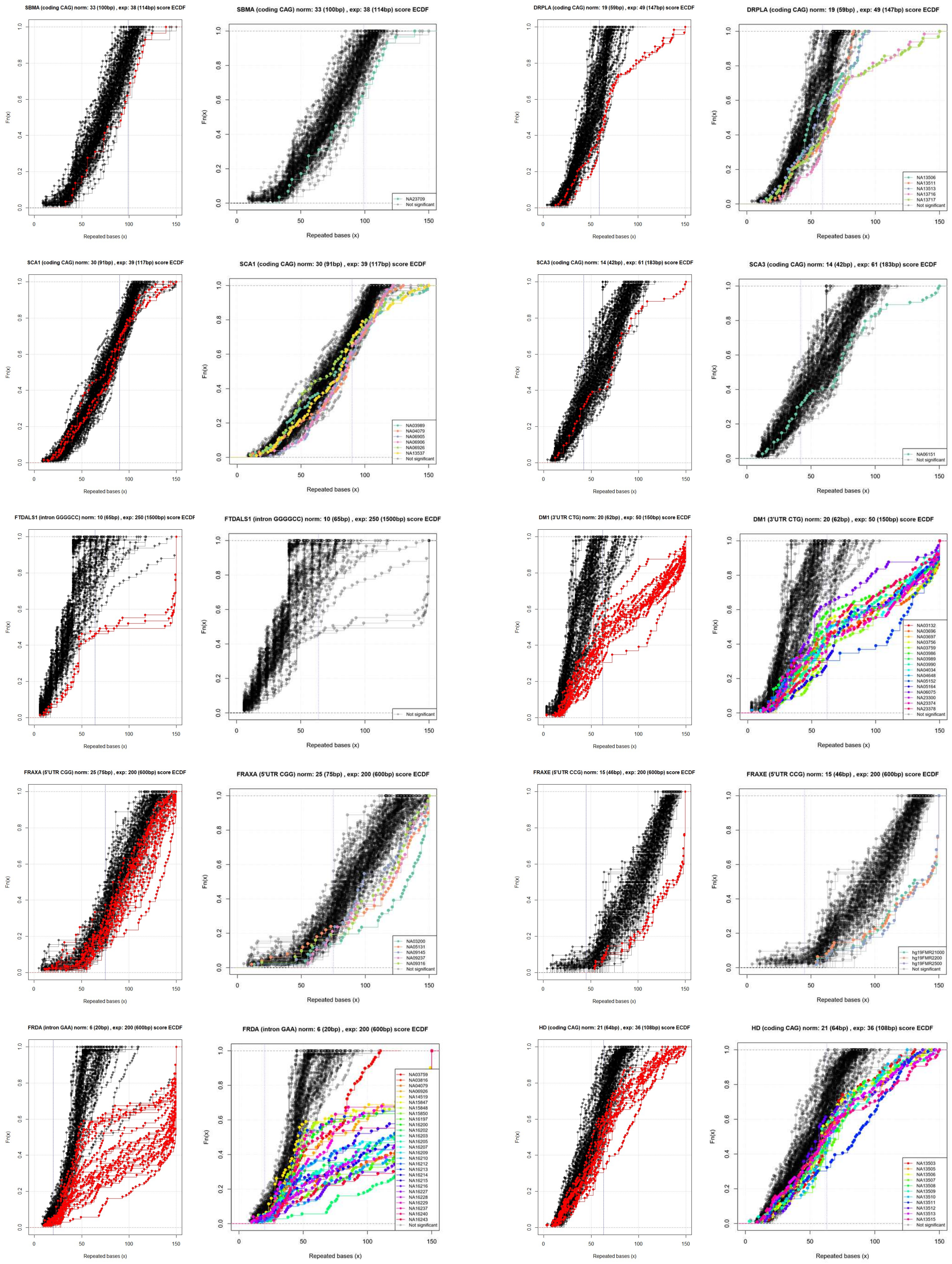

### *C. exSTRa ECDF and t-sum plots of BWA-aligned EGA and simulated genomes tested with 100 controls*

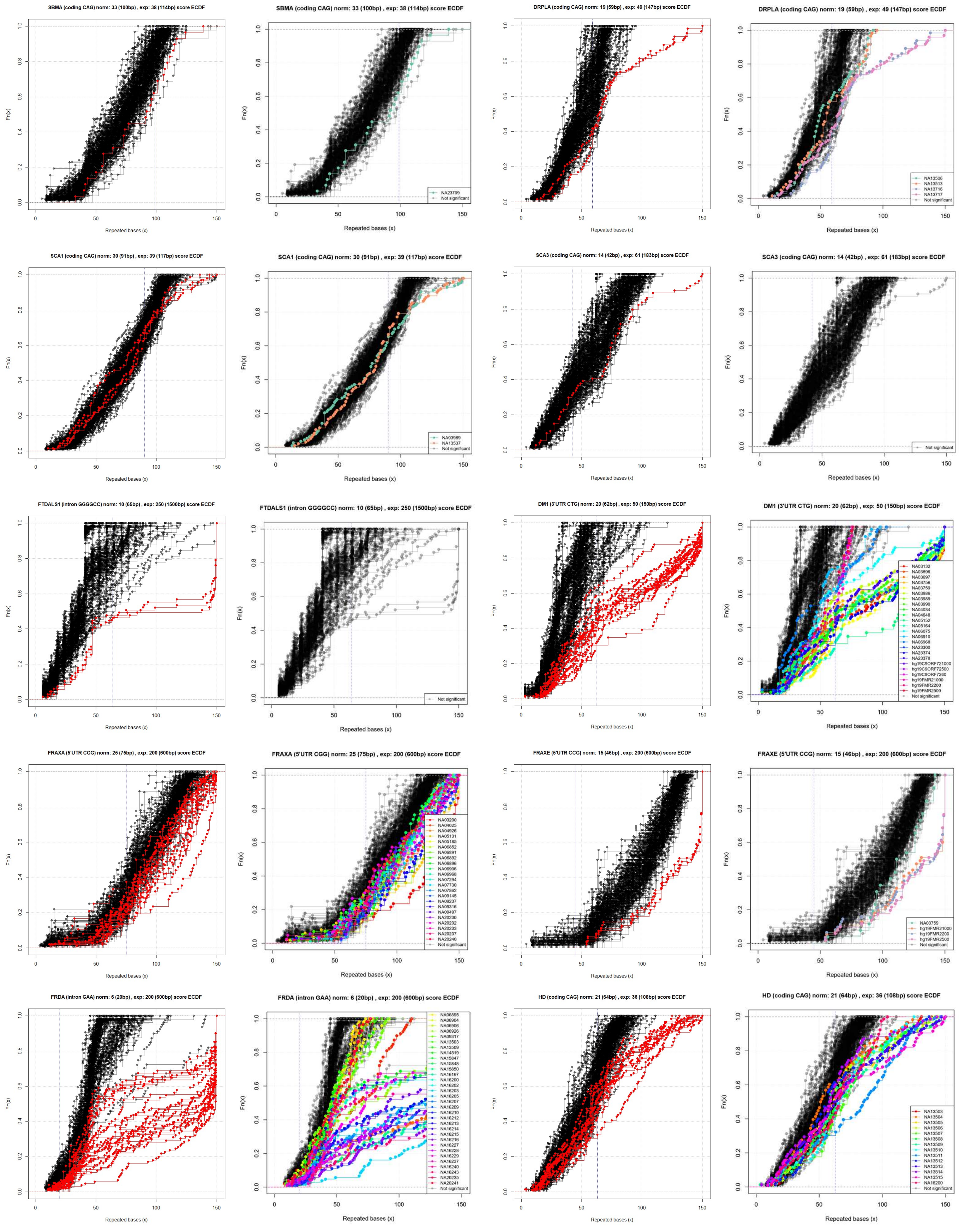

**Supplementary Figure 2.** Decision tree classification of the STR calls of the Isaac-aligned EGA and simulated genome sequence (GS) data by ExpansionHunter versions 2 and 3 (EH_v2 and EH_v3), GangSTR, TREDPARSE, STRetch, and exSTRa using default parameters. Panel (A) shows the decision tree generated by the classifier on the training dataset. Node #0 at the top of the tree is the root node. Each node lists an STR tool (feature). The “samples” number represents the total number of data points present within a particular node, and “value” shows the number of expanded (or full-mutation or FM) and non-expanded (non-FM) data points. The shade of the colour of each node reflects the proportion of expanded to non-expanded data points, with deeper blue and orange meaning more non-expanded and expanded data points, respectively. Gini index shows the impurity at each node. The terminal nodes shown in the last rows are the leaves. Leaves with a Gini of 0 have data points belonging to either the expanded or the non-expanded class. Panel B shows the ROC and precision-recall plots generated by the classifier on the test dataset. Panel C shows the ranking of the STR tools that contributed to the decision tree model.

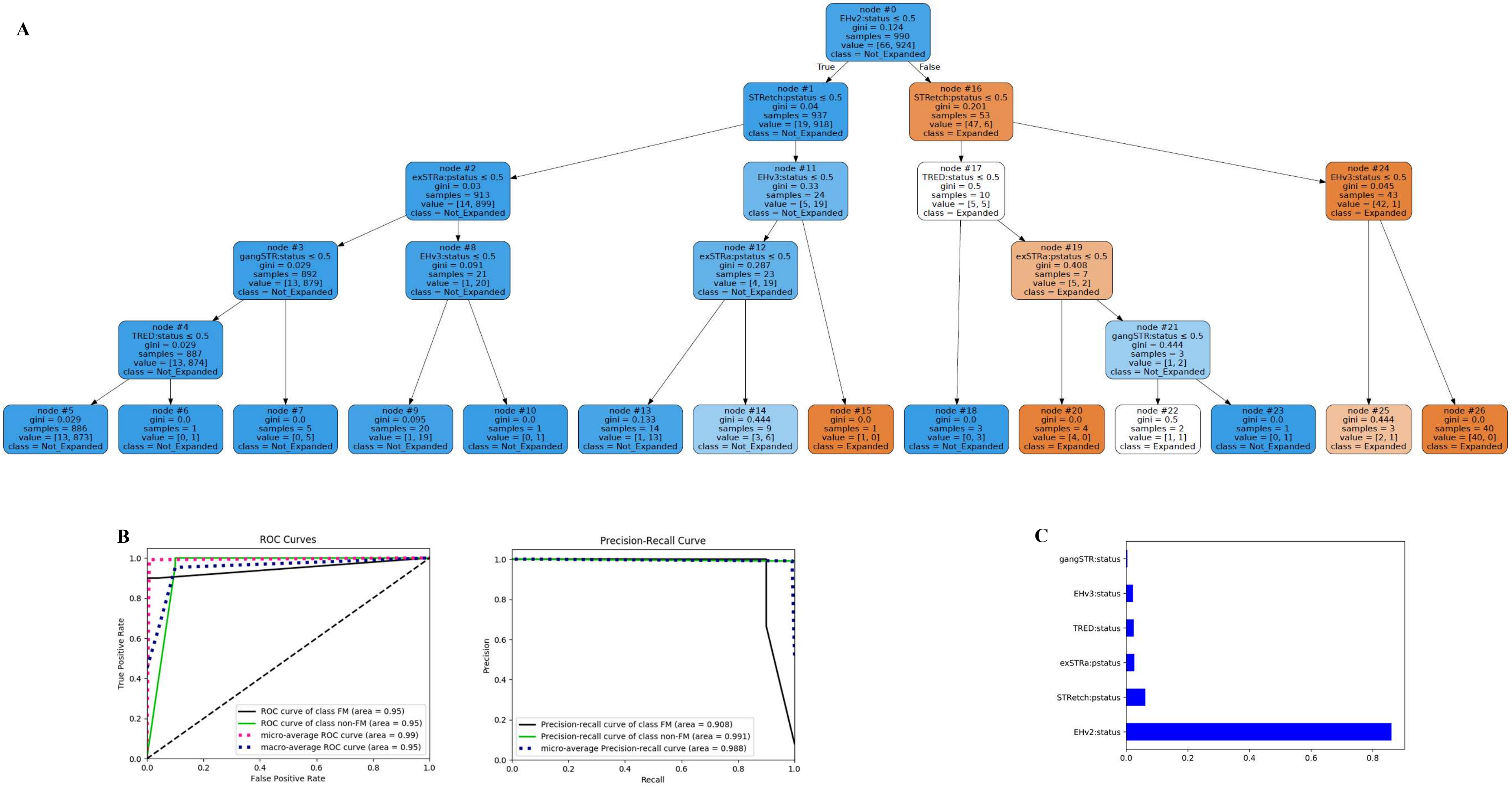
A
node #0
EHv2:status ≤ 0.5
gini = 0.124
samples = 990
value = [66, 924]
class = Not_Expanded
True
False
node #1
STRetch:pstatus ≤ 0.5
gini = 0.04
samples = 937
value = [19, 918]
class = Not_Expanded
node #16
STRetch:pstatus ≤ 0.5
gini = 0.201
samples = 53
value = [47, 6]
class = Expanded
node #2
exSTRa:pstatus ≤ 0.5
gini = 0.03
samples = 913
value = [14, 899]
class = Not_Expanded
node #11
EHv3:status ≤ 0.5
gini = 0.33
samples = 24
value = [5, 19]
class = Not_Expanded
node #17
TRED:status ≤ 0.5
gini = 0.5
samples = 10
value = [5, 5]
class = Expanded
node #24
EHv3:status ≤ 0.5
gini = 0.045
samples = 43
value = [42, 1]
class = Expanded
node #3
gangSTR:status ≤ 0.5
gini = 0.029
samples = 892
value = [13, 879]
class = Not_Expanded
node #8
EHv3:status ≤ 0.5
gini = 0.091
samples = 21
value = [1, 20]
class = Not_Expanded
node #12
exSTRa:pstatus ≤ 0.5
gini = 0.287
samples = 23
value = [4, 19]
class = Not_Expanded
node #19
exSTRa:pstatus ≤ 0.5
gini = 0.408
samples = 7
value = [5, 2]
class = Expanded
node #4
TRED:status ≤ 0.5
gini = 0.029
samples = 887
value = [13, 874]
class = Not_Expanded
node #21
gangSTR:status ≤ 0.5
gini = 0.444
samples = 3
value = [1, 2]
class = Not_Expanded
node #5
gini = 0.029
samples = 886
value = [13, 873]
class = Not_Expanded
node #6
gini = 0.0
samples = 1
value = [0, 1]
class = Not_Expanded
node #7
gini = 0.0
samples = 5
value = [0, 5]
class = Not_Expanded
node #9
gini = 0.095
samples = 20
value = [1, 19]
class = Not_Expanded
node #10
gini = 0.0
samples = 1
value = [0, 1]
class = Not_Expanded
node #13
gini = 0.133
samples = 14
value = [1, 13]
class = Not_Expanded
node #14
gini = 0.444
samples = 9
value = [3, 6]
class = Not_Expanded
node #15
gini = 0.0
samples = 1
value = [1, 0]
class = Expanded
node #18
gini = 0.0
samples = 3
value = [0, 3]
class = Not_Expanded
node #20
gini = 0.0
samples = 4
value = [4, 0]
class = Expanded
node #22
gini = 0.5
samples = 2
value = [1, 1]
class = Expanded
node #23
gini = 0.0
samples = 1
value = [0, 1]
class = Not_Expanded
node #25
gini = 0.444
samples = 3
value = [2, 1]
class = Expanded
node #26
gini = 0.0
samples = 40
value = [40, 0]
class = Expanded
B
ROC Curves
True Positive Rate
False Positive Rate
ROC curve of class FM (area = 0.95)
ROC curve of class non-FM (area = 0.95)
micro-average ROC curve (area = 0.99)
macro-average ROC curve (area = 0.95)
Precision-Recall Curve
Precision
Recall
Precision-recall curve of class FM (area = 0.908)
Precision-recall curve of class non-FM (area = 0.991)
micro-average Precision-recall curve (area = 0.988)
C
gangSTR:status
EHv3:status
TRED:status
exSTRa:pstatus
STRetch:pstatus
EHv2:status

**Supplementary Figure 3.** Decision tree classification of the STR calls of the BWA-aligned EGA and simulated GS data by ExpansionHunter versions 2 and 3 (EH_v2 and EH_v3), GangSTR, TREDPARSE, STRetch, and exSTRa using default parameters. The decision tree generated by the classifier on the training dataset (A), ROC and precision-recall plots generated by the classifier on the test dataset (B) and ranking of the STR tools that contributed to the decision tree model (C) are shown.

**A**

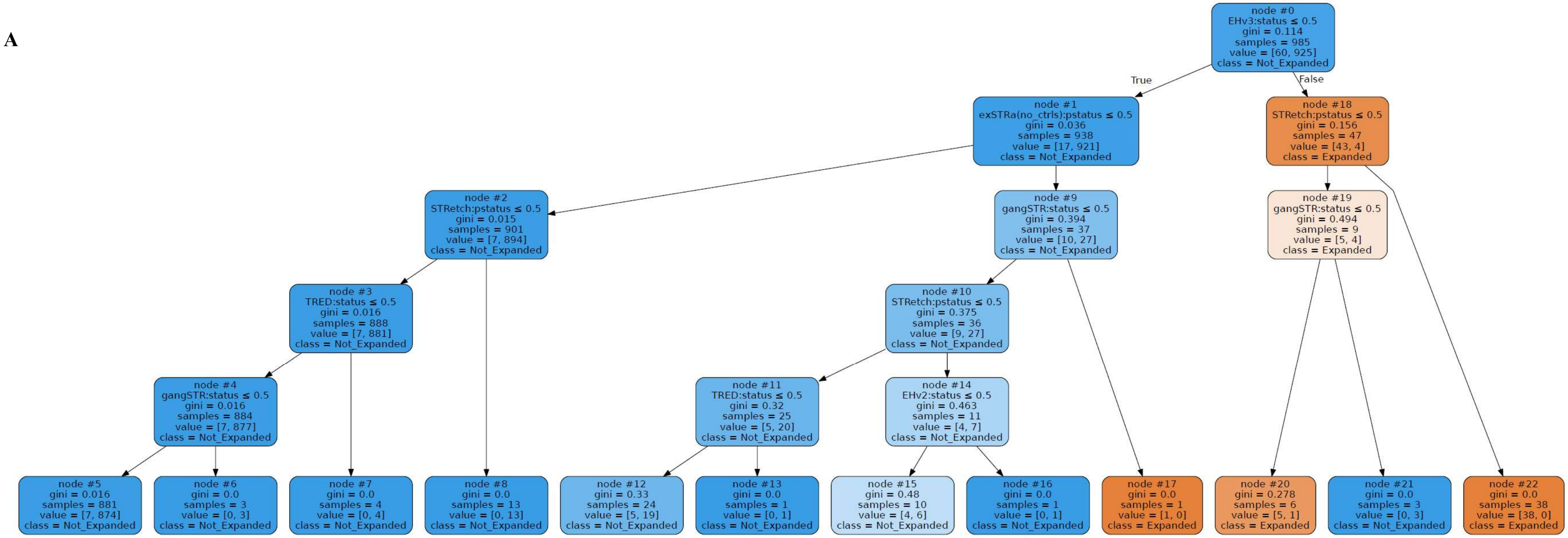


**B**

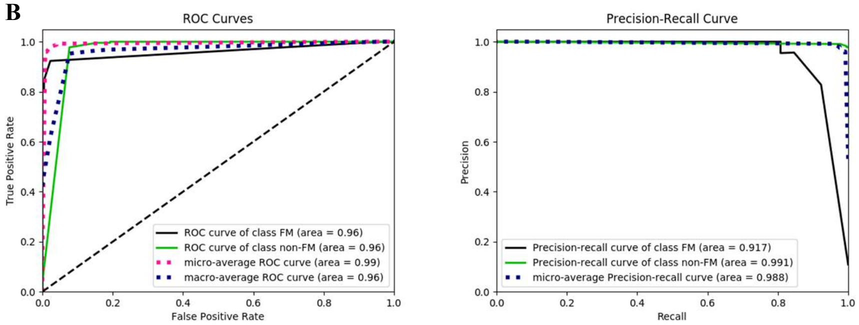


**C**

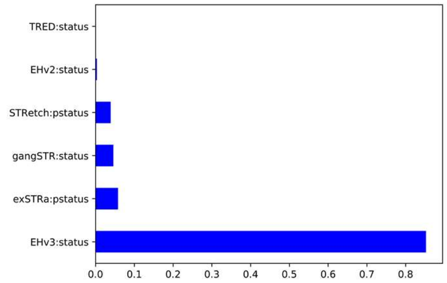

**Supplementary Figure 4.** Allelic distributions of repeat lengths for STR loci in *AR*, *ATN1*, *ATXN1*, *ATXN2*, *ATXN3*, *ATXN7*, *ATXN8/ATXN8OS*, *CACNA1A*, *CBL*, *CNBP*, *CSTB*, *DMPK*, *FMR1*, *FMR2*, *HTT*, *JPH3*, *NOP56*, *PHOX2B*, *PPP2R2B*, and *TBP* genes for CAUSES exomes as genotyped by ExpansionHunter (versions 2 and 3) and GangSTR. The 0 repeat alleles indicated in the plots for X chromosome STRs are that of hemizygous males, in whom the second allele was set to 0 and do not reflect the actual genotype calls of the tools.

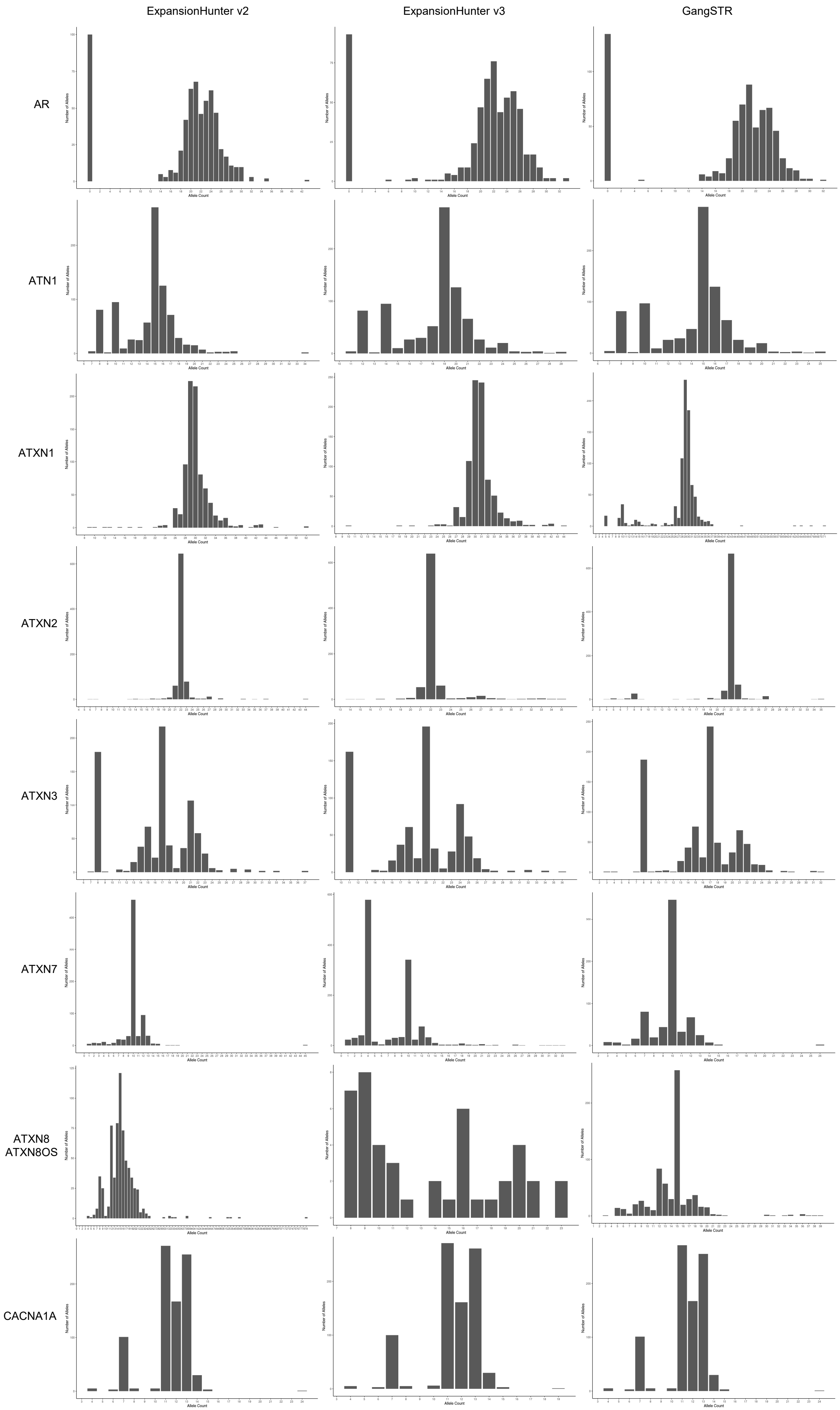
ExpansionHunter v2
ExpansionHunter v3
GangSTR
AR
ATN1
ATXN1
ATXN2
ATXN3
ATXN7
ATXN8
ATXN8OS
CACNA1A
Number of Alleles
Allele Count

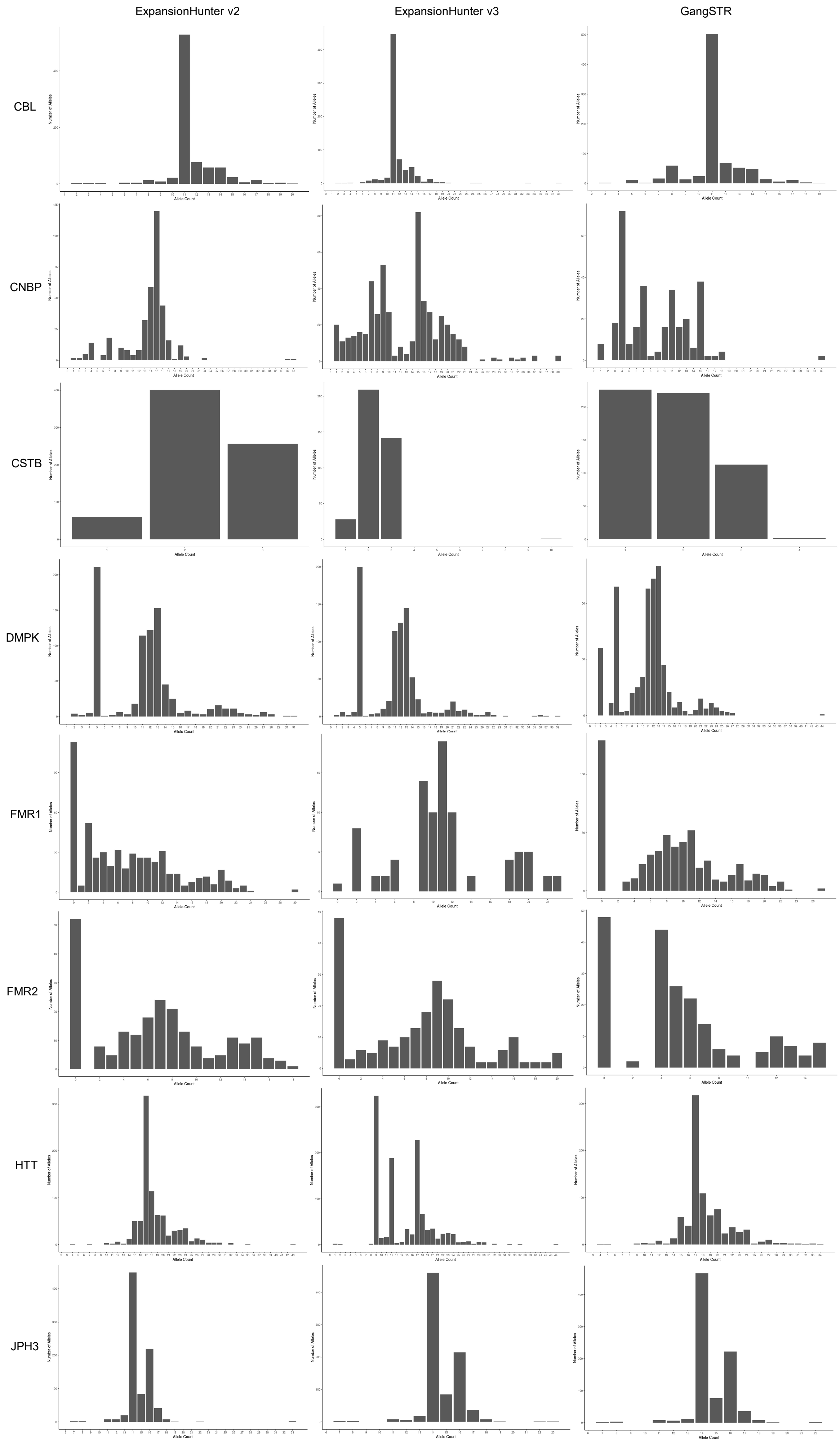
ExpansionHunter v2
ExpansionHunter v3
GangSTR
CBL
CNBP
CSTB
DMPK
FMR1
FMR2
HTT
JPH3
Number of Alleles
Allele Count

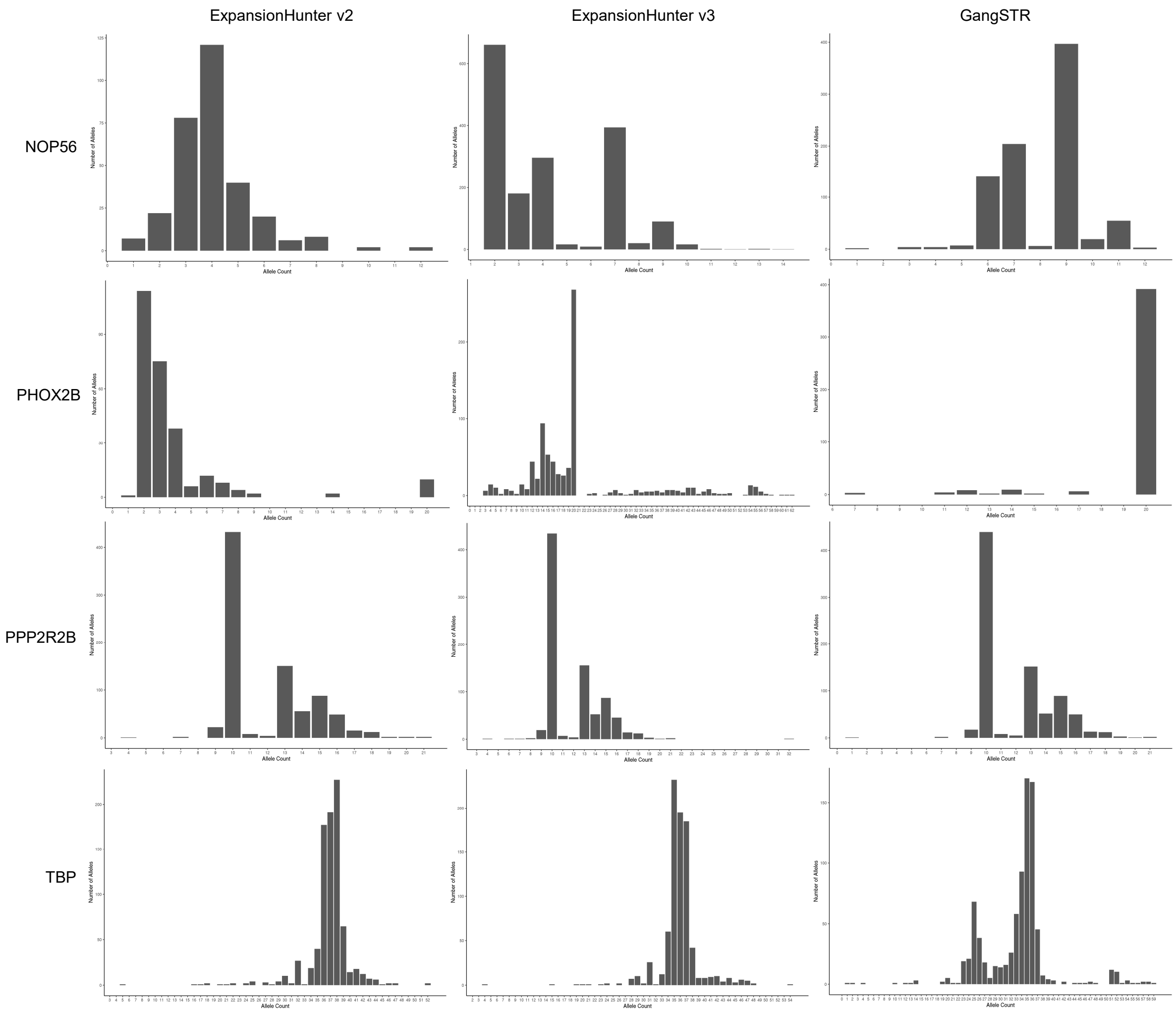
ExpansionHunter v2
ExpansionHunter v3
GangSTR
NOP56
PHOX2B
PPP2R2B
TBP
Number of Alleles
Allele Count

**Supplementary Figure 5.** Allelic distributions of repeat lengths for STR loci in *AR*, *ATN1*, *ATXN1*, *ATXN2*, *ATXN3*, *ATXN7*, *ATXN8/ATXN8OS*, *ATXN10*, *C9ORF72*, *CACNA1A*, *CBL*, *CNBP*, *CSTB*, *DMPK*, *FMR1*, *FMR2*, *FXN*, *HTT*, *JPH3*, *NOP56*, *PHOX2B*, *PPP2R2B*, and *TBP* genes for CAUSES and IMAGINE genomes as genotyped by ExpansionHunter (versions 2 and 3) and GangSTR. The 0 repeat alleles indicated in the plots for X chromosome STRs are that of hemizygous males, in whom the second allele was set to 0 and do not reflect the actual genotype calls of the tools.

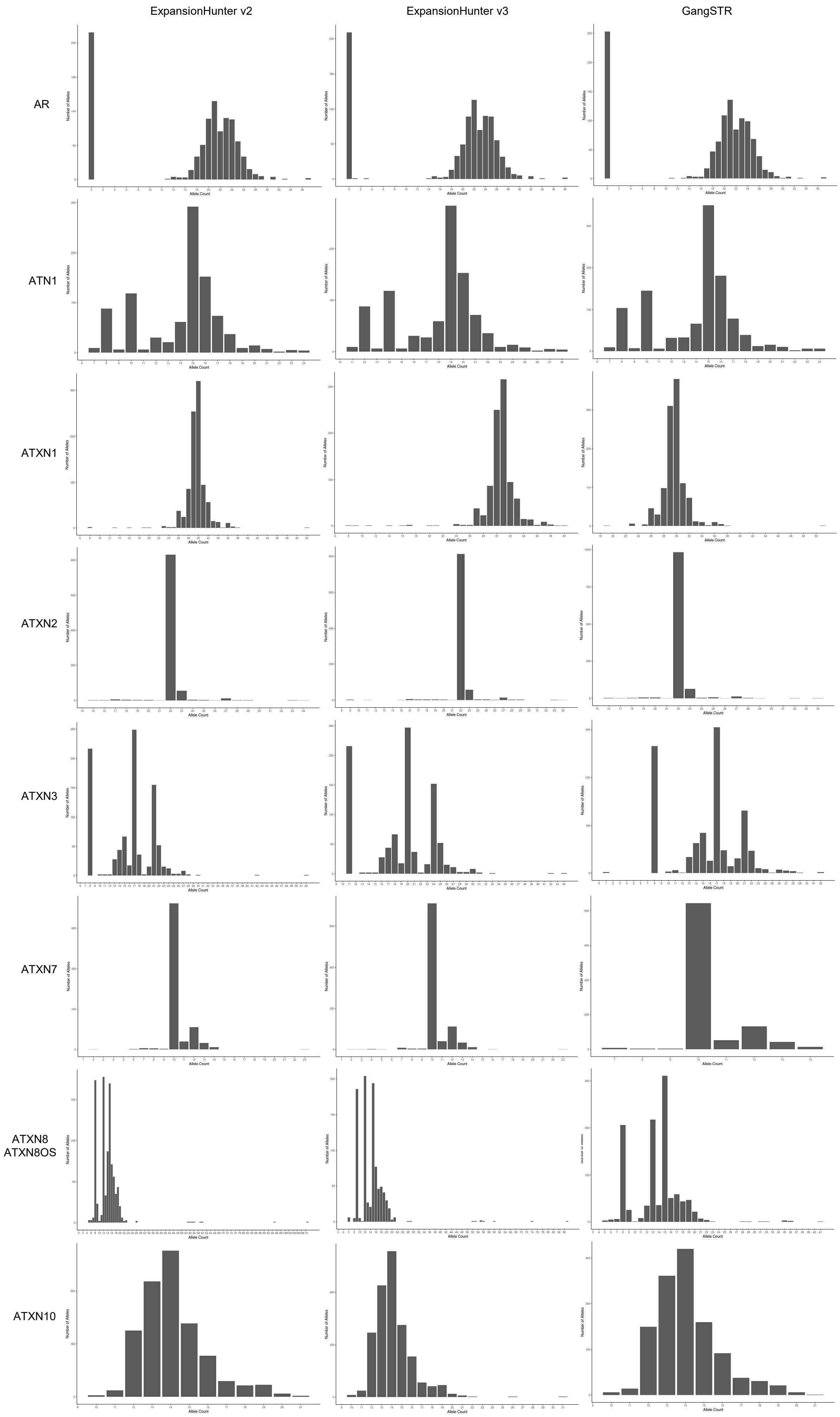
ExpansionHunter v2
ExpansionHunter v3
GangSTR
AR
ATN1
ATXN1
ATXN2
ATXN3
ATXN7
ATXN8
ATXN8OS
ATXN10
Number of Alleles
Allele Count

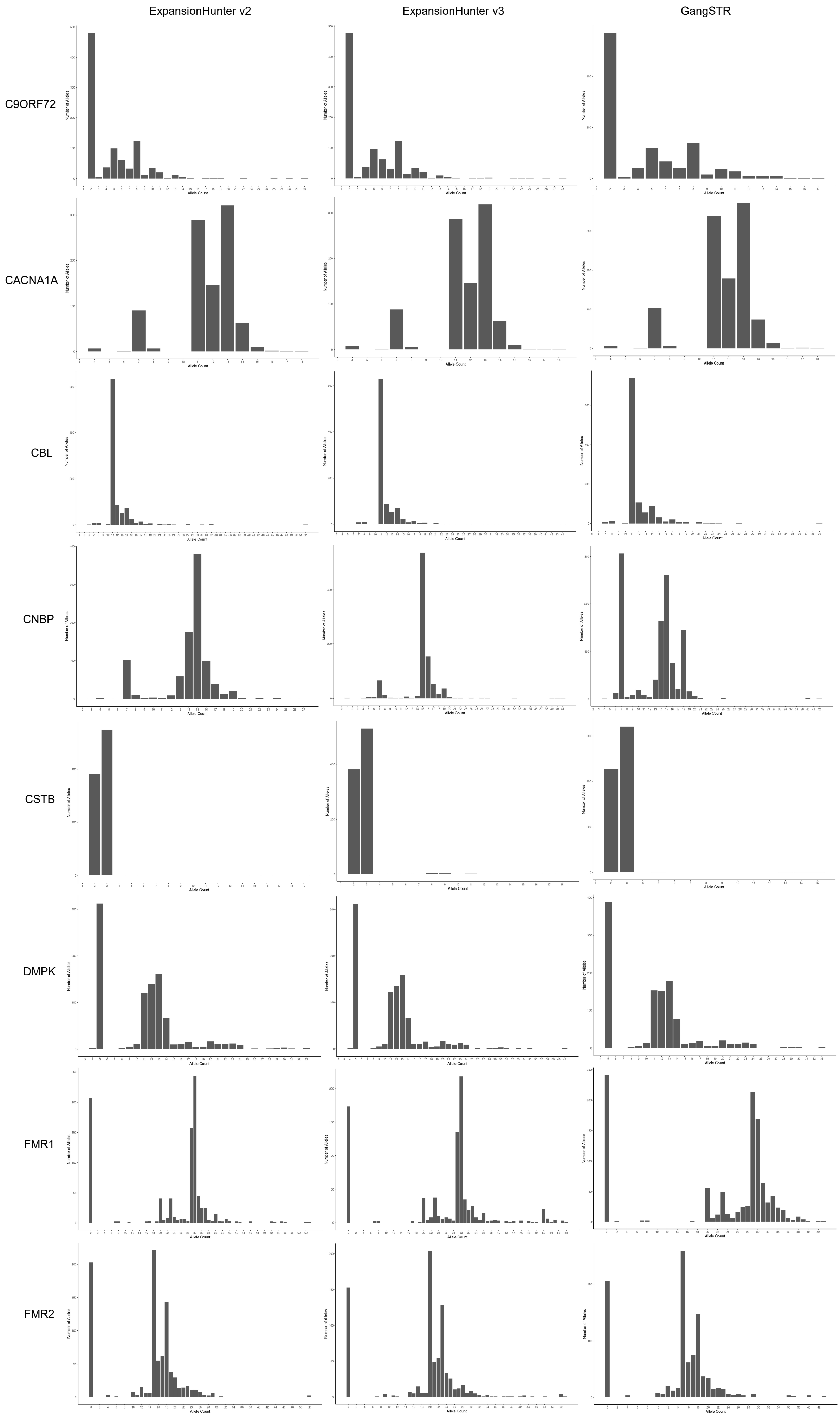
ExpansionHunter v2
ExpansionHunter v3
GangSTR
C9ORF72
CACNA1A
CBL
CNBP
CSTB
DMPK
FMR1
FMR2
Number of Alleles
Allele Count

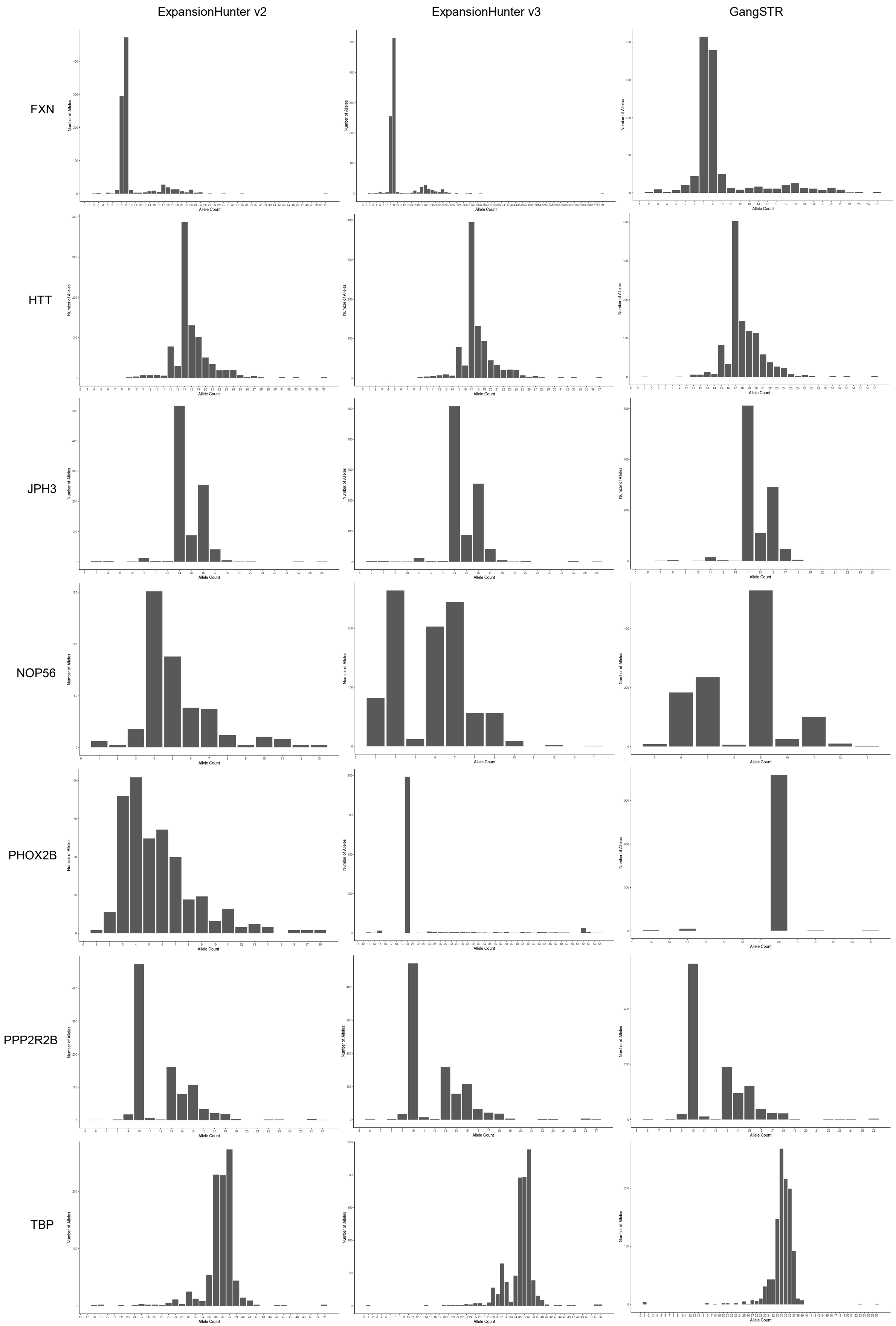
ExpansionHunter v2
ExpansionHunter v3
GangSTR
FXN
HTT
JPH3
NOP56
PHOX2B
PPP2R2B
TBP
Number of Alleles
Allele Count

**Supplementary Figure 6.** Analysis of the *DMPK* locus with and without off-target sites in Isaac- (A) and BWA-aligned (B) EGA GS data. The leftmost plots in Panels A and B show the estimated repeat lengths of the *DMPK* locus analysed in the EGA GS by ExpansionHunter version 2 with and without off-target sites alongside the consensus repeat size of known *DMPK* expansion-positive samples. The horizontal red-dotted and green-dotted lines define the lower- and upper-bound repeat lengths of *DMPK* full-mutation and normal alleles, respectively. The rightmost plots in Panels A and B show the Pearson's correlation between estimated and consensus repeat lengths of full-mutation alleles.

### Supplementary Figure 6

#### A. *DMPK Analysis in Isaac EGA GS*

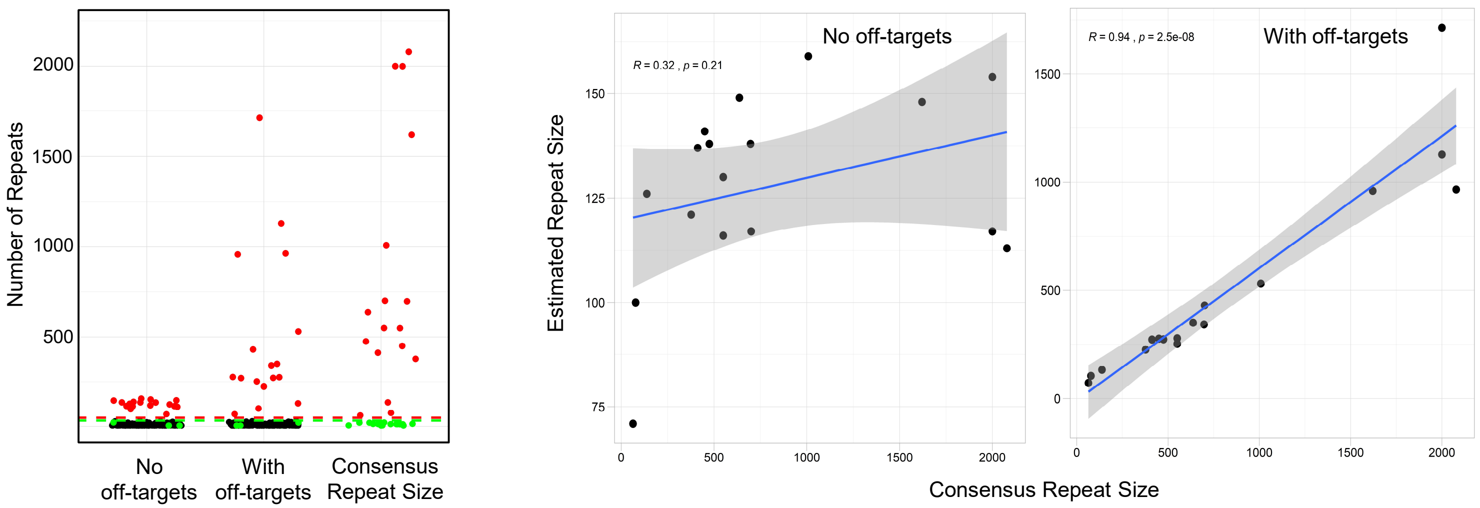


#### B. *DMPK Analysis in BWA EGA GS*

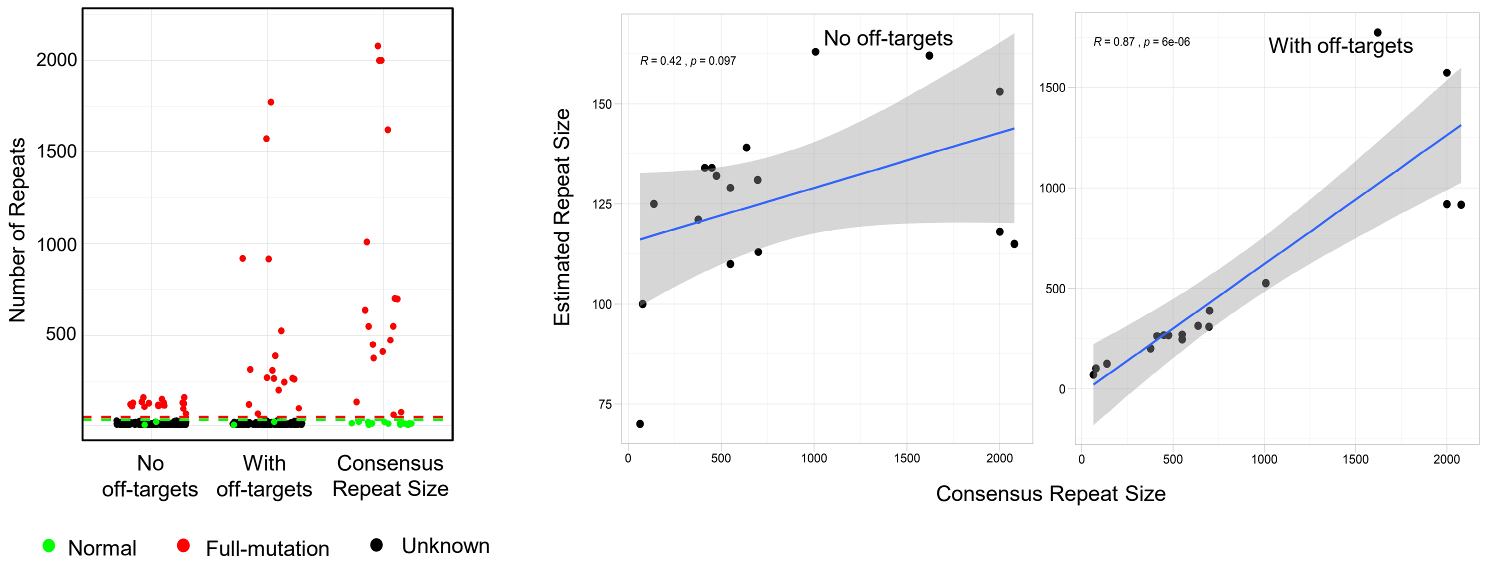
