## Supplementary Methods for "Genome-Wide Sequencing as a First-Tier Screening Test for Short Tandem Repeat Expansions"

### Coriell European Genome-phenome Archive Dataset True Repeat Size

The genotype status and repeat length information for most of the analyzed Coriell genomes have been previously characterized by PCR and/or SB analysis. The repeat length information for some of the NL and FM alleles were not available on the Coriell website (www.coriell.org). Therefore, we performed a literature search to retrieve this information from other studies that have determined repeat lengths in these cell lines using PCR and/or SB techniques. We used the Coriell data together with repeat lengths retrieved from other studies (cited in Supplementary Table 1a) to create a consensus repeat size catalog (last column of Supplementary Table 1a). We relied on Coriell's repeat length information for *AR*, *ATN1*, *ATXN1*, *ATXN3*, and *FXN* reference samples that lacked experimental data.

In general, PCR and capillary electrophoresis-based methods can determine the repeat lengths of most STR alleles with fewer than ~150 copies of a trinucleotide repeat, while SB is recommended to resolve hyperexpanded FM alleles such as those in the *DMPK*, *FXN*, and *FMR1* loci[1]. Therefore, for *HTT* reference samples with expansions that only ranged from 44 to 72 CAGs, we used data from a PCR study[2]. For *DMPK* and *FMR1* reference samples, we relied on PCR-based estimates for most alleles in the NL to PM size ranges and used available SB data for FM expansions. The repeat size data for some NL *DMPK* and *FXN* alleles and *FMR1* FM lengths of some fragile X samples could not be identified. In addition, we could not establish the consensus size estimate for two samples (NA06896 and NA20241) which appear to be "mosaic" and were reported to have very varied repeat lengths by different studies. For a few other cell lines that showed two distinct repeat lengths (e.g., NA06075 and NA09145) or had slight variation in reported repeat lengths (e.g., NA20233 and NA20240), we used their mean values as consensus size.

**ART WGS Data Simulation**

We simulated genomes using the ART next-generation sequencing read simulator[3]. Expanded alleles at STR loci in two genes were simulated from the hg19 human reference genome using the GATK FastaAlternateReferenceMaker tool (Supplementary Table 1b). Mutated genomes were simulated at 15x coverage with 150 bp paired-end reads to yield R1 and R2 fastq files. An unmutated reference genome was also simulated with the same parameters. The following ART command was used to simulate the genomes using the built-in quality profile for the Illumina HiSeq X PCR-free sequencing system:

```
art_illumina -ss HSXn -i $fasta_file -l 150 -f 15 -o $output_prefix -m 500 -s 50
```

The simulated fastq files from the unmutated reference genome were concatenated with the fastq file of the corresponding read for each mutated genome. This produced 30x mutated genomes generated as females with two X chromosomes and heterozygous STR alleles. These fastq files were then aligned to the hg19 human reference genome using BWA.
